# The SPN-4 Rbfox RNA-binding protein selects maternal mRNAs for CCR4-NOT-dependent clearance in early *Caenorhabditis elegans* embryos

**DOI:** 10.1101/2025.10.15.682621

**Authors:** Caroline A. Spike, Dylan M. Parker, Tatsuya Tsukamoto, Naly Torres-Mangual, Erika C. Tsukamoto, Karissa Coleman, Micah D. Gearhart, David Greenstein, Erin Osborne Nishimura

## Abstract

The elimination of maternal mRNAs is an essential feature of the maternal-to-zygotic transition. We report an essential pathway that clears many maternal transcripts from early *C. elegans* embryos using the Rbfox-related SPN-4 RNA-binding protein as a specificity factor and the CCR4-NOT deadenylase complex as an effector. We biochemically identified SPN-4-associated mRNAs from late-stage oocytes and found that the set of SPN-4-associated transcripts is enriched for maternal mRNAs that undergo early decay. Single-molecule fluorescence *in situ* hybridization experiments established that many SPN-4-associated mRNAs fail to be eliminated in the absence of SPN-4. In the 3’UTRs of two target mRNAs, we identified Rbfox motifs that are required for SPN-4-dependent clearance *in vivo* and bind SPN-4 *in vitro*. In a genetic screen to identify factors that work with SPN-4, we isolated mutant alleles of CCR4-NOT components. Auxin-induced degradation of the LET-711/NOT1 scaffold and the CCF-1 deadenylase disrupted clearance of two SPN-4-associated transcripts. Our results support a model in which SPN-4 initiates expression in late-stage oocytes, associates with maternal mRNA targets through RNA sequences in their 3’UTRs and promotes CCR4-NOT-mediated decay during early embryogenesis.

## INTRODUCTION

The transition from parental to embryonic control is an important aspect of reproduction. Because oocytes and early-stage embryos of most animals are transcriptionally quiescent, early embryonic development is orchestrated by parentally (predominantly maternally) encoded mRNAs and proteins. As genetic control is handed off from the maternal to the zygotic genome, mRNA clearance and zygotic genome activation (ZGA) work in tandem to sculpt the dynamic embryonic transcriptome (Tadros and Lipshitz, 2009; Vastenhouw et al., 2019; Yang et al., 2024). In several model organisms, an early phase of maternal mRNA clearance is driven by maternally provided factors, whereas a later phase of maternal mRNA clearance depends on zygotic gene products (Despic and Neugebauer, 2018). Together, these mRNA clearance mechanisms select transcripts for removal once their protein products have fulfilled their functions.

Maternal mRNA clearance is intimately coordinated with oogenesis, oocyte meiotic maturation, egg activation, fertilization and post-fertilization phases of meiosis and embryogenesis. In most animals, oocytes arrest in meiotic prophase I for a prolonged period. Hormonal signaling triggers the process of oocyte meiotic maturation in which oocytes resume meiosis and enter meiotic metaphase I (Huelgas-Morales and Greenstein, 2018; Mehlmann, 2005; Stetina and Orr-Weaver, 2011). The timing of the meiotic divisions with respect to fertilization varies between species, as does the timing of the onset of maternal mRNA clearance and its progression. In mice, early maternal mRNA clearance commences with meiotic maturation (Bachvarova et al., 1985; Paynton et al., 1988; Su et al., 2007); whereas in *Drosophila* it begins later, concurrent with egg activation, which happens after meiotic maturation when the oocyte enters the oviduct (Tadros and Lipshitz, 2009; Tadros et al., 2007). In this study, we investigate the mechanism of early maternal mRNA clearance in the nematode *C. elegans* and its links to oocyte meiotic maturation and early post-fertilization development.

In *C. elegans*, the onset of ZGA occurs at the 4-cell stage and progressively continues from the 4-to 32-cell stages (Baugh et al., 2003; Edgar et al., 1994; Hashimshony et al., 2015; Seydoux and Fire, 1994; Seydoux et al., 1996; Tintori et al., 2016). Although several early studies had observed maternal mRNA clearance (Baugh et al., 2003; Seydoux and Fire, 1994; Seydoux et al., 1996; Tintori et al., 2016), these reports focused on the elimination of maternal transcripts occurring at and after the onset of ZGA, which likely corresponds to the later phases of transcript turnover. One landmark study characterized the extent of maternal mRNA clearance within the first two cell divisions (Stoeckius et al., 2014), prior to the onset of ZGA. Stoeckius et al. (2014) compared the transcriptomes of 1-cell and 2-cell embryos to purified oocytes, thereby capturing instances of early mRNA clearance. This showed that approximately 25% of oocyte transcripts degrade during early embryogenesis. Noticing that a subset of these decaying maternal mRNA transcripts (∼10%) were complementary to endogenous small-interfering RNAs (siRNA), Stoeckius et al. (2014) proposed a role for siRNAs in early maternal mRNA clearance. Indeed, a later study provided evidence that siRNAs and the Argonaute CSR-1 do participate in maternal mRNA clearance, albeit at a later stage, occurring gradually from the 4-20-cell stage and onwards (Quarato et al., 2021).

One transcript that undergoes dramatic early maternal mRNA clearance is *lin-41* (Stoeckius et al., 2014). *lin-41* encodes the tripartite motif (TRIM)-NHL (NCL-1, HT2A, and LIN-41) RNA-binding protein that promotes meiotic prophase I arrest, an essential step in oogenesis (Spike et al., 2014a; Spike et al., 2014b; Tocchini et al., 2014). Loss of *lin-41* results in premature entry into meiotic metaphase in which pachytene-stage oocytes prematurely cellularize, disassemble the synaptonemal complex and activate the CDK-1 cyclin-dependent kinase precociously, leading to sterility (Spike et al., 2014a). Premature CDK-1 activation in *lin-41* null mutant oocytes causes them to transcribe genes that are normally expressed at or after ZGA (Allen et al., 2014; Spike et al., 2014a; Tocchini et al., 2014). LIN-41 collaborates with and antagonizes OMA-1 and OMA-2, two tristetraprolin/TIS11-related RNA-binding proteins (collectively referred to as the OMA proteins) (Spike et al., 2014a; Spike et al., 2014b). OMA proteins counter LIN-41 activity by promoting entry to meiotic metaphase I thereby triggering meiotic maturation. By regulating the translational efficiencies of their maternal mRNA targets, these two RNA-binding proteins dynamically shape the proteome to ensure proper oogenesis and early post-fertilization development.

LIN-41 and the OMA proteins mediate a translational repression-to-activation switch in which LIN-41 represses, and the OMA proteins activate, translation of the Rbfox RNA-binding protein SPN-4 (Tsukamoto et al., 2017). Late in oogenesis, LIN-41 is inactivated as a translational repressor and subjected to ubiquitin-mediated protein degradation (Spike et al., 2018), enabling the translation of SPN-4 concurrent with the onset of oocyte meiotic maturation oocyte meiotic maturation (Fig. 1). SPN-4 translation continues into early embryogenesis when SPN-4, a divergent member of the Rbfox family (Kuroyanagi, 2009), has an essential role (Gomes et al., 2001). Prior results using RNA interference (RNAi) and a hypomorphic allele suggested a potential role for SPN-4 in promoting the degradation of maternal transcripts in the early embryo (Parker et al., 2020).

**Fig. 1.**
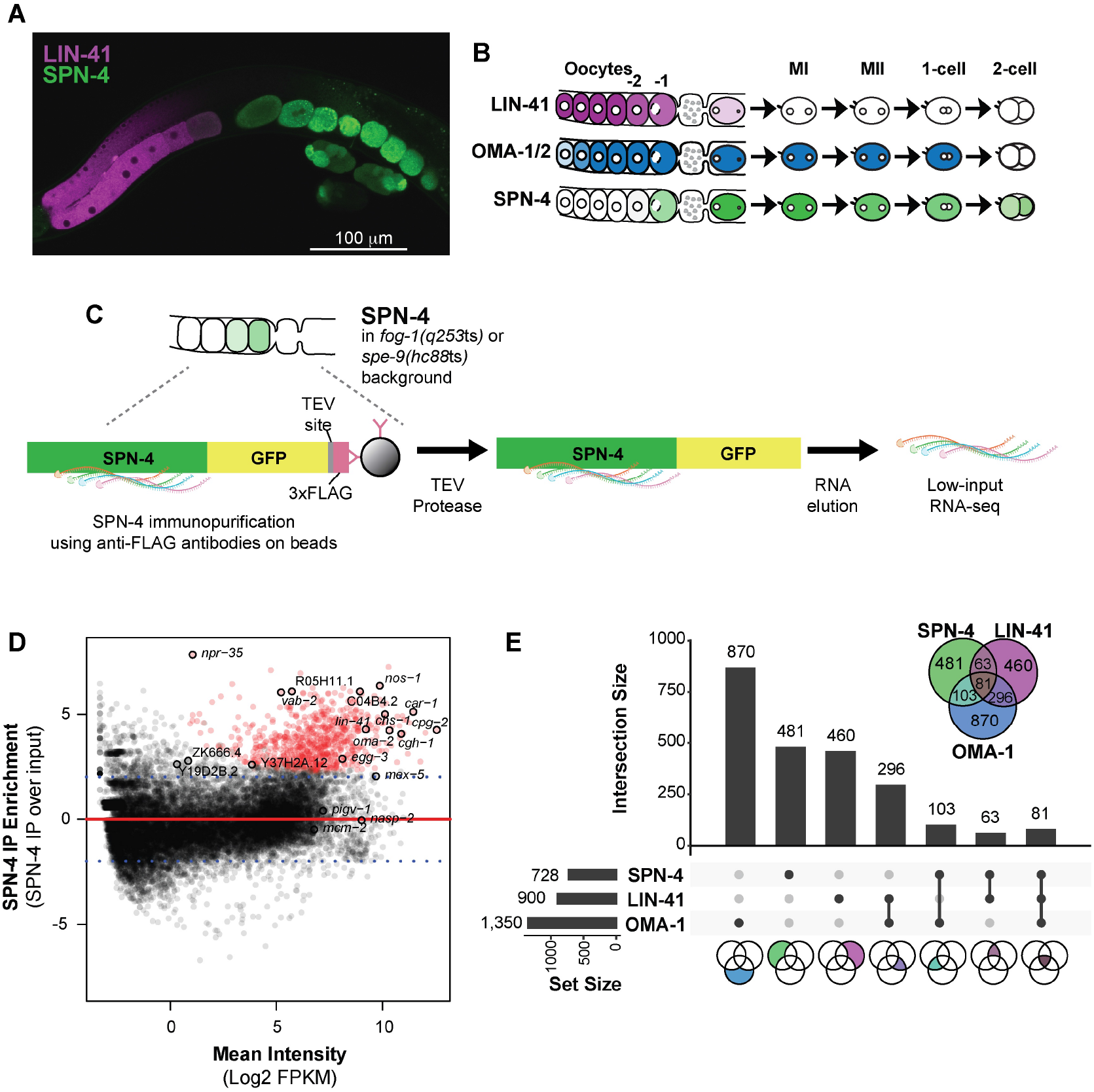
SPN-4 associates with mRNAs in oocytes. (A) LIN-41(magenta) and SPN-4 (green) expression patterns are largely mutually exclusive. (B) Depictions of LIN-41, OMA-1/2 and SPN-4 in oocytes and early embryos. (C) Purification of SPN-4 and its associated mRNAs from strains in which SPN-4 was endogenously tagged with GFP (yellow), a TEV cleavage site (grey) and 3xFLAG tags (pink). Immunopurifications were conducted in *fog-1(q253ts)* or *spe-9(hc88ts)* fertilization incompetent mutant backgrounds. (D) Enrichment of transcripts in SPN-4 immunopurifications (over input) plotted over mRNA abundance. Statistically significant SPN-4-associated transcripts (red) and key transcripts (labeled) are shown. (E) LIN-41- and OMA-1-associated transcripts were analyzed using the same protocol, and intersections between the sets of SPN-4-, LIN-41- and OMA-1-associated transcripts (4-fold enrichment with *P*>0.05) are shown via a Venn Diagram and an Upset Plot. Data from five to six replicates.

Here, we provide evidence that SPN-4 functions as an essential regulator of early maternal mRNA clearance. We present a model in which SPN-4 associates with maternal mRNA targets upon the onset of oocyte meiotic maturation and directs their CCR4-NOT-dependent clearance during early embryogenesis. We estimate that 10.77% of maternal transcripts subject to early decay are dependent on SPN-4 for their elimination. We suggest that the pleiotropic and early nature of strong loss-of-function *spn-4* alleles is caused by perturbation of the early embryonic transcriptome.

## RESULTS

### Identification of SPN-4-associated mRNAs

To understand how SPN-4 impacts early embryogenesis, we sought to identify SPN-4-associated mRNAs. One challenge is that SPN-4 may stimulate the degradation of its targets. For example, SPN-4 directly binds *nos-2* 3’UTR sequences, represses *nos-2* 3’UTR reporters in early embryos (Jadhav et al., 2008), and promotes *nos-2* mRNA decay (Parker et al., 2020). To address this issue, we immunopurified SPN-4 from the oocytes of sterile adult hermaphrodites that do not produce embryos, either because the major sperm protein signal for oocyte meiotic maturation is absent *(fog-1(q253ts)* genetic background) or because fertilization does not occur *((spe-9 (hc88ts)* genetic background).

The cohorts of mRNA that associate with immunopurified SPN-4 were identified using a low-input RNA-sequencing (RNA-seq) method (Fig. 1C). We previously characterized LIN-41- and OMA-1-associated mRNAs using a similar but less sensitive approach (Tsukamoto et al., 2017). LIN-41- and OMA-1-associated mRNAs were reassessed using the same low-input RNA-seq method, also from sterile oocytes, so that we could compare SPN-4, LIN-41 and OMA-1 mRNA targets. The two methods of library preparation for RNA-seq yielded high correlations for LIN-41 and OMA-1 (r^2^ = 0.76 and r^2^ = 0.82, respectively; Fig. S1), demonstrating that the low-input RNA-seq method generates similar transcriptome profiles while accommodating the lower abundance of SPN-4 in the germline (Fig. 1A,B).

We identified 728 mRNAs exhibiting a statistically significant four-fold or greater enrichment in SPN-4 purifications compared to input RNA (*P*<0.05, Benjamini-Hochberg adjusted *P*-value; Fig. 1D, Table S1). Using similar metrics, we identified 900 mRNAs that associate with LIN-41 and 1,350 mRNAs that associate with OMA-1. The overlap between mRNA cohorts was greatest between LIN-41 and OMA-1 (Fig. 1E, Fig. S2A,B). Of the LIN-41-associated mRNAs, 33% were shared with OMA-1, supporting previous findings that OMA-1 and LIN-41 reside in shared complexes and target overlapping sets of mRNAs (Tsukamoto et al., 2017). In contrast, when comparing SPN-4-associated mRNAs to either LIN-41- or OMA-1-associated sets, only 4-14% of mRNA sets overlapped.

We performed Gene Ontology (GO) analysis to determine whether the three RNA-binding proteins associate with mRNAs of either similar or distinct functions (Fig. S2C). SPN-4-associated mRNAs, independent of OMA-1 or LIN-41 association, were enriched for categories “eggshell formation,” variations on “sexual reproduction,” “vesicle-mediated transport” and “cell cycle process.” The “cell cycle process” category is noteworthy as loss or depletion of *spn-4* leads to early pleiotropic defects in spindle orientation and embryonic cell divisions (Gomes et al., 2001). Some defects observed in *spn-4* mutants could therefore be caused by an altered dosage of proteins encoded by SPN-4-associated mRNAs. The “eggshell formation” category is also noteworthy; the eggshell and its constituent layers are essential for early embryogenesis. These proteins are often degraded once the eggshell has formed (Maruyama et al., 2007) suggesting that both the transcript and its encoded protein are destroyed in tandem as part of a coordinated developmental transition.

### SPN-4-associated transcripts include many maternal mRNAs that turn over in the early embryo

RNAi knockdown of *spn-4* leads to an overabundance of *chs-1, cpg-2* and *nos-2* transcripts in early embryos (Parker et al., 2020). We identified *chs-1, cpg-2* and *nos-2* as SPN-4-associated transcripts (Table S1), leading us to examine whether the larger set of 728 SPN-4-associated mRNAs exhibits evidence of early embryonic clearance. Single-cell resolution RNA-seq (scRNA-seq) data from the early embryo (Tintori et al., 2016) tracks mRNA abundance in the 31 blastomeres encompassing the 1-to 16-cell stages. We filtered this dataset based on SPN-4, LIN-41 or OMA-1 associations and plotted the mean abundance of each cohort organized by cell and time. SPN-4-associated mRNAs as a class showed decreasing abundance in somatic cells over developmental time (Fig. 2). In contrast, mRNAs that associate with LIN-41 and OMA-1 did not exhibit this trend or did so to a lesser degree (Fig. 2). The germline P-lineage retains a zygote-like transcriptome during these stages owing in part to its prolonged transcriptional quiescence (Guven-Ozkan et al., 2008; Seydoux and Dunn, 1997; Seydoux et al., 1996). To further examine the relationship between SPN-4 association and maternal mRNA clearance, we clustered transcripts by RNA-expression class (Fig. S3A,B). Cluster Sets 3 and 5 represent maternal mRNAs that undergo decay. When expression level is considered, SPN-4-associated mRNAs account for 9.74% of Cluster Set 3 and 13.14% of Cluster Set 5, suggesting that direct SPN-4 action could account for the clearance of up to 10.77% of the maternal dowry (when the two clusters were calculated together) (Fig. S3C). We used mosaic plot analysis to determine whether the correlation between SPN-4 association and maternal mRNA clearance was statistically significant. All RNA-binding protein-associated cohorts were over-represented in Cluster Set 3, and SPN-4-associated cohorts were over-represented in Cluster Set 5 (Fig. S3D). These analyses are consistent with the idea that SPN-4-associated mRNAs are likely to undergo early mRNA clearance.

**Fig. 2.**
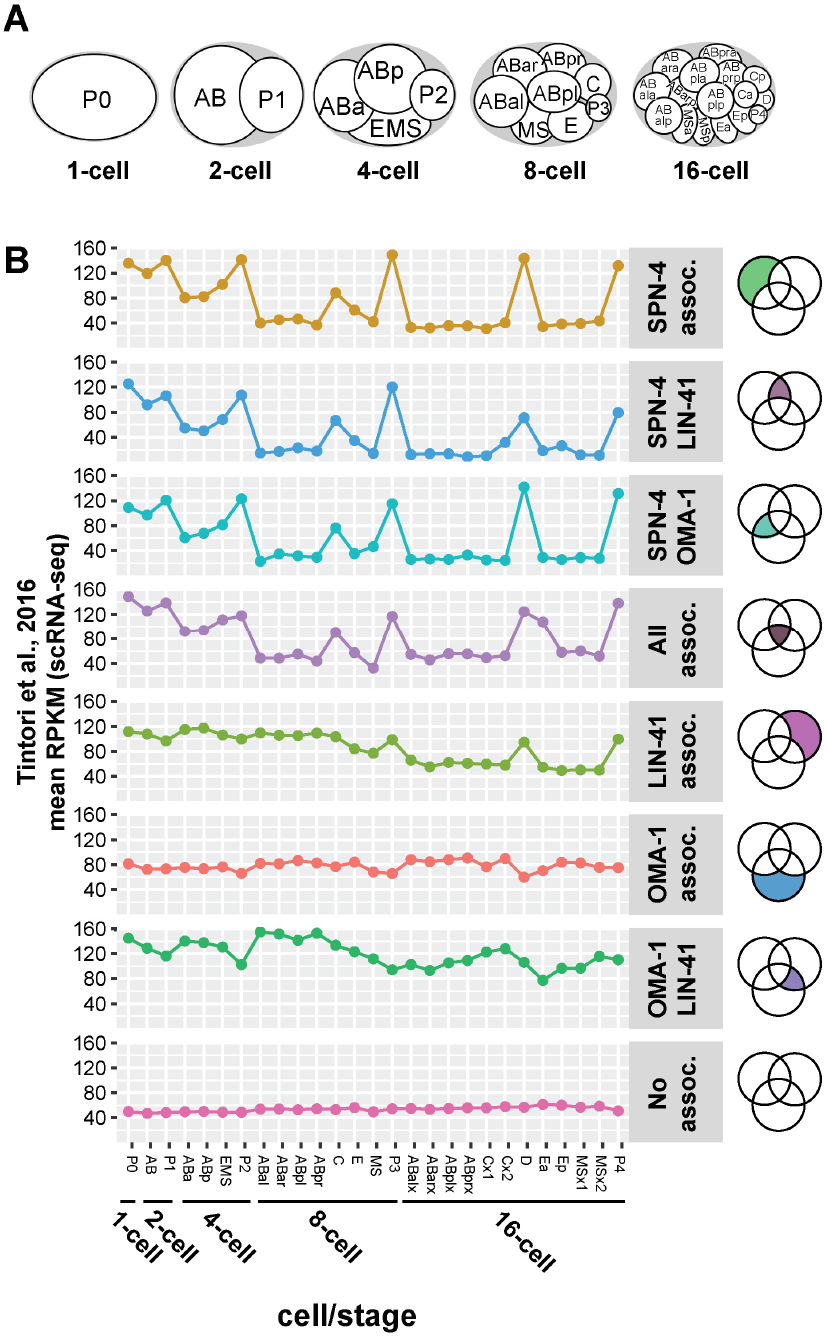
SPN-4-associated mRNAs undergo early embryonic decay. (A) Cartoon depiction of the first five cell divisions of *C. elegans* embryonic development. (B) Plots of the mean abundance of different transcript classes over early development, grouped according to their association with SPN-4, LIN-41 and OMA-1. The mean transcript abundance in reads per kilobase of transcript per million mapped reads (RPKM) is taken from a scRNA-seq dataset generated from hand-dissected, early-stage blastomeres (Tintori et al., 2016).

### SPN-4 is required for maternal mRNA clearance of associated transcripts

To determine whether SPN-4 is required for early mRNA clearance of its associated transcripts, we assessed the effect of *spn-4* on transcript abundance using single molecule fluorescence *in situ* hybridization (smFISH) or an inexpensive alternative (smiFISH). To compare embryos with and without maternal *spn-4* activity, we edited the endogenous *spn-4* locus to create a *tmC3* balancer chromosome that expresses SPN-4::GFP (Fig. 3A). Large populations of embryos were isolated from a strain containing this balancer chromosome and the *spn-4 (tm291)* deletion allele. Embryos with maternal *spn-4* activity express SPN-4::GFP and are easily differentiated from embryos produced by *spn-4* null mutant parents, which lack GFP fluorescence (Fig. 3B). Adult hermaphrodites with a *spn-4::gfp* genotype exhibit brood sizes that are no different than the wild type (Fig. S4A). However, as we will show, *spn-4::gfp* behaves as a hypomorph in smFISH assays (Fig. S4B) and sensitized genetic tests.

**Fig. 3.**
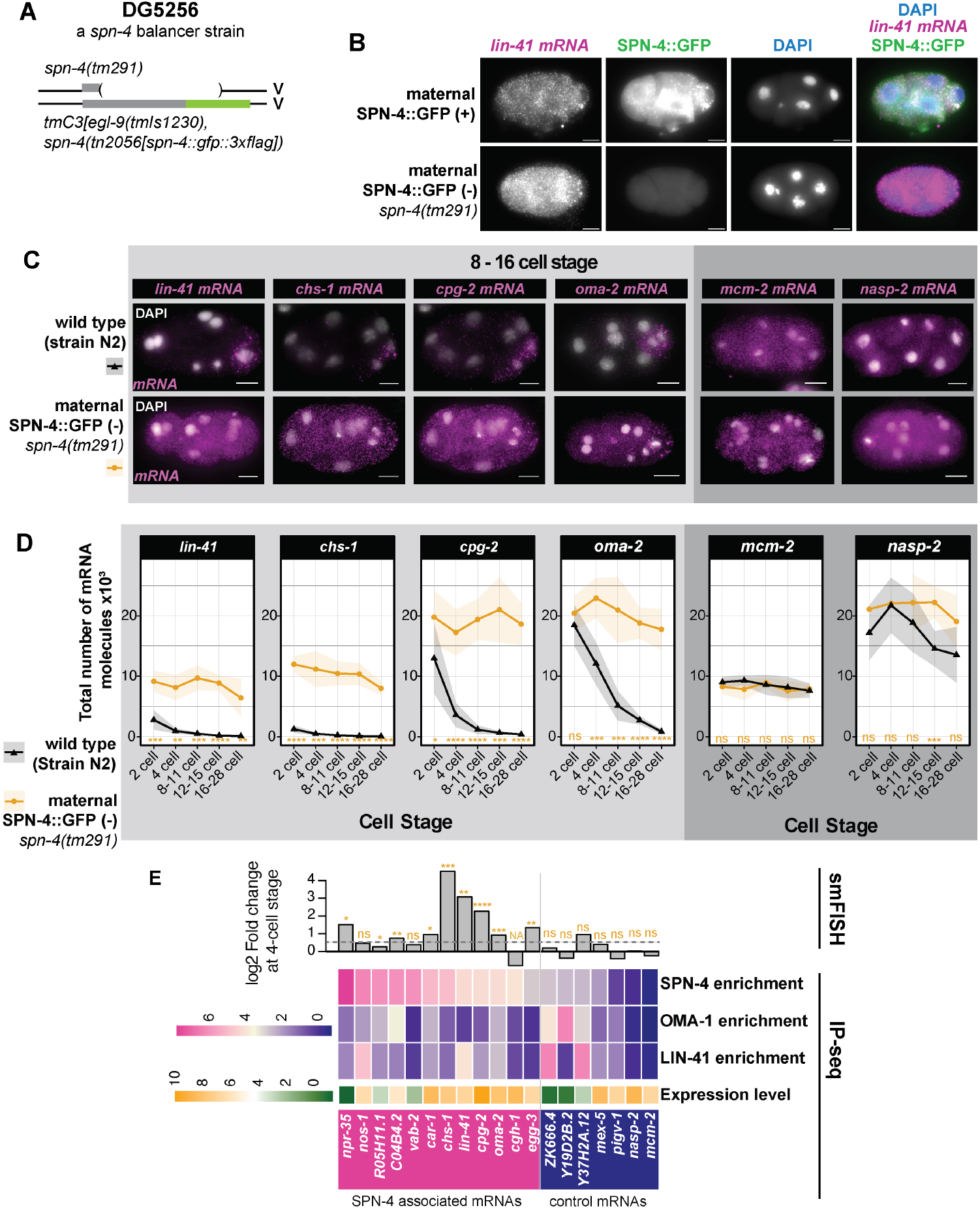
SPN-4 is required for clearance of SPN-4-associated mRNAs. (A) The *spn-4* balancer strain used to generate embryos with and without maternal *spn-4* activity. (B) Embryos produced by *spn-4::gfp* heterozygous and homozygous mothers exhibited GFP fluorescence (top panel) whereas embryos produced by *spn-4(tm291)* homozygous null mutants lacked fluorescence and yielded elevated *lin-41* mRNA levels (bottom panel). (C,D) The abundance of key transcripts in the wild type (strain N2) or without maternal *spn-4* activity as imaged by smiFISH (C) and quantified over five developmental stages (D). Four SPN-4-associated transcripts and two controls are illustrated. Bars, 10 µm. Typically, 7 embryos (with a range of 4-12) per transcript, genotype and stage combination were collected across two to three replicates. Ribbons, standard deviation. Statistics = Welch’s *t* -tests adjusted by Benjamini-Hochberg multiple test correction. ****: *P*≤0.0001, ***: *P*≤0.001, **: *P*≤0.01, *: *P*≤0.05, and ns: *P* >0.05 (not significant). (E) 19 mRNAs tested for mRNA abundance by smiFISH with and without maternal *spn-4* activity. The ratio of mRNA abundance in *spn-4(tm291)* null mutants versus the wild type (strain N2) at the 4-cell stage is plotted above the enrichment values for each RNA-binding protein and transcript abundance. Statistics as in (D).

Using our assay and embryos derived from *spn-4* balancer animals, we compared the abundance of 12 SPN-4-associated transcripts (*lin-41, chs-1, cpg-2, oma-2, car-1*, C04B4.2, *nos-1, egg-3*, R05H11.1, *cgh-1, vab-2, npr-35*), 3 transcripts associated with OMA-1 and/or LIN-41 (Y19D2B.2, Y37H2A.12, and ZK666.4) and 4 controls (*mcm-2, mex-5, nasp-2*, and *pigv-1*) in embryos with and without maternal *spn-4* activity (Fig. 3C,D and S5). These transcripts provide a wide range of expression levels, SPN-4 enrichment values, and associations with the other two RNA-binding proteins (Fig. 3E). Altogether 9 of the 12 SPN-4-associated mRNAs exhibited a statistically significant increase in transcript abundance in *spn-4(tm291)* null mutant embryos compared to the wild type (e.g., *lin-41, chs-1, cpg-2, oma-2, car-1*, C04B4.2, *nos-1, egg-3*, and R05H11.1). In contrast, only one of the seven control transcripts (four with no binding; three with only LIN-41 and/or OMA-1 binding) exhibited an increase in abundance in *spn-4(tm291)* embryos. Together, these results indicate that SPN-4 specifically regulates the stability of its target transcripts.

The timing of *spn-4*-dependent mRNA clearance was gene-specific, occurring at or before the 2-cell stage for *lin-41* and *chs-1*, and slightly later for *cpg-2* and *oma-2* (Fig. 3D). We compared maternal mRNA clearance in wild-type embryos and those expressing maternal *spn-4::gfp* (Fig. S4 and S5). We observed that transcripts were modestly increased in *spn-4::gfp* embryos (Fig. S4B) suggesting a mild hypomorphic phenotype. Comparing *spn-4(tm291)* with SPN-4::GFP-expressing embryos therefore underestimates the magnitude of the effect of *spn-4* activity on mRNA clearance (Fig. S5). We examined *chs-1* transcript abundance in the oocytes of wild-type dissected gonads and detected 12,052 ± 1,024 (n=15) *chs-1* transcripts, similar to the number of *chs-1* transcripts observed in *spn-4(tm291)* null mutant embryos in which maternal mRNA clearance of this transcript appears to be blocked (Fig. 3D). This is consistent with a model in which *spn-4* is responsible for maternal mRNA clearance at early embryonic stages.

### The divergent Rbfox protein SPN-4 binds Rbfox motifs to promote mRNA clearance

Nuclear Rbfox proteins promote alternative splicing (Conboy, 2017), and the RNA Recognition Motif (RRM) within these proteins binds with nanomolar affinity to consensus Rbfox motifs 5’-GCAUG-3’ (Auweter et al., 2006; Ye et al., 2023). The *Drosophila* cytoplasmic Rbfox protein represses the translation of *pumilio* mRNA in the ovary by binding to 5’-GCAUG-3’ motifs (Carreira-Rosario et al., 2016). Intriguingly, the *lin-41* and *chs-1* 3’UTRs each contain a single 5’-GCAUG-3 Rbfox motif. We used genome editing to delete the *lin-41* Rbfox motif (Fig. 4A). A 38 bp deletion, *lin-41(*Δ*1006-1043)* or small FoxΔ (Fig. 4A), interfered with *lin-41* mRNA clearance; we observed elevated levels of *lin-41* mRNA in 2- and 4-cell embryos (Fig. 4B-D). Larger deletions encompassing the *lin-41* Rbfox motif, *lin-41 (*Δ*762-1112)* and *lin-41 (*Δ*777-1054)* or large and medium FoxΔ, respectively (Fig. 4A), had a slightly stronger effect (Fig. 4B-D). These results suggest that the Rbfox motif mediates *lin-41* mRNA clearance, but that adjacent sequences also participate. Stoeckius et al. (2014) noted that many of the mRNAs subject to early maternal mRNA clearance, including *lin-41*, have a polyC motif in their 3’UTR. We created small, *lin-41 (*Δ*311-326)*, and large, *lin-41 (*Δ*71-498)*, deletions that remove the *lin-41* polyC motif (Fig. 4A) but observed no significant effect on *lin-41* mRNA clearance (Fig. 4C,D and S6). It remains possible that the polyC motif is important later in embryogenesis.

**Fig. 4.**
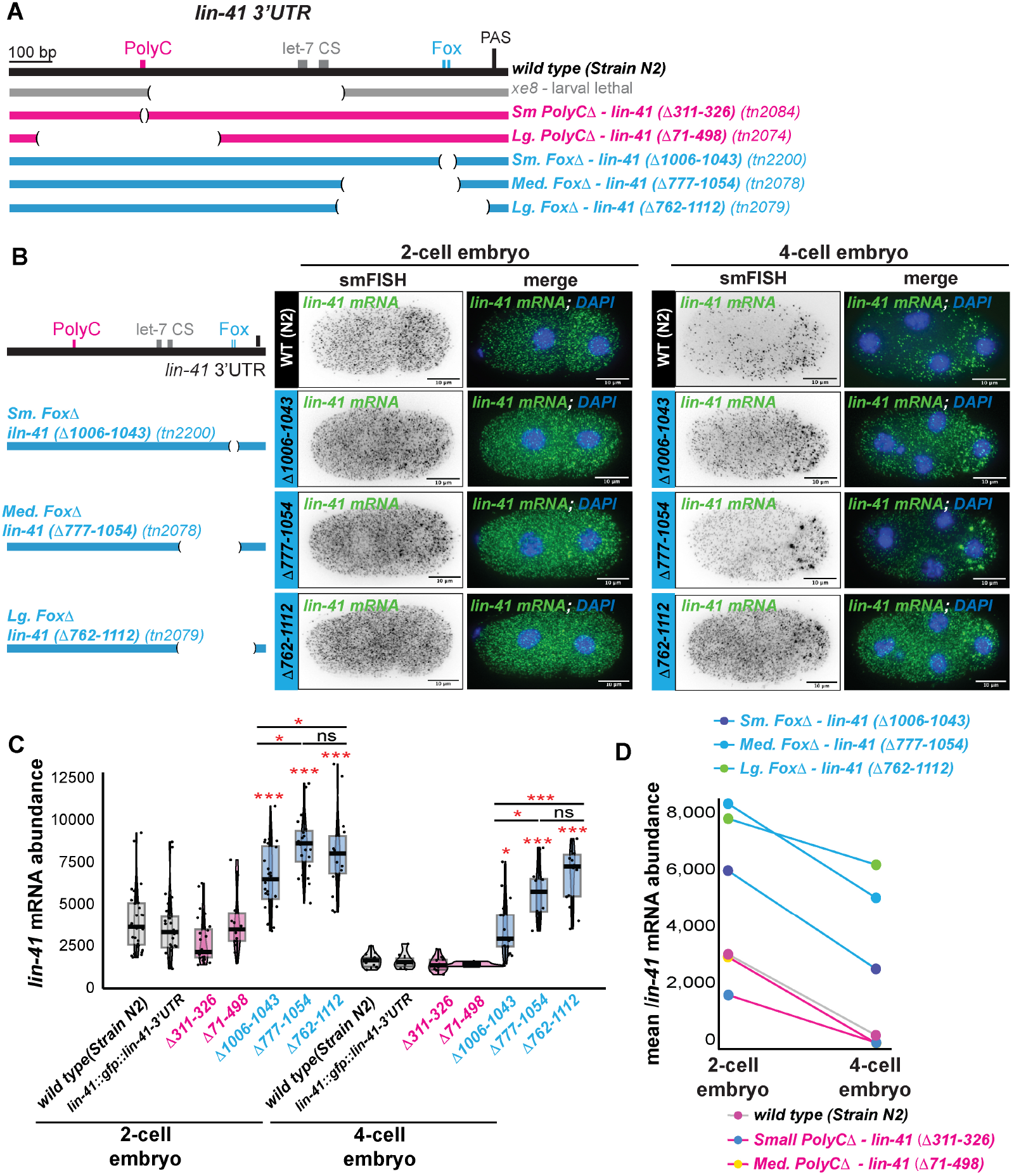
An Rbfox1-binding element in the *lin-41* 3’UTR is required for *lin-41* mRNA clearance. (A) The *lin-41* 3’UTR showing deletions generated by genome editing in the *lin-41(tn1541[gfp::lin-41])* genetic background. The *xe8* deletion affects regulation in larvae by the *let-7* microRNA that is not relevant in embryonic stages (Ecsedi et al., 2015) and is shown for comparison. (B) Representative micrographs of the Rbfox-motif deletions compared to full length. Total *lin-41* mRNA abundance was measured using smFISH in 3D across all z-stacks. Images shown are single-channel representations (left) or merged with DAPI DNA staining (right). Bars,10 µm. (C) Quantification of mRNA abundance for all *lin-41* 3’UTR deletion strains assayed. *n*-values ranged from five to 31 for each condition combination over 2-3 replicates. Statistical tests were performed within each cell stage using one-way ANOVA analysis followed by Tukey’s Honest Significant Differences (HSD) post-hoc analysis that accounts for multiple testing. *** *P*-adj <0.001; ** *P*-adj <0.01; * *P*-adj <0.05; ns = not significant; see Table S6 for exact *P*- and *n*-values. (D) Mean *lin-41* mRNA abundance between the 2-to 4-cell stages for each deletion strain.

Removing SPN-4 elevated *lin-41* transcript levels (Fig. 3B-E), as did removing the Rbfox motif from the *lin-41* 3’UTR (Fig. 4B-D). To test whether SPN-4 affects *lin-41* transcript levels through the Rbfox motif or functions independently, we quantified the number of *lin-41* transcripts when the *spn-4(tm291)* null mutation was combined with small FoxΔ [*lin-41 (*Δ*1006-1043)*]. We observed that the *spn-4(tm291)* null mutation did not synergize with small FoxΔ (Fig. 5A-C). Similarly, combining small FoxΔ and the hypomorphic *spn-4::gfp* allele did not further elevate the levels of *lin-41* transcripts (Fig. S7). These results suggest that SPN-4 functions through the Rbfox motif to decrease *lin-41* transcript levels.

**Fig. 5.**
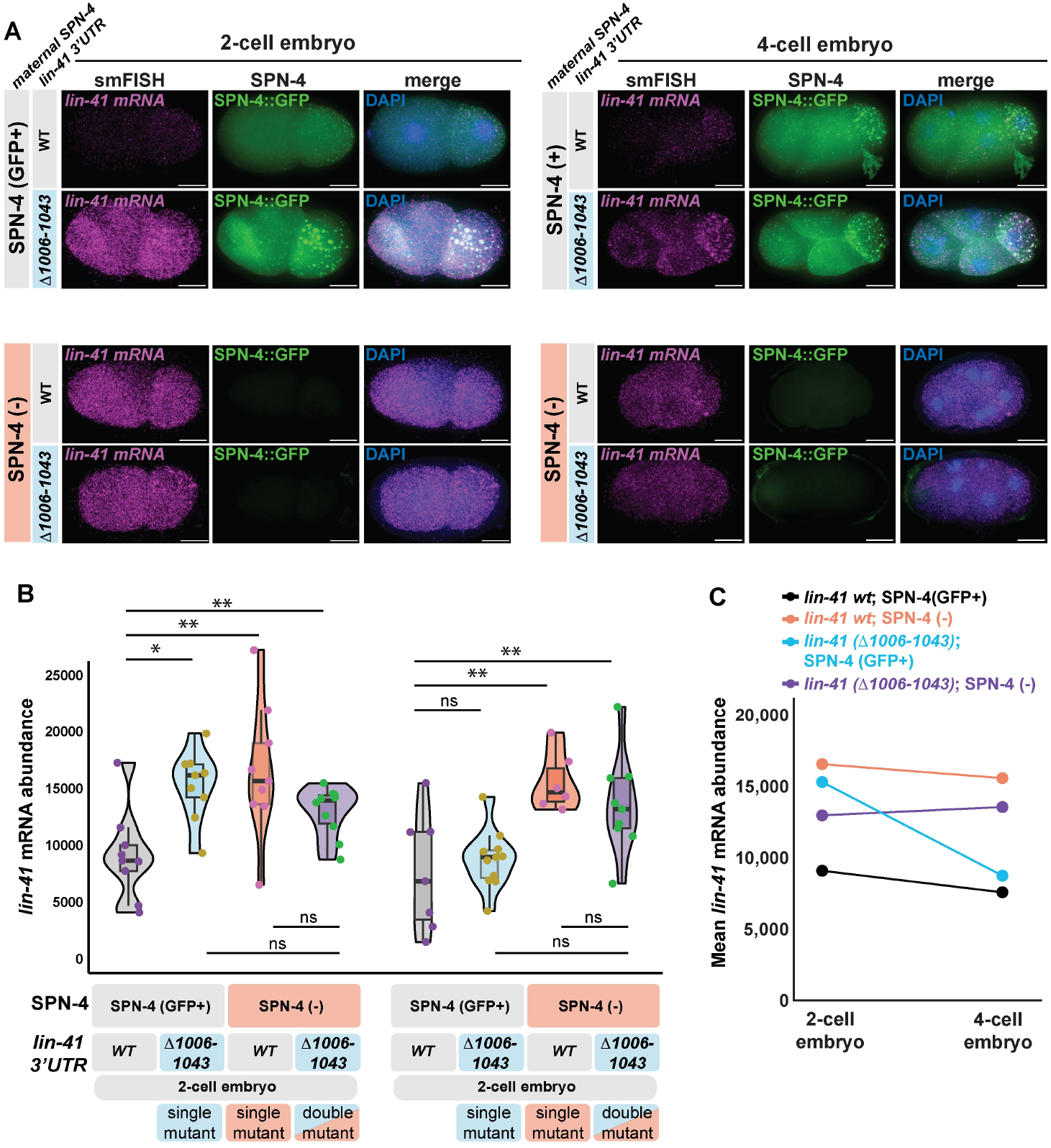
SPN-4 promotes *lin-41* maternal mRNA clearance via an Rbfox-binding motif. (A) Double mutants were constructed by combining a small Rbfox-motif deletion (Δ*1006-1043)* with the *spn-4* null mutation. Micrographs showing *lin-41* mRNA levels by smFISH, comparing embryos with and without maternal *spn-4::gfp* activity in the presence of a wild-type *lin-41* 3’UTR or *lin-41* containing a small Rbfox-motif deletion (Δ*1006-1043)*. Embryos were obtained from strains DG5256 *lin-41(+) spn-4(tm291)/tmC3[spn-4(tn2056[spn-4::gfp])]* or DG5965 *lin-41(tn2239 fox*Δ*); spn-4(tm291)/tmC3[spn-4(tn2056[spn-4::gfp])*. Embryos without maternal *spn-4::gfp* activity lack GFP fluorescence. Images are single-channel maximum intensity projections (smFISH only, left), GFP fluorescence (middle), and merged with DAPI stain (right). Bars,10 µm. (B) Quantification of *lin-41* mRNA abundance, represented as violin and box plots with jittered data points. *n*-values ranged from 6-31 over 2-4 replicates. Statistical tests for each cell stage were performed using one-way ANOVA followed by Tukey’s HSD post-hoc analysis accounting for multiple testing. \*\*\**P*-adj <0.001; \*\**P*-adj <0.01; \**P*-adj >0.05; ns = not significant; see Table S7 for exact *P*- and *n*-values. (C) Mean *lin-41* mRNA abundance across the two cell stages for each strain.

Likewise, we deleted the Rbfox motif in the *chs-1* 3’UTR, creating *chs-1(*Δ*56-72)* FoxΔ and found this interfered with the clearance of *chs-1* mRNA (Fig. 6A-D). We next asked which sequence motifs are enriched in the 3’UTR sequences of SPN-4-associated mRNAs. The top-ranking sequence motifs strongly resemble the consensus-binding site for Rbfox proteins but also include the 5’-extension UUUAUU (Fig. 6E). *lin-41* and *chs-1* 3’UTRs contain the extended-motif sequence as well as degenerate matches to Rbfox motif sequences (Fig. 6G).

**Fig. 6.**
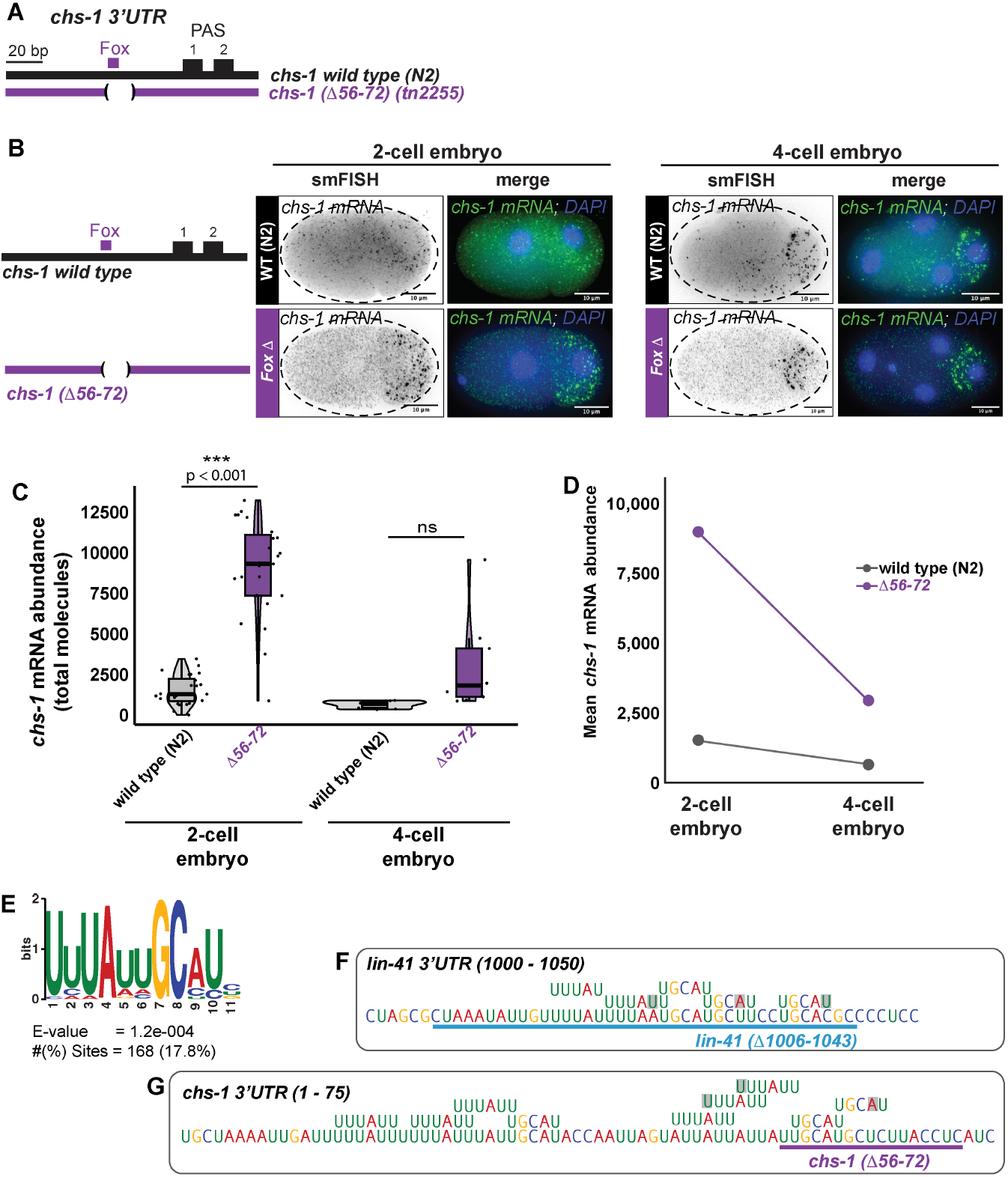
An Rbfox-binding motif is required for *chs-1* mRNA clearance. (A) The *chs-1* 3’UTR region showing the location of the Rbfox motif and the *chs-1(tn2255* Δ*56-72)* deletion mutation. (B) Micrographs of *chs-1* mRNA detected in the wild type and the *chs-1* mutant using smFISH in a single channel (left) or merged with DAPI (right). Bars, 10 µm. (C) Violin and box plots with jittered data points showing *chs-1* mRNA abundance from the full dataset. *n*-values ranged from 9-24 for each condition combination across 2-3 reps. Statistical tests for each cell stage were performed using one-way ANOVA followed by Tukey’s HSD post-hoc analysis accounting for multiple testing. *** *P*-adj <0.001; ** *P*-adj <0.01; * *P*-adj <0.05; ns = not significant; see Table S8 for exact *P*- and *n*-values. (D) Mean *chs-1* mRNA abundance across the two cell stages. (E) Motif discovery using 3’ UTR regions of SPN-4-associated mRNAs as the query set and the 3’UTR regions of OMA-1- and LIN-41-associated mRNAs as the background set. (F,G) Locations within the 3’UTRs of *lin-41* (F) or *chs-1* (G) where degenerate matches to the sequence motif or sections of it are found.

To determine whether SPN-4 can directly recognize Rbfox motifs, we assessed sequence-specific RNA binding of the SPN-4 RRM using electrophoretic mobility shift assays (EMSA). We observed specific and saturable binding to a 60-nucleotide RNA probe containing the Rbfox motif, *lin-41 (995-1054)*, with a measured K_d_ of 307.7 ± 17.9 nM (Fig. 7A,C and S8A,C). A 25-nucleotide RNA probe containing the Rbfox motif, *lin-41*(1023-1047), had slightly stronger binding with a measured K_d_ of 285.2 ± 29.4 nM (Fig. 7C and S8B,D). Only non-saturable binding was observed for a 43-nucleotide RNA probe in which the Rbfox motif region had been removed, *lin-41 (995-1027, 1043-1054)*, and a 40-nucleotide RNA probe from another region of the *lin-41* 3’UTR, *lin-41 (141-180)* (Fig. 7B,C and S9A-D).

**Fig. 7.**
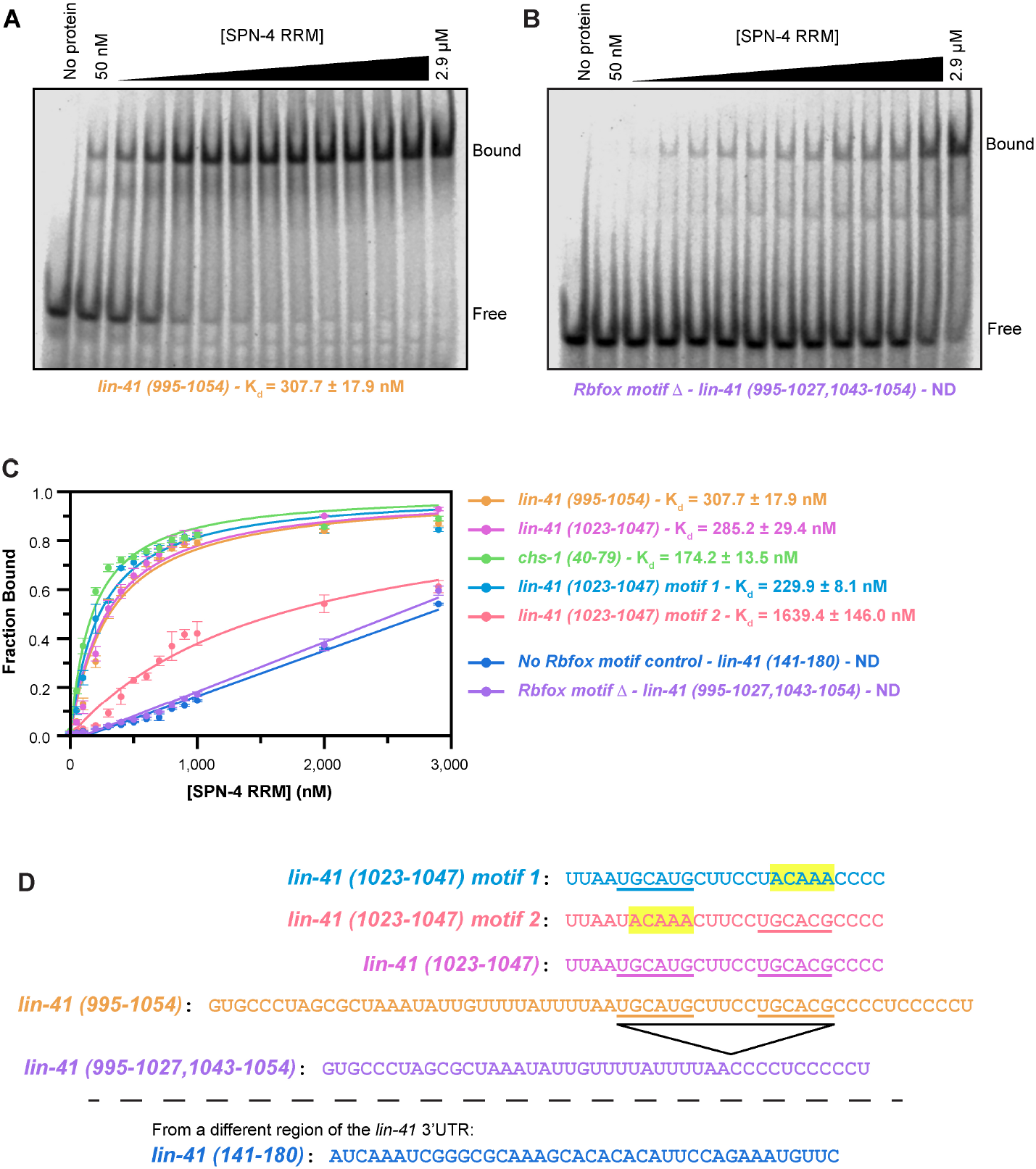
The SPN-4 RRM binds Rbfox-motif sequences in the *lin-41* and *chs-1* 3’UTRs. (A,B) Binding of the SPN-4 RRM to RNA to a 60-nt *lin-41 (995-1054)* 3’UTR RNA probe (A) or a 43-nt *lin-41 (995-1027,1043-1054)* deletion probe lacking the Rbfox motif (B). (C) Binding curves and the dissociation constants for all the SPN-4 RRM interactions analyzed. ND = Not Determined. (D) Sequences of all *lin-41* probes used for binding assays.

The *lin-41 (1027-1032)* 3’UTR region contains a match to the Rbfox consensus sequence, 5’-GCAUG-3’ (motif 1), as well as a degenerate match, 5’-GCACG-3’ from position 1038-1043 (motif 2) (Fig. 7D). Published results suggest that Rbfox proteins can also recognize 5’-GCACG-3’ sequences (Ye et al., 2023). We investigated whether both motifs contribute to binding by analyzing the effects of base substitutions known to interfere with base recognition by Rbfox. Motifs 1 and 2 were each individually replaced with 5’-ACAAA-3’ in the context of the 25-nt RNA probe (Fig. 7D). The SPN-4 RRM bound the RNA probe containing an intact version of motif 1 (*lin-41 (1023-1047)* motif 1) with a K_d_ of 229.9 ± 8.1 nM, which is similar to its affinity to the original 25-nt probe (Fig. 7C and S10A,C). In contrast, the SPN-4 RRM bound the RNA probe containing an intact version of motif 2 (*lin-41 (1023-1047)* motif 2) relatively weakly with a K_d_ of 1639.4 ± 146.0 nM (Fig. 7C and S10B,D). Thus, motif 1, which matches the Rbfox consensus sequence is sufficient to mediate *in vitro* binding.

The *chs-1* 3’UTR also contains an Rbfox motif necessary for mRNA clearance (Fig. 6). The SPN-4 RRM binds to a 40-nucleotide RNA probe containing this sequence, *chs-1(40-79)*, with a measured K_d_ of 174.2 ± 13.5 nM (Fig. 7C and S11). Collectively, these results suggest that SPN-4 can directly recognize RNA sequences within the 3’UTRs of *lin-41* and *chs-1* to promote their elimination in the early embryo.

### A genetic screen identifies components of the CCR4-NOT deadenylase complex as candidate effectors of SPN-4-mediated maternal mRNA clearance

PUF-3 and PUF-11 are nearly identical Pumilio-related RNA-binding proteins that, like LIN-41, mediate the translational repression of SPN-4 in oocytes (Hubstenberger et al., 2012; Tsukamoto et al., 2017). In a wild-type genetic background, the expression of SPN-4::GFP is limited to the –1, and to a lesser extent, the –2 oocyte (∼10% of gonad arms; Fig. 8A-E). In contrast, in *puf-11(q971) puf-3(q966)* double null mutants, referred to as *puf-3/11*Δ, SPN-4::GFP expression can be extended from the –1 to –4 oocytes (Fig. 8E). While making these strains, we observed that *spn-4(tn1699[spn-4::gfp])* weakly suppressed the maternal-effect lethality of *puf-3/11*Δ. *puf-3/11*Δ hermaphrodites have a brood size of 0 (n=22) and *puf-3/11*Δ;*spn-4(tn1699[spn-4::gfp])* hermaphrodites exhibit a brood size of 20.3 (n=12). Further, the *spn-4(tm291)* null mutation is a weak dominant suppressor— *puf-3/11*Δ; *spn-4(tm291)/+* hermaphrodites exhibit a brood size of 7.5 (n=24). Based on these results, we hypothesized that early expression of SPN-4 in proximal oocytes in *puf-3/11*Δ might induce precocious maternal mRNA clearance, causing lethality.

**Fig. 8.**
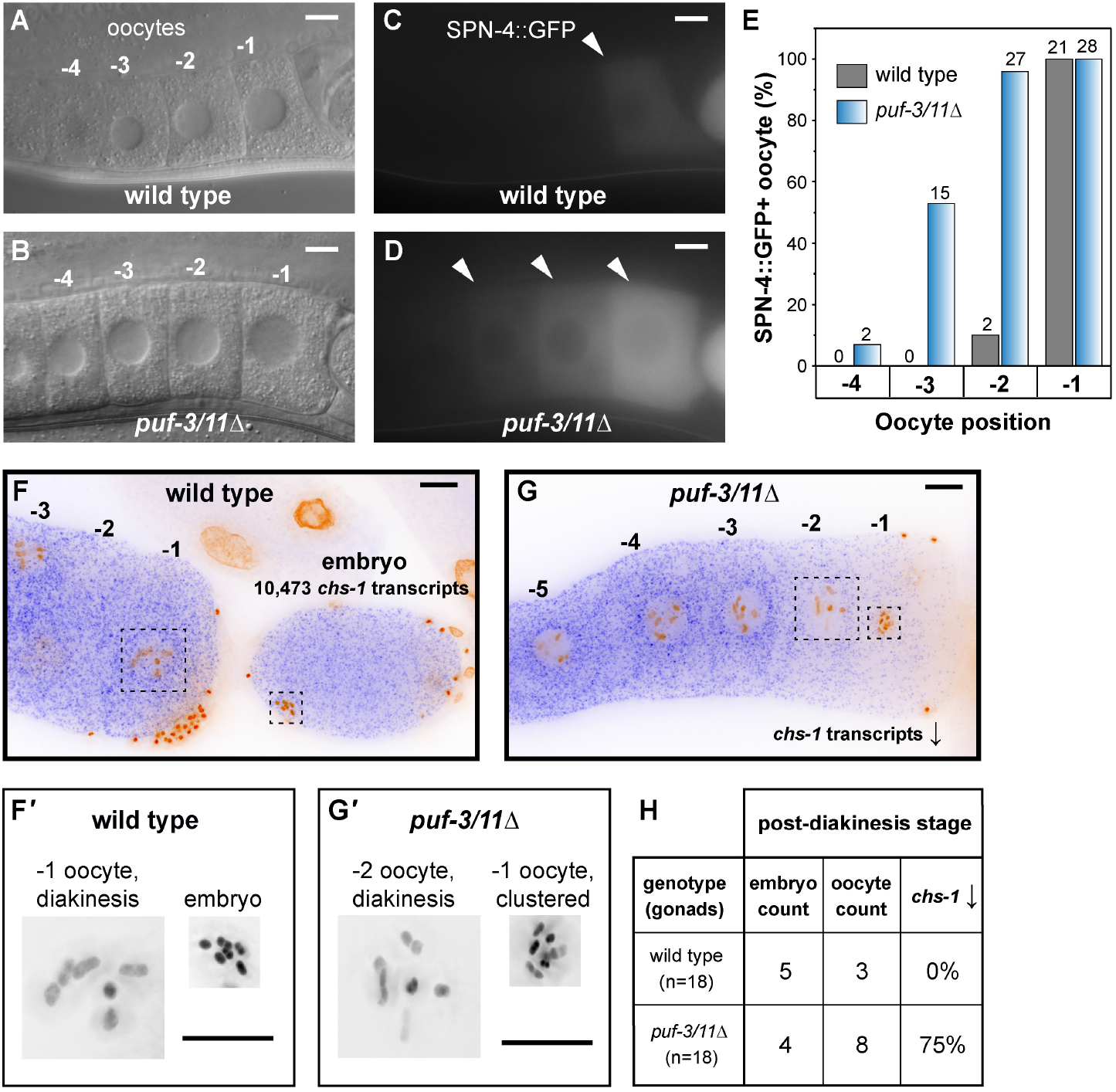
SPN-4 is ectopically expressed in proximal oocytes in *puf-3/11*Δ mutants. (A-E) Late-stage oocytes (A,B) express SPN-4::GFP (C,D; arrow-heads); expression is expanded in *puf-3/11*Δ mutants relative to the wild type (C-E). The percentage of SPN-4::GFP-positive oocytes at each position (E) with the number observed above each bar. (F-G’) *chs-1* transcripts (blue) are abundant in late-stage oocytes and newly fertilized embryos of wild-type animals but appear decreased in some *puf-3/11*Δ oocytes. Orange shows DNA (F,G); this channel is magnified to illustrate diakinesis and post-diakinesis nuclei (F’,G’). The newly fertilized wild-type embryo (F) was sufficiently separated from nearby oocytes to determine the total number of transcripts. (H) Total numbers of gonads and embryos or oocytes that had progressed beyond diakinesis are shown. Overt decreases in *chs-1* transcript levels in *puf-3/11*Δ oocytes and embryos are associated with meiotic progression beyond diakinesis. Proximal oocytes (–1 to –5) are indicated. Bars, 10 µm.

To test this hypothesis, we compared the number of *chs-1* transcripts in oocytes from the wild type and *puf-3/11*Δ (Fig. 8F,G and S12). Whereas wild-type oocytes express ∼12,000 *chs-1* transcripts (Fig. S12C), we frequently observe fewer *chs-1* transcripts (e.g., 2,000-10,000) in *puf-3/11*Δ oocytes or newly fertilized embryos within the spermatheca (Fig. 8G,H and S12B,D). We also noticed that some *puf-3/11*Δ oocytes exhibited an apparent acceleration of the meiotic cell cycle such that clustered chromosomes were observed in –1 oocytes in day-1 adults (Fig. 8G-H). This clustered chromosome morphology is characteristic of newly fertilized embryos in the wild type and correlated with an overt decrease in *chs-1* transcript levels in *puf-3/11*Δ mutants (Fig. 8F-H). We suggest that the precocious expression of SPN-4 in proximal *puf-3/11*Δ oocytes, together with a potential acceleration of meiotic progression, contributes to precocious and lethal maternal mRNA clearance; this inference will require further study.

Based on these results, we reasoned that mutations in other genes that affect maternal mRNA clearance might suppress *puf-3/11*Δ like *spn-4*. We sought dominant suppressors as the *spn-4(tm291)* null mutation exhibited dominance owing to haploinsufficiency. We screened 28,498 EMS-mutagenized haploid genomes and isolated 24 fertile suppressed *puf-3/11*Δ strains (Fig. 9A).

**Fig. 9.**
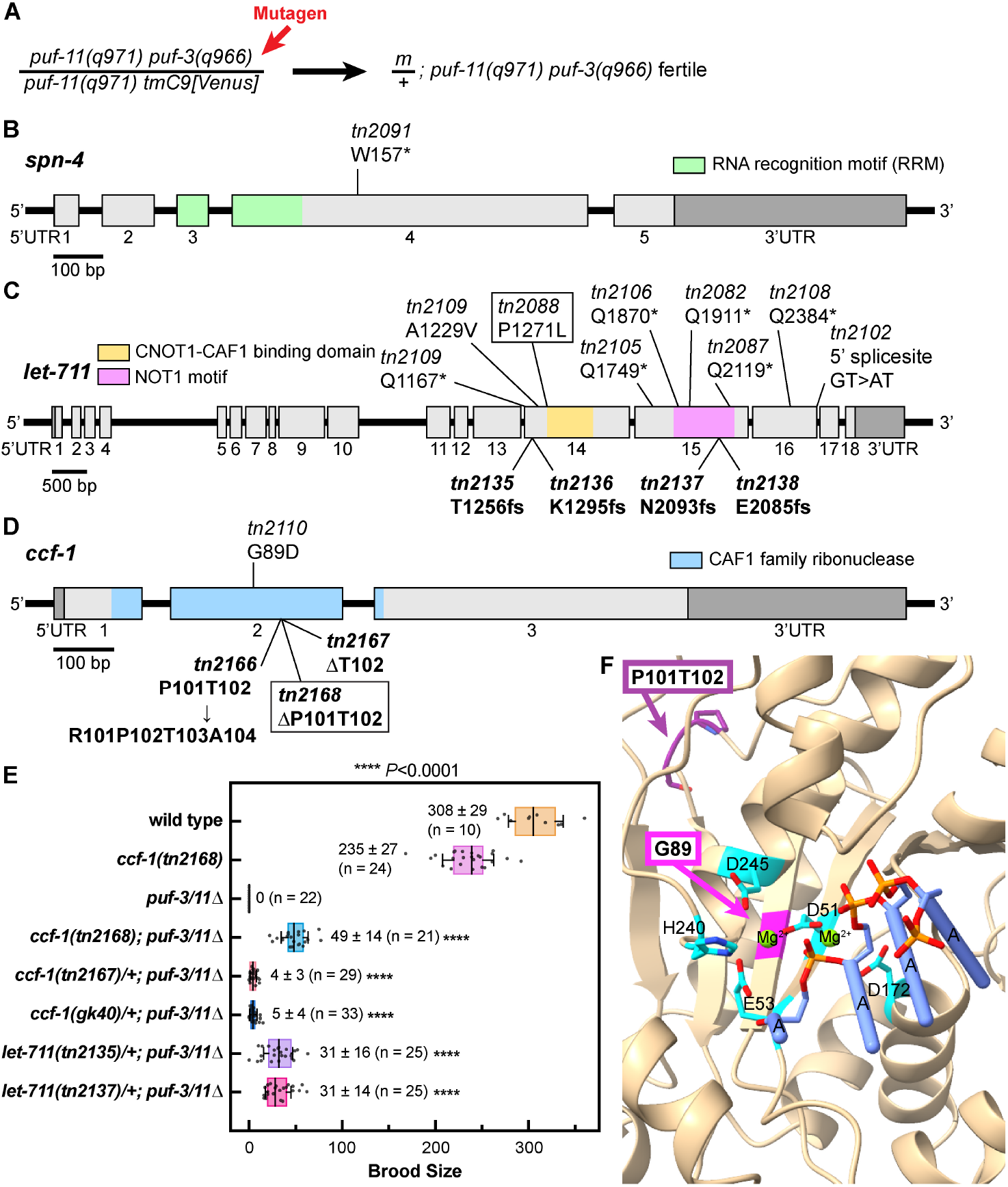
Mutations affecting the CCR4-NOT complex suppress *puf-3/11*Δ maternal-effect lethality. (A) A genetic screen for dominant suppressors of *puf-3/11*Δ maternal-effect lethality. (B-D) Mutations isolated as dominant suppressors of *puf-3/11*Δ maternal-effect lethality. Mutations isolated following EMS mutagenesis (roman font) and genome editing (bold font) are shown. Mutations that can be maintained as fertile homozygous strains are boxed. fs, frameshift mutations. (E) Brood-size measurements. A two-sample t-test with Welch’s correction was used to compare the suppressed strains to *puf-3/11*Δ. (F) A model of the CCF-1 active site with a poly(A) RNA substrate generated with AlphaFold and using information from the human CCR4-CAF1 deadenylase complex crystal structure (Chen et al., 2021). The active site is shown containing DEDDH residues (cyan) that coordinate divalent Mg^2+^ ions (green). The predicted positions of the G89 residue, affected by the *tn2110* G89D mutation, and the P101T102 residues, affected by the *tn2168* ΔP101T102 mutation, are shown.

Suppressors were backcrossed and candidate mutations identified by whole-genome and Sanger sequencing. We isolated a strong loss-of-function *spn-4* allele, validating the underlying rationale (Fig. 9B). In addition, we isolated 10 mutations affecting the CCR4-NOT deadenylase complex, 9 in *let-711* (Fig. 9C), which encodes the largest subunit, and 1 in *ccf-1* (Fig. 9D), which encodes one of two deadenylases. Six *let-711* mutations introduce premature stop codons and are likely strong loss-of-function alleles (Fig. 9C). *ccf-1(tn2110 G89D)* affects a conserved amino acid residue near the active site of CCF-1 (Fig. 9D,F) (Nousch et al., 2013).

To confirm that strong loss-of-function mutations in *let-711* dominantly suppress *puf-3/11*Δ, we targeted the *let-711* gene in the *puf-3/11*Δ background using genome editing. Sixteen suppressed strains were isolated, four were sequenced and all were heterozygous for *let-711* frameshift mutations (Fig. 9C). We also used a genome editing approach to confirm that *ccf-1* mutations could suppress *puf-3/11*Δ, targeting a region of *ccf-1* encoding amino acid residues near G89 and the active site. Five suppressed strains were isolated, three with altered amino acid sequences near the CCF-1 active site (Fig. 9D, F). In an otherwise wild-type genetic background, *ccf-1(tn2166* P^101^T^102^ to R^101^P^102^T^103^A^104^) and *ccf-1(tn2167* ΔT^102^)) are sterile (n>10), whereas *ccf-1(tn2168* ΔP^101^ΔT^102^) hermaphrodites are healthy fertile adults with a brood size of 235 ± 27 (Fig. 9E). For each gene, mutations were returned to a *puf-3/11*Δ background and verified to be suppressors.

*ccf-1(tn2168)* homozygotes exhibited the strongest suppression of *puf-3/11*Δ (Fig. 9E). Strong loss-of-function *let-711* heterozygotes were better dominant suppressors than infertile *ccf-1* mutations, including the *ccf-1(gk40)* deletion allele (Fig. 9E). Dosage of LET-711/NOT1 may be crucial as it plays a role in scaffolding the CCR4-NOT complex. Taken together, these genetic studies implicate the activity of the CCR4-NOT complex in SPN-4-dependent maternal mRNA clearance.

### The CCR4-NOT deadenylase complex is required for SPN-4-dependent maternal mRNA clearance

We tested whether the CCR4-NOT complex participates in SPN-4-mediated maternal mRNA clearance by depleting LET-711 and CCF-1 in late oogenesis using the auxin-inducible degradation (AID) system (Zhang et al., 2015). Protein depletion conditions, which include a degron-tagged allele, oocyte-expressed TIR1 and an auxin analog, will be referred to as LET-711AID and CCF-1AID, respectively. For both LET-711AID and CCF-1AID, highly penetrant embryonic lethality followed a brief (3 h) treatment with auxin analog (Fig. 10A,B) with a 3-4-fold increase in poly(A) RNA levels in early embryos (Fig. 10C-F’; n=5-17). The increase in poly(A) RNA signal could be due to increased mRNA stability or an increase in poly(A)-tail length or both. We next conducted two-color smFISH analyses for *lin-41* (Fig. 10G-J) and *chs-1* (Fig. 10K-N) mRNA, using the control *set-3* mRNA, which is not SPN-4-associated (Table S1). Quantitative analysis indicated substantially increased levels of *lin-41* mRNA and *chs-1* mRNA following LET-711AID and CCF-1AID relative to the wild type (e.g., > 7-fold median increases in 4-cell embryos; Fig. 10O,P and S13). Levels of *set-3* mRNA were only mildly affected (> 1.6-fold median increases; Fig. S13C). These results indicate the CCR4-NOT complex is required for the SPN-4-dependent clearance of the two targets we tested.

**Fig. 10.**
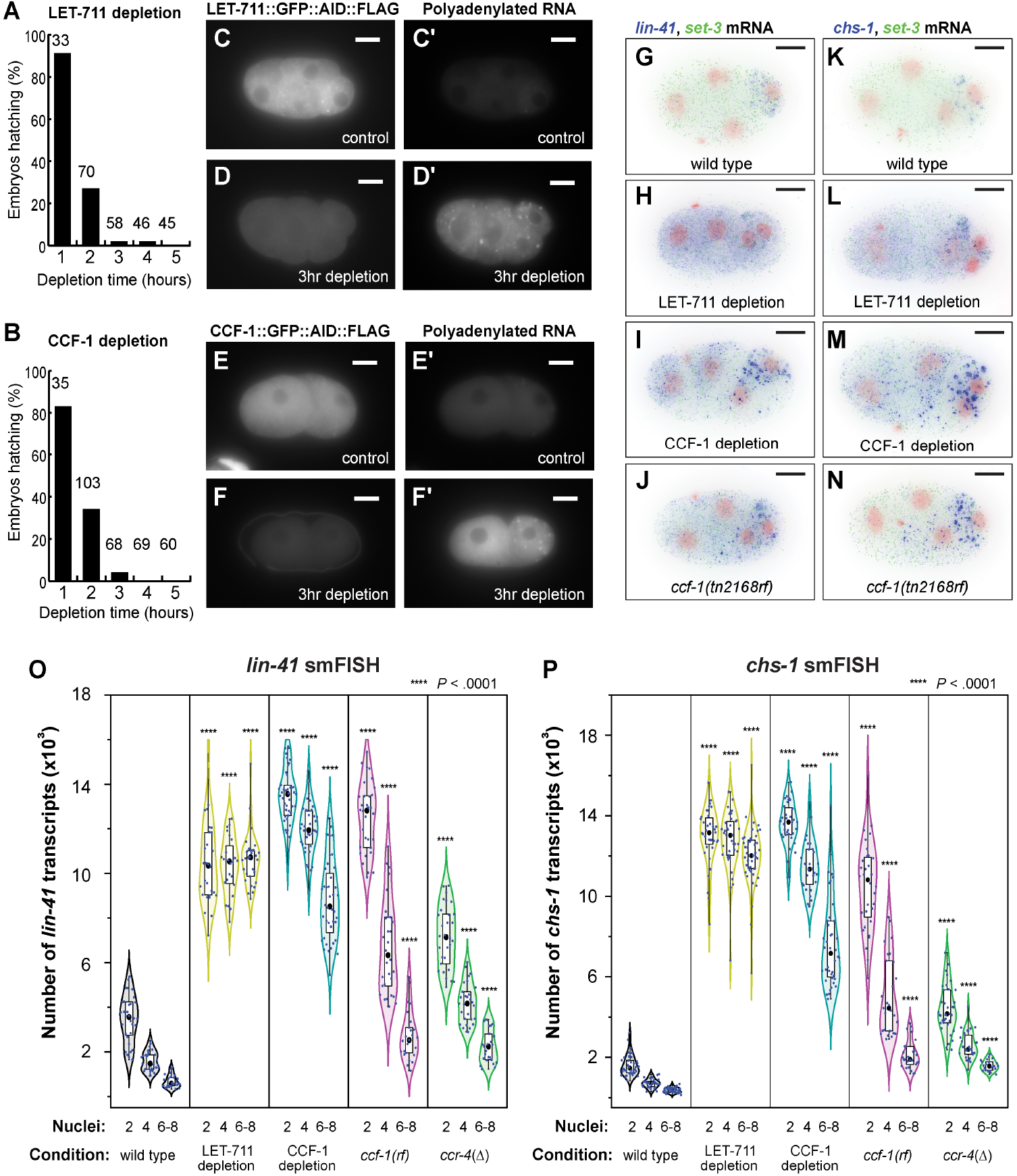
CCR-4-NOT subunits LET-711 and CCF-1 mediate *lin-41* and *chs-1* mRNA clearance. (A,B) LET-711 and CCF-1 depletion causes embryonic lethality 3-4 h after addition of auxin analog (K-NAA). The number of embryos examined is indicated. (C-F’) LET-711::GFP::AID*::3xFLAG expression (C,D) or CCF-1::GFP::AID*::3xFLAG expression (E,F) and Poly-A+ RNA detected by FISH (C’-F’) in 4-cell embryos after a 3 h K-NAA treatment of the parents. The TIR1 cofactor required for LET-711 and CCF-1 depletion is absent from (C,C’,E, E’) but present in (D,D’,F,F’). (G-N) *lin-41* (blue in panels G-J), or *chs-1* (blue in panels K-N), and *set-3* (green) transcripts in wild-type, LET-711 depletion, CCF-1 depletion, or *ccf-1(tn2168rf)* embryos with 4 nuclei (DNA, salmon). (O-P) Violin and box plots with individual data points depicting *lin-41* (O) or *chs-1* (P) transcript levels. Two or three replicate experiments for each genotype; total *n*-values > 21 at each developmental stage. Significance assessed at each stage using Welch’s one-way ANOVA with a Games-Howell post-hoc test accounting for unequal variances and multiple comparisons; it is reported relative to the wild type. See Table S9 for exact *P*- and *n*-values. Bars, 10 µm.

Both the *let-711(tn2130)* and *ccf-1(tn2132)* degron-tagged alleles, despite being viable and fertile, represent hypomorphic conditions. We noted significantly elevated levels of *lin-41* and *chs-1* transcripts in early embryos carrying each degron-tagged allele on its own (Fig. S13). In these embryos, the levels of *lin-41* mRNA and *chs-1* mRNA decline rapidly—most notably when compared to LET-711AID and CCF-1AID—as embryos progress from the 2-to the 6-8-cell stages, eventually approaching levels similar to those observed in the wild type (Fig. S13). Similar transcript degradation kinetics were observed for the viable and fertile *ccf-1(tn2168)* reduction-of-function allele originally identified as a *puf-3/11*Δ suppressor (Fig. 10O,P and S13).

The *ccr-4(tm1312)* null mutation, which affects the second deadenylase in the complex, is viable and fertile with a slightly reduced brood size (Nousch et al., 2013). We tested whether CCR4 plays a role in maternal mRNA clearance by examining *lin-41* and *chs-1* mRNA transcripts in *ccr-4(tm1312)* embryos. Both transcripts were moderately elevated and declined rapidly, approaching levels seen in wild-type embryos by the 6-8 cell stage (Fig. 10O,P and S13). We next generated *ccf-1(tn2168rf); ccr-4(tm1312null)* double mutants and found they produce only ∼5 progeny that grow to late larval or early adult stages, due to a combination of reduced fertility, embryonic lethality and larval arrest (n=2256 eggs laid by 24 parents). Inviability of the *ccf-1(tn2168rf); ccr-4(tm1312 null)* strain is consistent with the idea that both CCF-1 and CCR-4 contribute to maternal mRNA clearance.

## DISCUSSION

Maternally provided RNA-binding proteins figure prominently in the process of early maternal mRNA clearance (Despic and Neugebauer, 2018). In this study, we report a mechanism of early maternal mRNA clearance in *C. elegans* that involves SPN-4, a divergent member of the Rbfox RNA-binding protein family. For two maternal transcripts we have studied in detail, *lin-41* and *chs-1*, SPN-4-binding sites within the 3’UTR are required for transcript clearance. The 3’UTR sequences of these and other SPN-4-associated transcripts contain sequences that are strikingly similar to Rbfox motifs. We estimate that this mechanism is responsible for the elimination of approximately 11% of maternal transcripts fated for decay.

Our genetic and smFISH experiments suggest that the CCR4-NOT deadenylase complex is the major effector of SPN-4-dependent maternal mRNA clearance. A genetic screen designed to recover mutant alleles of genes that function with SPN-4 isolated alleles of *let-711* and *ccf-1*. Because components of the CCR4-NOT complex are required for viability, we utilized the AID system to deplete LET-711 and CCF-1 late in oogenesis and found that these treatments blocked elimination of *lin-41* mRNA and *chs-1* mRNA. Our analyses in *C. elegans* are consistent with findings from other systems that the CCR4-NOT deadenylase complex is a major effector of maternal mRNA clearance (Ma et al., 2015; Mishima and Tomari, 2016; Semotok et al., 2005; Sha et al., 2018; Soeda et al., 2023; Yu et al., 2016). The CCR4-NOT deadenylase complex also participates in translational regulation by microRNAs illustrating how multiple mRNA targeting triggers can deploy the complex to promote RNA decay or translational repression (Braun et al., 2011; Chekulaeva et al., 2011; Fabian et al., 2011). In *Drosophila, Xenopus* and zebrafish, zygotically expressed microRNAs mediate a late phase of maternal mRNA clearance (Bushati et al., 2008; Giraldez et al., 2006; Lund et al., 2009).

As we found for *C. elegans*, maternally provided RNA-binding proteins figure prominently in the process of early maternal mRNA clearance in other organisms, as they select specific transcripts for elimination by virtue of sequence-specific binding. For example, the *Drosophila* Smaug RNA-binding protein and its orthologs are key RNA regulators, influencing localization, degradation, translation, and phase state (Aviv et al., 2003; Carey et al., 2025; Chakravarty et al., 2020; Fernández-Alvarez et al., 2022; Semotok et al., 2005; Smibert et al., 1996; Tadros et al., 2007). Smaug directly binds many of its known targets to promote their elimination (Chen et al., 2014; Semotok et al., 2005; Tadros et al., 2007). Smaug also functions to repress target translation, as in the case of *nanos* mRNA that is not localized to the posterior pole of the *Drosophila* embryo (Dahanukar et al., 1999; Smibert et al., 1996). Smaug-dependent translational repression of *nanos* mRNA requires the eIF4E-T paralog Cup to outcompete translation initiation machinery. In contrast, Smaug-dependent mRNA decay requires the CCR4-NOT deadenylase complex (Pekovic et al., 2023; Semotok et al., 2005). The mechanism by which Smaug selects some mRNAs for elimination and others for repression is unclear but may relate to developmental timing and the cellular context. Indeed, many Smaug targets, including *nanos*, are subject to both fates (Chen et al., 2014). The translation of germ granule-associated *nanos* mRNA, despite its association with Smaug, depends on the germline determinant Oskar, which protects *nanos* mRNA from translational repression (Chen et al., 2024).

In *C. elegans*, SPN-4 and other RNA-binding proteins work together to repress expression of the *nanos* homolog *nos-2* in early embryonic blastomeres such that NOS-2 protein is first expressed in the P4 germline blastomere. This occurs through a 3’UTR-dependent mechanism (D’Agostino et al., 2006; Jadhav et al., 2008). The CCCH zinc-finger protein POS-1 activates expression of *nos-2* in P4 by interfering with SPN-4-mediated repression (Jadhav et al., 2008). It is currently unclear what determines whether SPN-4-associated mRNAs will be chosen for elimination or translational repression. It is possible that SPN-4, like *Drosophila* Smaug, is influenced by cellular and developmental context, possibly involving cell identity factors or cell cycle regulators to determine whether its targets are translationally repressed, destroyed, or both.

Studies in several systems have established a mechanistic connection between cell cycle transitions during oocyte meiotic maturation, egg activation, and early post-fertilization development and the process of maternal mRNA clearance. In *Drosophila*, the Pan Gu kinase complex becomes active during egg activation (Hara et al., 2017), and it promotes the translation of Smaug to trigger early maternal mRNA clearance (Tadros et al., 2007). In *C. elegans*, the RNA-binding protein LIN-41 is the key determinant of the extended meiotic prophase of *C. elegans* oocytes (Spike et al., 2014a). Late in oogenesis, LIN-41 is inactivated as a translational repressor and eliminated by ubiquitin-mediated protein degradation triggered by CDK-1 activation (Spike et al., 2014a; Spike et al., 2018). Since LIN-41 represses the translation of *spn-4* (Tsukamoto et al., 2017), this mechanism readies the oocyte for maternal mRNA clearance by generating SPN-4 ribonucleoprotein complexes (RNPs) containing many mRNAs that will be eliminated following fertilization. Importantly, our data suggest that SPN-4-associated mRNAs are stable in oocytes but destabilized in early embryos. The mechanistic basis for this is unclear but may involve the regulated recruitment of the CCR4-NOT complex to SPN-4 RNPs.

Our findings reveal a mutually antagonistic relationship between LIN-41 and SPN-4. During oogenesis, LIN-41 binds and translationally represses *spn-4* mRNA. Once LIN-41 repression is relieved, the resulting SPN-4 proteins can bind *lin-41* mRNA and promote its elimination thereby preventing LIN-41 translation in the embryo. This dual negative regulation, combined with the rapid clearance of LIN-41 protein (Spike et al., 2018), results in a dramatic boundary between LIN-41 marking late-stage oocytes and SPN-4 in early-stage embryos (Fig. 1A). This defines a dramatic cell state change at a critical point in the oocyte-to-zygote transition.

Our analysis of viable and fertile CCR4-NOT alleles demonstrated that some delays in maternal mRNA clearance are tolerated by the embryo. What seems to be important is that maternal mRNAs are sufficiently depleted by the time the zygotic genome is required. Our smFISH data in hypomorphic situations affecting the CCR4-NOT complex suggest that maternal mRNA clearance, although delayed, might be completed well before new transcription is required for viability at the ∼87-cell stage (Edgar et al., 1994). Thus, there appears to be some robustness in the system.

Two broad classes of models have been proposed for the function of maternal mRNA clearance (Tadros and Lipshitz, 2009). In the first scenario, the elimination of specific maternal transcripts prevents the synthesis of proteins that are toxic or incompatible with embryonic development. Indeed, maternal factors could hinder nuclear reprogramming or cell cycle regulation (Kojima et al., 2024). In a second scenario, maternal mRNA clearance might be required to establish the proper dosage of mRNAs needed to faithfully execute embryonic development or to diversify embryonic blastomeres. Several observations suggest that SPN-4-dependent mRNA clearance may be needed, in part, to provide a clean slate for establishing the zygotic transcriptome. A key observation is that several proteins encoded by SPN-4-associated transcripts are subjected to a second layer of control—protein degradation in late-stage oocytes or early embryos. For example, LIN-41 and the OMA proteins are subjected to ubiquitin-mediated protein degradation in the early embryo (Nishi and Lin, 2005; Shirayama et al., 2006; Spike et al., 2018; Stitzel et al., 2006) and CHS-1 protein is rapidly internalized and degraded after eggshell formation (Maruyama et al., 2007). Although we cannot exclude the possibility that the protein products of some SPN-4-associated transcripts are incompatible with embryonic development, mRNA clearance can be delayed without deleterious consequences in several mutant backgrounds. Further, *spn-4* null mutants exhibit highly pleiotropic defects that manifest within the first few cell divisions. Based on these observations, we suggest that SPN-4-dependent mRNA clearance is required for sculpting the embryonic transcriptome.

Maternal mRNA clearance is vital in the transfer of regulatory control from gamete to zygote across metazoans. In humans, mutations in the maternal-effect gene, BTG4, disrupt the process of maternal mRNA clearance and block embryonic cell divisions, accounting for instances of human infertility and failure of *in vitro* fertilization (Zheng et al., 2020). In mice, the BTG4 protein functions by recruiting the CCR4-NOT complex to its targets (Sha et al., 2018; Yu et al., 2016). These observations illustrate how features of maternal mRNA clearance are evolutionarily conserved and how our findings in *C. elegans* contribute to a greater understanding of this process.

## MATERIALS AND METHODS

### Strains, genetic analysis, and whole-genome sequencing for mutant identification

#### Worm husbandry

The genotypes of strains used in this study are reported in Table S2. Genes and mutations are described in WormBase (Sternberg et al., 2024) or in the indicated references. Culture and genetic manipulations were conducted at 20°C as described (Brenner, 1974), except for the analysis of strains containing *fog-1(q253ts)* or *spe-9(hc88ts)* which were propagated at the permissive temperature of 15°C and analyzed at the non-permissive temperature of 25°C. *C. elegans* were maintained on nematode growth medium (NGM: 3 g l^-1^ NaCl; 17 g l^-1^ agar; 2.5 g l^-1^ peptone; 5 mg l^-1^ cholesterol; 1 mM MgSO_-4_; 2.7 g l^-1^ KH_2_PO_4_; 0.89 g l^-1^ K_2_HPO_4_) and fed standard *E. coli* strains OP50 or OP50-1, except when grown on peptone-enriched medium in which strain NA22 was used. The balancer chromosomes used to maintain sterile *ccf-1* alleles were *tmC29[unc-49(tmIs1259)]* III (Dejima et al., 2018) or *hT2[umnIs60(myo-2p::mKate2)* I; *hT2[bli-4(e937)]* III. Strong loss-of-function *let-711* alleles were maintained using *hT2[umnIs60(myo-2p::mKate2)* I; *hT2[bli-4(e937)]* III. *ccr-4(tm1312)* was routinely maintained heterozygous with *tmC9[F36H1*.*2(tmIs1221)]* IV. *puf-3/11* strains contained a modified *tmC9* balancer chromosome A recombinant chromosome in which *puf-11(q971)* was linked to the *myo-2p::Venus*-tagged chromosome rearrangement *tmC9[F36H1*.*2(tmIs1221)]* IV (Dejima et al., 2018) was used to balance the *puf-11(q971) puf-3(q966)* double mutant (Haupt et al., 2020). This chromosome was identified by picking many uncoordinated *tmC9*-homozygous progeny from *puf-11(q971) +/+ tmC9* parents, allowing them to lay eggs, and then testing each animal for the *puf-11(q971)* deletion using three-primer PCR (Table S3). Six recombinants were identified among the 83 animals tested. A *puf-11(q971) tmC9* homozygote was identified in the next generation and used to create the DG5324 *puf-11(q971) tmC9[F36H1*.*2(tmIs1221)]* strain. The recombinant balancer chromosome was used to generate DG5365 *puf-11(q971) puf-3(q966)/puf-11(q971) tmC9* and additional strains. In all strain constructions, *puf-11(q971), puf-3(q966)* and *tmC9[F36H1*.*2(tmIs1221)]* were genotyped using a combination of phenotypic analysis and three-primer PCR (Table S3) when a genetic cross permitted the possibility of recombination between previously linked loci [e.g., *puf-11(q971)* and *puf-3(q966)* or *tmC9*].

#### Brood counts

To determine the number of viable progeny produced by adult hermaphrodites of the indicated genotypes, L4-stage hermaphrodites were cultured individually and transferred to new media every 12-24 h. Viable progeny were counted before they reproduced. Where indicated in the text, the number of embryos that were laid, hatched, and grew to late larval stages were counted. To generate heterozygous *let-711* or *ccf-1* mutant alleles in the *puf-11(q971) puf-3(q966)* genetic background for brood counts, mutant animals of genotype *mutant/hT2; puf-11(q971) puf-3(q966)/puf-11(q971) tmC9* were crossed to *puf-11(q971) puf-3(q966)* males. Cross progeny without either balancer chromosome were identified and used for brood counts.

#### Genetic screen for dominant suppressors of *puf-3/11* maternal-effect lethality

To isolate dominant mutations able to suppress the maternal-effect lethality of *puf-11(q971) puf-3(q966)* double mutants, L4-stage hermaphrodites of genotype *puf-11(q971) puf-3(q966)/puf-11(q971) tmC9* were mutagenized with 50 mM ethyl methanesulfonate (EMS) (Brenner, 1974). 10 individual homozygous *puf-11(q971) puf-3(q966)* L4 animals from the first filial (F1) generation were cultured on each of many 60 mm x 15 mm Petri dishes and suppressor mutations were recognized by their ability to grow and reproduce and exhaust the food supply of the culture. 24 suppressed strains were obtained from the progeny of 14,250 F1 animals. The suppressor strains were backcrossed to *puf-11(q971) puf-3(q966)* males between one and six times for whole-genome sequencing.

#### Whole genome sequencing for mutant identification

Because *spn-4* was a predicted target of the genetic screen, the suppressor mutants isolated were subjected to Sanger sequencing for *spn-4*, which identified *spn-4(tn2091 W157stop)* in the heterozygous state. Whole genome sequencing validated this mutant as a heterozygote. For whole genome sequencing, genomic DNA was prepared using the QIAGEN DNeasy Blood and Tissue Kit. Illumina libraries of genomic DNA were prepared and paired-end (2×150) sequenced by Azenta GENEWIZ to approximately 60-200x coverage. Reads were trimmed using TrimGalore (0.6.0) and mapped to the BSgenome.Celegans.UCSC.ce11 (1.4.2) genome using BWA mem (0.7.17-r1188). Aligned reads were sorted and quality filtered with Samtools (1.21). Duplicates were identified using MarkDuplicates (Picard, 2.18.16). The GATK Haplotype caller (4.1.2.0) was used to identify sequence variations with respect to the reference ce11 genome (Li and Durbin, 2009; Li et al., 2009; McKenna et al., 2010). The VariantAnnotation package (1.50.0) within R version (4.4.0) was used to filter variants for heterozygous and homozygous, single nucleotide polymorphisms with read depth ≥ 25 that were not present in all samples. These variants were annotated using the TxDb.Celegans.UCSC.ce11.ensGene (3.15.0) annotation package to consider all splice donor/acceptor mutations and non-synonymous protein coding mutations. Suppressed strains that possessed wild-type *puf-3(+)* sequences owing to the break-down of the *tmC9* balancer chromosome during mutagenesis, and the resulting formation of an extrachromosomal array, were not analyzed further.

#### Auxin-inducible degradation

A 100 mM filter-sterilized solution of the water-soluble auxin analog 1-Napthaleneacetic acid, potassium salt, or K-NAA (PhytoTech Labs), was prepared and diluted to a 4 mM working concentration in sterile water. The K-NAA working solution was added to young gravid adults growing on OP50-1-seeded NGM plates in sufficient volume to cover the surface and diffuse to a final concentration of ∼0.5 mM in the substrate volume. In test experiments to determine efficacy (e.g., Fig. 10A,B), 0.625 ml of 4 mM K-NAA was added to 35 mm x 15 mm Petri dishes containing 5 ml NGM. Embryo viability was assessed by moving 5 or 10 of the treated animals at each time point to a pre-equilibrated plate containing 1 mM K-NAA for a 1 h egg-lay; embryos that failed to hatch after 24 h were considered dead. In subsequent smFISH or FISH experiments, 3.125 ml of 4 mM K-NAA was added to 100 mm x 15 mm Petri dishes containing 25 ml of NGM. Embryos were harvested and fixed after a 3-3.25-h incubation.

#### Genome editing

The plasmids and repair templates used for clustered regularly interspaced short palindromic repeats (CRISPR)-Cas9 genome editing are described in Table S3. Plasmids expressing single-guide RNAs (sgRNAs) under the control of the U6 promoter were created from plasmid pRB1017 and sequence-specific oligonucleotides as described (Arribere et al., 2014). Deletion repair templates were purchased as Ultramer DNA oligos (Integrated DNA Technologies). A *dpy-10(cn64)* co-conversion strategy was used to enrich for deletion edits (Arribere et al., 2014). *lin-41* and *chs-1* 3’UTR deletion injection mixes contained pJA58 (7.5 ng µl^-1^), AF-ZF-827 (500 nM), target-specific sgRNA plasmid(s) (25 ng µl^-1^ each), target-specific repair template (500 nM), and the Cas9-expressing plasmid pDD162 (50 ng µl^-1^). Essentially the same injection mix, lacking a target-specific repair template, was used to create the CRISPR-Cas9-induced alleles of *let-711* and *ccf-1* that suppress *puf-3(q971) puf-11(q966)* maternal-effect lethality. Two sgRNAs directed against *let-711* sequences, *let-711* sgRNA3 and *let-711* sgRNA4, were individually injected with other genome editing reagents into *puf-11(q971) puf-3(q966)/tmC9[myo-2p::venus] puf-11(q971)* hermaphrodites and non-balancer containing progeny were scored for suppression (fertility). The efficiency with which suppressed lines were recovered was 10.2% for *let-711* sgRNA3 (6 suppressed F2 lines, n=59) and 23.8% for *let-711* sgRNA4 (10 suppressed F2 lines, n=42) in contrast to a control *dpy-10* sgRNA, which generated no suppressed F2 lines (n=88). Using *ccf-1* sgRNA, we isolated 5 suppressed strains (3.9% efficiency, n=127). The plasmid repair templates used to tag *ccf-1* and *let-711* with *gfp::aid*::3xflag* were created from pJW1583 as described (Ashley et al., 2021; Dickinson et al., 2015). These injection mixes contained target-specific sgRNA plasmid (25 ng µl^-1^), target-specific plasmid repair template (10 ng µl^-1^), pDD162 (50 ng µl^-1^), and the injection marker *pMyo2::Tdtomato* (4 ng µl^-1^). Standard methods were used to identify insertions and subsequently remove the self-excising cassette (SEC) (Dickinson et al., 2015). DG4158 *spn-4(tn1699[spn-4::gfp::tev::3xflag]* was previously generated and described (Tsukamoto et al., 2017). To generate DG5254 *tmC3[tmIs1230 spn-4(tn2056[spn-4::gfp::tev::3xflag])*, used as a balancer in the smFISH experiments in Fig. 3, 5 and S5, the SEC injection mix used by Tsukamoto et al. (2017) was injected into *dpy-11(e224)/tmC3[tmIs1230]* adult hermaphrodites and four independent lines with correct targeting onto the *tmC3* balancer were identified (Tsukamoto et al., 2017). In all cases, genome edits were validated by sequencing through repair junctions using the PCR primers described in Table S3.

### Immunopurification of RNA-binding proteins and RNA sequencing of associated transcripts

#### Lysate preparation

The following strains were used for immunopurification: DG4398 *fog-1(q253ts)* I; *spn-4(tn1699[spn-4p::spn-4::tev::3xflag])* V, DG4400 *spe-9(hc88ts)* I; *spn-4(tn1699)* V, DG3923 *fog-1(q253ts) llin-41(tn1541[gfp::tev::s-tag::lin-41])* I, DG4485 *spe-9(hc88ts) lin-41(tn1541)* I, DG2566 *fog-1(q253ts)* I; *oma-1(zu405te33)* IV; *tnIs17[pCS410 oma-1p::oma-1::s-tag::tev::gfp, unc-119(+)]*, DG2581 *spe-9(hc88ts)* I; *oma-1(zu405te33)* IV; *tnIs17*. The temperature-sensitive alleles, *fog-2(q253ts)* and *spe-9(hc88ts)*, were used to generate sterile adults lacking sperm or containing fertilization-defective sperm, respectively. Four L4-stage larvae were cultured at 15°C on each of 16 60 mm x 15 mm Petri dishes with NGM seeded with the *E. coli* strain OP50-1 as food source. After culturing for 10 days, starved larvae were collected and transferred to 40 large 100 mm x 15 mm Petri dishes with peptone-enriched (20 g l^-1^ peptone) NGM seeded with *E. coli* strain NA22. After culturing at 15°C for five days, embryos were collected by alkaline hypochlorite treatment (20% bleach and 0.5N NaOH), washed in M9 buffer and allowed to hatch in 10 100 mm x 15 mm Petri dishes containing 25 ml of M9 solution. After incubating at 25°C for 1 day, L1-stage larvae were cultured at 25°C at a density of 30,000 L1-stage larvae on each of approximately 60 jumbo 150 mm x 15 mm Petri dishes with peptone-enriched (10 g l^-1^ peptone) NGM seeded with NA22. The cultures were never permitted to exhaust the food supply and were refed in the L4 stage with concentrated NA22 bacteria the day before harvesting (approximately the *E. coli* produced by a 200 ml LB liquid culture per Petri dish). After growing at 25°C for approximately 48 hours, day-1 adult hermaphrodites were washed free of *E. coli* using M9 and harvested in freezing buffer (50 mM HEPES, pH 7.5, 1 mM MgCl_2_, 100 mM KCl, 10% glycerol). The worm slurry was frozen dropwise in liquid nitrogen and stored in –80°C. Because lysates were prepared by pooling many individually grown cultures, reproducibility was assessed using three technical replicates in both the *fog-1(q253ts)* and *spe-9(hc88ts)* genetic backgrounds.

To extract proteins, the frozen worms were ground using a mortar and pestle with liquid nitrogen, and the frozen worm powder was collected in 50 ml tubes. 1 ml of lysis buffer [75 mM HEPES, pH 7.5, 1.5 mM MgCl_2_, 150 mM KCl, 15% glycerol, 0.0001% tergitol and cOmplete mini EDTA-free protease inhibitor (Roche, 1 tablet/ 5 ml)] was added to 1 g worm powder and subjected to sonication at 4°C with a Sonic Dismembrator Model 500 (Thermo Fisher Scientific Inc.) set at 30% amplitude using three cycles of 15 s pulses with 45 s intervals between pulses. The crude extract was centrifuged twice at 20,000 g for 10 min at 4°C. This low-speed supernatant was centrifuged at 100,000 g for 1 h at 4°C using an SW41 Ti rotor in a Beckman L-80 ultracentrifuge. This high-speed supernatant was collected, and 1 ml aliquots were flash-frozen on powdered dry ice and stored at –80°C.

#### Immunopurification and RNA extraction

For each genotype analyzed, three replicate immunopurifications were conducted, each starting with 160 mg of protein. For the SPN-4 immunopurifications, 720 µg of anti-FLAG monoclonal antibody M2 (F1804, Millipore Sigma) was cross-linked to 90 mg of Dynabeads protein G (Invitrogen) according to the manufacturer’s instructions. For the LIN-41 and OMA-1 immunopurifications, hybridoma cell lines producing anti-GFP monoclonal antibodies 12A6 and 4C9 (Sanchez et al., 2014) were obtained from the Developmental Studies Hybridoma Bank (University of Iowa). Hybridoma culture and antibody purification were as described (Tsukamoto et al., 2017). 360 µg of anti-GFP monoclonal antibody 12A6 and 360 µg of monoclonal anti-GFP antibody 4C9 were combined and crosslinked with 90 mg of Dynabeads protein G. Approximately 10 ml of high-speed supernatant (containing 160 mg of protein) was applied to a batch binding with antibody-conjugated Dynabeads for 1 h at 4°C. Basic IP buffer was 50 mM HEPES (pH7.5), 1 mM MgCl_2_, 100 mM KCl. After binding, unbound proteins were removed in the supernatant. The Dynabeads were washed three times with 10 ml of IP wash buffer [basic IP buffer with 300 mM KCl (final concentration), 0.05% NP-40, 5 mM 2-mercaptoethanol, 5 mM sodium citrate, 10 µM ZnCl_2_, cOmplete mini EDTA-free protease inhibitors (Roche, 1 tablet in 10 ml) and RNasin (Promega, 20 units ml^-1^)]. Bound RNA-binding proteins and their associated mRNAs were cleaved from the Dynabeads by adding and mixing with 4 ml of cleavage solution containing 225 units ml^-1^ of AcTEV protease (Life Technologies) for 18 h at 4°C. The TEV eluate was collected in the supernatant by placing tubes on DynaMag a magnet (Thermo Fisher Scientific, Inc.). 200 µl aliquots of the TEV eluate were dispensed into 20 individual 1.5 ml microcentrifuge tubes and mixed with 600 µl of TRIzol LS (Life Technologies) and flash-frozen on powered dry ice and stored at –80°C.

Twenty tubes containing 800 µl of the TEV eluate/TRIzol LS mixture were thawed at room temperature and 160 µl of chloroform was added to each tube and mixed vigorously for 15 s. After incubation at room temperature for 10 min, the mixture was centrifuged at 12,000 g for 15 min at 4°C. Supernatants were collected and applied onto two RNeasy MinElute spin columns using the RNeasy Micro Kit (74004, Qiagen). Finally, about 40 µl of RNA solution was collected from each immunopurification. RNA was quantified using Qubit fluorometric quantification (Thermo Fisher Scientific, Inc).

#### RNA sequencing library construction and RNA sequencing

RNA sequencing libraries were prepared using KAPA RNA HyperPrep Kit (KK8540-08098093702, Roche) according to the manufacturer’s instructions and barcoded during amplification. Library preparation utilized 2 ng of RNA for each of the SPN-4 immunopurifications, 10 ng of RNA for each of the LIN-41 immunopurifications, and 40 ng of RNA for each of the OMA-1 immunopurifications. Total RNA was also extracted from 200 µl of the high-speed supernatant of the lysates using RNeasy Micro Kit (74004, Qiagen) as a control. A Ribo-Zero rRNA Removal Kit (MRZH116, Illumina) was used to remove ribosomal RNA (rRNA) from the total RNA. Duplicate sequencing libraries were prepared from rRNA-depleted total RNA from the high-speed supernatant of the lysates using the same method so that enrichment over input could be calculated. The number of amplification cycles utilized was determined empirically using 1 µl of the sequencing library in a 10 µl test reaction containing a 1:10,000 dilution of SYBR Green I (Invitrogen), 2x KAPA HiFi HotStart ReadyMix (Roche) and custom barcoding primers. The empirically determined number of amplification cycles was used in a subsequent 50 µl reaction with the remaining eluate in the absence of SYBR Green using Kapa HiFi HotStart Readymix (Roche). Libraries were size-selected on a 2% E-Gel EX agarose gel (Invitrogen) with visualization using a UV-free Dark Reader transilluminator (Claire Chemical Research) and fragments between 250 bp and 400 bp were extracted using a MiniElute Gel Extraction Kit (Qiagen). Libraries were submitted to the University of Minnesota Genomics Center for 2x 36-bp sequencing using the Illumina NextSeq 500 to a depth of 37-55 million reads per sample. Sequencing data is available through the NCBI Gene Expression Omnibus (GEO) database under accession number GSE307638. Short read, raw sequencing files are available through the SRA Short Read Archives under accession number PRJNA1322436.

#### RNA mapping and Transcript Quantification

After trimming adapters with Trimmomatic (0.32), reads were mapped to the WBcel235/ce11 genome with STAR (v 2.5.3a) guided by gene annotations defined in Ensembl (release 91) and sorted and indexed with Samtools (v1.21). PCR duplicates were removed with Picard MarkDuplicates (v2.17.10). Gene-level abundance was estimated for Ensembl-defined annotations using the featureCounts function in the Bioconductor package Rsubread (v1.28.1). Principal component analysis and inspection of 5’-vs 3’-read coverage indicated that one IP sample from DG4400 (IP2) contained degraded RNA and was excluded from further analysis. An average of 10 million high-quality (MAPQ > 55) reads mapped to non-ribosomal genes within each sample. In prior work, we found that the sets of proteins and mRNAs associated with OMA-1 were not substantially different in the presence and absence of sperm (Spike et al., 2014b). This dataset also included LIN-41 and transcripts that associated with both OMA-1 and LIN-41. We also observed that SPN-4 expression in the most proximal oocytes was independent of the presence of sperm in the gonad (Tsukamoto et al., 2017), and that the same set of proteins copurify with SPN-4 in the presence and absence of sperm (unpublished results). Thus, we combined the data from the *fog-1(q252ts)* and *spe-9(hc88ts)* backgrounds to enhance the analytical power of the data (Table S1). A comparison of LIN-41- and OMA-1-associated mRNAs with our prior datasets (Tsukamoto et al., 2017) indicated replicability (Fig. S1). Differential gene expression of Ensembl-annotated genes was determined using DESeq2 (v1.26.0). Median ratio normalized counts were fit separately for each tagged protein to a regression model which included terms to account for differences between the *fog-1(q252ts)* and *spe-9(hc88ts)* backgrounds. A Welch-corrected t-test was used to identify transcripts over-represented in the IP samples relative to the lysates (altHypothesis=“greater”). *P* values were adjusted for multiple test correction using the Benjamini–Hochberg procedure. Transcripts enriched greater than 4-fold with adjusted *P*-values less than 0.05 were considered statistically significant.

#### Comparative analyses

To analyze the overlap between the sets of SPN-4-, LIN-41- and OMA-1-associated transcripts, the list of genes was filtered for a minimum expression level of log_2_(mean intensity) > -2.5 and culled of cTel and 21U gene entries to yield 16,917 genes. SPN-4 immunopurification replicates were averaged and plotted as an MA Plot in which Fold Change (IP over input) was tabulated over Expression Level [log_2_(mean intensity)] (Fig. 1D).

#### Upset plots

Upset plots were generated to illustrate the categorical overlap of genes that associate with SPN-4, LIN-41 and/or OMA-1 using the UpSetR Package (v. 1.4.0) (Fig. 1E).

#### Distance matrix

To assess the differences between RNA-binding protein enrichment values for SPN-4, LIN-41 and OMA-1, distance matrices on the set of 16,917 genes (see above) were calculated using the dist R function and method = “Euclidean” (Fig. S2A).

#### Heatmaps of dynamic genes

Heatmaps were produced to illustrate the patterns and clusters of transcripts as they enriched in the SPN-4, LIN-41 and OMA-1 immunopurification assays. Genes were selected based on a minimum expression level, association with at least one RNA-binding protein, and for minimum dynamics across the RNA-binding proteins. Specifically, genes were filtered for spn4_enrichment > -2, Log2_SPN.4_LYSATE_FPKM > 4, and stddev > 0.6. This yielded 2,480 genes. Heatmaps were generated using the pretty heatmap package (pheatmap v1.0.12) on enrichment values without z-scoring using options: clustering_distance_rows = “euclidean”, clustering_method = “complete” (Fig. S2B).

#### Gene ontology

GO categorical analysis was performed using clusterProfiler (v. 4.10.1) with settings: cutoff = 0.6, by = “p.adjust”, select_fun = min, and measure = “Wang” (Wu et al., 2021). Analyses were performed on each set of RNA-binding protein-associated genes (independent of their status for the other RNA-binding proteins). Biological Process categories are shown (Fig. S3C). Molecular Function and Cellular Component ontologies were also performed and are included in github materials. Individual categories, scores, and key genes driving each category are reported (Table S10).

#### scRNA-seq comparative analysis

To assess how the sets of SPN-4-, LIN-41- and/or OMA-1-associated transcripts behaved across the first five cell divisions of embryogenesis with single-cell resolution, we used a previously reported scRNA-seq dataset (Table 2 from (Tintori et al., 2016)). In that study, hand-dissected *C. elegans* blastomeres from 1-cell, 2-cell, 4-cell, 8-cell and 16-cell stage embryos were profiled by scRNA-seq methods with four to nine replicates per cell. To split the scRNA-seq dataset by behavior, we first filtered the transcriptome for expressed, dynamic genes. Specifically, we filtered for total RPKM > 5 and variance > 10 to yield 14,776 genes. To determine the clustering method that best split maternal mRNA genes from zygotically activated genes, we tested 10 clustering protocols. The clustering settings that yielded the greatest separation between maternal and zygotic gene clusters was: scale = “row”, clustering_distance_rows = “canberra”, clustering_method = “complete”, cutree_rows = 5. This split the data into 5 clusters, and SPN-4-, LIN-41- and OMA-1-associated RNA categories were annotated onto the resulting clustered heatmap (Fig. S3A). The same data could be displayed in alternative ways as a lineplot averaged across clusters and cells (Fig. S3B) or as lineplots split by RNA-binding protein category (Fig. 2 and S3E). The percentage of SPN-4-associated genes within each cluster was also recorded (Fig. S3C). To test whether the different SPN-4-, LIN-41- and/or OMA-1-associated gene categories were enriched in specific scRNA-seq expression clusters at frequencies greater than predicted by random chance, we performed Mosaic Analysis using the mosaic function in the package Visualizing Categorical Data (vcd, v. 1.4-13). In a mosaic plot, contingency tables are first generated. Next, categories are drawn as tiles from vertical splits based on one selection method (RNA-binding protein association) and horizontal splits based on a second selection method (scRNA-seq clusters) with the resulting areas of each tile proportional to the corresponding category size (Friendly, 1994). Deviation from independence was calculated as Pearson Residuals and layered onto the mosaic plot as residual-based coloration.

#### RNA motif analysis

To identify RNA sequence motifs that are over-represented in the 3’UTRs of SPN-4-associated mRNAs, we used XSTREME (v. 5.5.7), a Motif Discovery Enrichment Tool within the MEME Suite ecosystem (Grant and Bailey, 2021). Scrambled sequences were used for de novo motif discovery, and the union set of OMA-1/LIN-41-associated 3’UTR sequences (excluding SPN-4 binders) was used to find distinguishing motifs that differentiate SPN-4 association from OMA-1 or LIN-41 association. XSTREME was run for RNA motifs using the Ray 2013 RNA motif database (Ray et al., 2013) like so: --streme-totallength 4000000 --meme-searchsize 100000 --fdesc description --dna --dna2rna --evt 0.05 -- minw 6 --maxw 15 --align center --meme-mod zoops --sea-noseqs –m db/motif_databases/RNA/Ray2013_rbp_All_Species.meme --p SPN4_merged_IPd_3UTRs_clean_gt5.fa -n OMA1LIN41_IPd_3UTRs_clean_gt5.fa. Similar outputs were generated independent of the background set.

### smFISH

#### Embryo smFISH protocol

smFISH assays were performed using an adapted version of the Stellaris TurboFISH protocol as previously described (Parker et al., 2021; Tsanov et al., 2016). Custom smFISH probes for desired targets were designed using Stellaris RNA FISH Probe Designer (Biosearch Technologies; www.biosearchtech.com/stellaris-designer; version 4.2) and labeled with CalFluor 610 or Quasar 670 (Biosearch Technologies, Teddington, Middlesex, UK). Probesets used in this work are listed in Table S4.

Briefly, synchronized gravid animals were washed twice using M9 (3 g KH_2_PO_4_, 6 g Na_2_HPO_4_, 5 g NaCl, ddH_2_O to 1 liter) and bleached for embryos (40 ml ddH textsubscript2O, 7.2 ml 5M NaOH 4.5 ml 6% NaCl; 9 min total). Embryos were fixed and permeabilized by resuspension in 1 ml -20°C methanol and tubes were immediately freeze-cracked in liquid nitrogen 1 min, then incubated at -20°C for 5 min, then fixed in 1 ml -20°C acetone at -20°C for 5 min. After fixation, embryos were equilibrated with 1 ml of Stellaris Wash Buffer A solution (600 µl Stellaris Wash A buffer [SMF-WA1-60, Biosearch Technologies], 300 µl deionized formamide, 2.1 ml of DEPC-treated H_2_O), then resuspended in 100 µl of hybridization buffer solution with probes (99 µl Stellaris RNA FISH Hybridization Buffer [SMF-HB1-10, Biosearch Technologies], 11 µl deionized formamide, 2 µl of diluted smFISH probes) and kept at 37°C for 24-48 h in a thermoshaker until imaged. Final probe concentration during the hybridization step was 125 nM for the T30 poly(A) probe and 20 - 90 nM, optimized for each gene-specific probe.

Unbound probes were removed by washing twice with 1 ml of Stellaris Wash Buffer A solution at 37°C for 30 min each in the Thermomixer, with the second wash containing 2 µl of either 1 µg µl^-1^ DAPI or 5 mg ml^-1^ DAPI. One to two final wash(es) with Stellaris Wash Buffer B (Biosearch Technologies, SMF-WB1-20) was (were) carried out before resuspending in 20-50 µl of mounting media (2.5 ml 100% glycerol, 25 mg N-propyl gallate, 100 µl 1M Tris pH 8.0, 2.4 ml H_2_O) and sample was allowed to rest for 10-30 min at room temperature before slide preparation. The embryos were mounted on a slide using equal volumes of VectaShield antifade (H-1000, Vector Laboratories) and the resuspended sample.

#### smiFISH protocol

Some probes were generated using a smiFISH approach (the “i” stands for inexpensive). These probesets are listed in Table S4. smiFISH was performed using FLAPY primary probe extensions and secondary probes (Parker et al., 2021). Briefly, between 12 and 24 primary probes were designed using Oligostan (Tsanov et al., 2016) and ordered in a 25 nmol 96-well format from IDT Technologies and diluted to 100 µM in IDTE buffer (pH 8.0). Secondary FLAPY probes were ordered from Stellaris LGC with dual 5’- and 3’-fluorophore labeling using either Cal Fluor 610 or Quasar 670 (Biosearch Technologies, BNS-5082 and FC-1065, respectively). Individual probes were combined to a final concentration of 0.833 µM. 2 µl of these primary probe combination mixtures were mixed with 1 µl 50 µM FLAPY secondary probe, 1 µl NEB buffer 3 and 6 µl DEPC-treated H2O. The primary and secondary probe mixtures were incubated at 85°C for 3 min, 65°C for 3 min and 25°C for 5 min to anneal. 2 µl of annealed probe mixtures were used in place of the smFISH probeset and all other steps of the protocol were conducted above.

#### Gonad smFISH and WGA staining protocol

Gonads were dissected from adult hermaphrodites in less than 10 min in a glass depression well in PBT [RNase-free 1x PBS, pH 7.4 (Invitrogen) + 1% Tween (Thermo Fisher Scientific Inc.)], and fixed for 25 min in PBT + 3.7% formaldehyde (Electron Microscopy Sciences). Fixed gonads were washed with PBT and then permeabilized. Sample permeabilization followed the method described by (Lee et al., 2016) for Fig. 8: PBT+0.1% Triton-X100 (Thermo Fisher Scientific Inc.) for 10 min at room temperature followed by a PBT wash and a subsequent incubation in 1 ml 70% ice-cold ethanol for 16+ h at 4°C. Sample permeabilization was a variation on the method described by (Yoon et al., 2016) for Fig. S12: 0.5 ml 100% ice-cold methanol for 1+ h at -20°C followed by gradual rehydration using PBT. Prior to adding ethanol (Fig. 8) or methanol (Fig. S12) the gonads were transferred to a 1.5-ml non-stick RNase-free microfuge tube (Invitrogen). After this transfer and prior to subsequent buffer changes, the gonads were spun for 2 min at 500 g to permit removal of supernatant. Permeabilized gonads were equilibrated with Stellaris wash buffer A solution and resuspended in Stellaris RNA fish hybridization buffer with probes as described above for embryos. Final *chs-1* probe concentration was 90 nM. Post-hybridization washes for Fig. 8 and sample mounting were as described above for embryos. The wheat germ agglutinin (WGA) staining shown in Fig. S12 was performed by adding PBT wash steps, both before and after a 10 min room-temperature incubation with 1-5 µg ml^-1^ 488-WGA (Thermo Fisher Scientific Inc.), after the smFISH washes were completed.

### Imaging and smFISH spot counting

#### DeltaVision image capture, processing, and analysis

Fluorescence microscopic images of smFISH samples that form the basis for the results shown in Fig. 3-6 and Fig. S5-S7 were acquired on a DeltaVision Elite inverted microscope (GE Healthcare), using a Photometrics Cool Snap HQ2 camera and an Olympus PLAN APO 60× (1.42 NA, PLAPON60XOSC2) objective, an Insight SSI 7-Color Solid State Light Engine and SoftWorx software (Applied Precision). Single-color capture was achieved using a beam Splitter suitable for DAPI, FITC, Alexa 594, CY5 and GFP/mCherry and filters with the following single-pass emission settings (DAPI = 435/48, CFP = 475/24, GFP/FITC = 525/48, YFP = 548/22, TRITC = 597/45, mCherry/Alexa594 = 625/45, Cy5 = 679/34). Images were collected using DataVision SoftWorx with identical exposures across a given dataset. Images were collected every 0.2 µm in the z-direction. Representative images were deconvolved using DataVision SoftWorx image analysis software and further processed using FIJI. A minimum of 7 embryos were imaged for each genetic condition and cell stage. Image analysis was performed on images prior to deconvolution.

Deltavision mRNA spot detection image analysis for Fig. 3 and S5 was performed using fish-quant (Mueller et al., 2013). Embryos were first manually outlined, then 3D LoG filtered using default FISH-quant parameters (size=5, s.d.=1). Spots were pre-detected using a local maximum fitting. mRNA spots were detected using a manually determined image-dependent intensity and quality threshold, with the following options: lambda_EM=610, lambda_Ex=590, NA=1.42, RI=1.518. Post-processing statistics and plotting were generated using customized R scripts that are included on the GitHub repository associated with this paper. Deltavision smFISH spots detection image analysis for Fig. 4, 5, 6, S6 and S7 was conducted on images prior to deconvolution using an in-house Python pipeline for embryo segmentation and spot detection. Embryos were first segmented using in-house, machine learning modifications to cellpose using 2D image inputs of the brightfield channel to determine embryo boundaries and cell stage (Pachitariu and Stringer, 2022; Stringer et al., 2021). Inaccuracies in stage assessment were either corrected or removed. Spot detection using bigFish (from of fish-quant v2) (Imbert et al., 2022) was restricted to the embryo regions of the field of vision. mRNA spots were identified in 3D using voxel size of 1448, 450, 450 nm and spot radius of 1409, 340, 340 and 1283, 310, 310 nm for Quasar 670 and Cal Fluor Red 610 channels, respectively. The voxel size and spot radii were optimized for accuracy and consistency. Post-processing statistics and plotting methods are included as customized R scripts in the associated GitHub repository.

#### Eclipse Ni-E image capture, processing and analysis

Images that form the basis for the smFISH results shown in Fig. 8, 10, S4B, S12 and S13 were acquired using a Plan Apo l 100x (numerical aperture 1.45) objective on an Eclipse Ni-E microscope (Nikon Inc) equipped with a motorized ProScan H101E1F XY stage (Prior Scientific), SOLA light engine (Lumencor) and an ORCA-Fusion C14440 digital camera (Hamamatsu). Images shown in Fig. 8A-D used a 60x Plan Apo (numerical aperture 1.40) objective on the same microscope. Filter sets 49009 ET-CY5 NX and 49306 ET-RED#1 FISH (Chroma) were used to discriminate between *set-3* (Quasar 670) and *chs-1* or *lin-41* (Calfluor Red 610) smFISH signals, respectively. Identical exposure settings were used to image each smFISH channel in the different stages and genotypes with images collected every 0.25 µm in the Z dimension. Z-stacks were deconvolved with version 6.02 of NIS-elements (Nikon Inc.) using either the fast deconvolution method with a custom noise level of 42 (images in Fig. 8F,G, S12A,B) or the Richardson-Lucy deconvolution method at medium noise level with auto stopping and a microscope-specific point spread function (images in Fig. 10). Embryo smFISH images shown in Fig. 10 are maximum-intensity projections created from full-depth Z-stacks. Fig. 8F,G and S12A,B are maximum intensity projections of a 2.5 µm Z-stack collected near the vertical mid-point of the oocyte. Image analysis was performed prior to deconvolution using the inhouse Python pipeline described above. A small number of images (>3%) with obvious segmentation or spot identification problems were redacted, as described in Table S9. Images were analyzed using a specified voxel size of 65×65×250 nanometers and spot sizes of 110×110×300 and 97×97×300 nanometers for the Quasar 670 and Cal Fluor Red 610 channels, respectively.

#### Measuring transcript abundance in gonad smFISH samples

*chs-1* transcript abundance in oocytes and newly fertilized embryos was determined using the 3D-measurement module in version 6.02 of NIS-elements (Nikon Inc.). Images were sharpened with a Gauss-Laplace transformation (setting 1.7) and individual smFISH transcripts, defined as 0.35-micron spots with a 50-fold increase in intensity, were identified using the “clustered spots” option. Oocytes and embryos were manually segmented and the number of spots in the segmented volume counted. We compared this method with the 3D spot detection pipeline used to analyze early embryos and found strong positive correlations between the two methods. At the 2-cell stage, the method used to count transcripts in oocytes and newly fertilized embryos identified, on average, 73-74% of the *chs-1* transcripts in both wild-type and LET-711-depleted embryos. LET-711-depleted embryos were analyzed because, like oocytes, they often contain > 10,000 *chs-1* transcripts. Pearson’s correlation coefficients for each comparison were *r* = 0.977, df = 36, *P* > 0.001 for wild type and *r* = 0.838, df = 34, *P* > 0.001 for LET-711-depleted embryos.

### SPN-4 RRM binding to RNA *in vitro*

#### SPN-4 RRM expression in *E. coli*

The expression plasmid for production of the SPN-4 RRM (Met1–Lys200) in *E. coli*, pECT30 (see Table S3 for DNA sequence), was constructed as shown in Fig. S14A. pECT30, and all plasmid intermediates, were validated by Sanger sequencing (Azenta Genewiz). The oligonucleotide primers used for construction are listed in Table S3. A *spn-4* cDNA (ET22), codon optimized for *E. coli*, was commercially synthesized using gBlock Hi-Fi (IDT) and used as a template for PCR to amplify DNA fragments encoding TEV::3xFLAG::SPN-4 RRM(Met1-Lys200) using the ET47/ET48 primer pair and Q5 DNA polymerase (New England Biolabs). The PCR product was digested with EcoRI and HindIII and gel-purified using the QIAquick gel extraction kit. (QIAGEN). The purified DNA fragment was ligated into the pTT314 plasmid between EcoRI and HindIII sites using T4 DNA ligase (New England Biolabs) to generate the pECT25 intermediate. Empirically, we found that moving the 6xHis tag to the C-terminus improved protein stability. To move the 6xHis tag from the N-terminus to the C-terminus, an *MBP::TEV::3xFLAG::spn-4 RRM::6xHis* fragment was amplified from pECT25 using the primer pair ET50/ET25 and digested with NcoI and HindIII. The digested PCR fragment was purified and inserted into pECT25 between NcoI and HindIII sites to generate pECT30.

#### Protein induction and purification

pECT30 was transformed into BL21-AI *E. coli* (Promega). The cells were grown at 30°C in LB medium containing 50 µg ml^-1^ of kanamycin until an OD_600_ of 0.4-0.6. MBP::TEV::3xFLAG::SPN-4 RRM::6xHis was induced for 4 h with 1 mM isopropyl-*β*-D-1-thiogalactopyranoside and 0.2% L-arabinose. Cells were harvested by centrifugation at 3500 g for 20 min at 4°C, and proteins were extracted using Bacterial Protein Extraction Reagent (B-PER; Thermo Fisher Scientific) containing EDTA-free Halt protease inhibitors (Thermo Fisher Scientific), 5 units ml^-1^ of DNase I (Thermo Fisher Scientific), and 100 µg ml^-1^ of lysozyme (Thermo Fisher Scientific). The cell lysate was centrifuged at 20,000 g for 20 min at 4°C, and the supernatant was used for protein purification. A colloidal Coomassie-stained 4-12% NuPAGE gel showing the purification steps is shown (Fig. S14B).

For the first step, 1 mM phenylmethylsulfonyl fluoride (PMSF) was added to the supernatant, and the supernatant was applied to a DEAE Sepharose Fast Flow (Cytiva) column at 4°C. The flowthrough was collected and washed with 1 column volume (cv) of DEAE wash buffer [25 mM Tris-HCl (pH 8.0), 50 mM NaCl, 1 mM EDTA, 1 mM DTT]. The flowthrough and the wash were combined. For the second step, 1 mM PMSF was added and applied to Amylose resin (New England Biolabs) at 4°C. After binding, the resin was washed with 12 cv of binding buffer [20 mM Tris-HCl (pH 7.4), 200 mM NaCl, 1 mM EDTA]. The bound proteins were eluted with 2 cv of elution buffer [20 mM Tris-HCl (pH 7.4), 200 mM NaCl, 1 mM EDTA, 10 mM maltose]. MBP::TEV::3xFLAG::SPN-4 RRM::6xHis was identified by electrophoresis on a 4-12% Bis-Tris NuPAGE gel (Invitrogen) stained with the Colloidal Blue staining kit (Invitrogen). The peak fractions were combined and dialyzed overnight at 4°C against dialysis buffer [10% glycerol, 25 mM NaCl, 50 mM Tris-HCl (pH 8.0)]. The dialyzed sample was digested with 0.4-1 µM of MBP::TEV protease (a kind gift of Mark Zweifel, University of Minnesota) overnight at 4°C. In the third step, the TEV protease-digested sample was applied to a Ni-NTA agarose (GoldBio) column at 4°C. After binding, the column was washed with 20 cv of wash buffer [50 mM NaH_2_PO_4_, 300 mM NaCl, 10 mM imidazole (pH 8.0)]. Bound proteins were eluted with 3 cv of elution buffer [50 mM NaH_2_PO_4_, 300 mM NaCl, 250 mM imidazole (pH 8.0)]. Peak fractions were identified as above. In the fourth step, the peak fractions were pooled and applied to an Amylose resin column. The flowthrough was collected and washed with 1 cv of 20 mM Tris-HCl (pH 7.4), 200 mM NaCl, 1 mM EDTA. The flowthrough and the wash were combined and dialyzed for 4 h at 4°C against 10% glycerol, 25 mM NaCl, 50 mM Tris-HCl (pH 8.0), 1 mM DTT. The purified 3xFLAG::SPN-4 RRM::6xHis was approximately 90% pure and stored at -80°C. (Fig. S14B, lines 13-15).

#### RNA binding by EMSA

HPLC-purified RNA oligonucleotides (Table S3) were commercially synthesized with a 5’Cy5 label (IDT). 50 nM to 2.9 µM of purified SPN-4 RRM was incubated with the 32 nM of labeled RNA in binding buffer [10 mM Tris-HCl (pH 7.5), 150 mM NaCl, 0.001% NP-40, 10% glycerol, 0.01 mg ml^-1^ tRNA, 0.025% Tween-20] for 15 min at room temperature. A longer incubation time (30 min) did not alter the fraction bound. The samples were run on an 8% TBE gel (Invitrogen) in 1x TBE buffer (89 mM Tris, 89 mM boric acid, 2 mM EDTA) at 100 volts, room temperature, for 45-60 min depending on the length of the RNA. The gels were imaged using a LI-COR Odyssey Fc with the 700 nm channel and quantified using ImageStudio (LI-COR). The fraction of bound RNA versus the protein concentration was plotted using labplot2 (v2.11.1) and fitted to Eqn 1 to determine the equilibrium dissociation constant (K_d_):

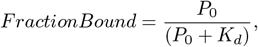

where *P*_*0*_ is the protein concentration and *K*_*d*_ is the dissociation constant. For all binding plots shown, the goodness of fit as measured by R^2^ was 0.963-0.979.

### Artificial intelligence (AI) statement

During the course of this work, the authors utilized AI to: (1) debug and streamline computer code for analysis of smFISH data; (2) inform and fact check statistical analyses performed using R; and (3) complement the use of PubMed for identifying published work potentially informative for the analyses herein.

### Accessibility statement

Where possible, we have tried to make colors in microscopy images and plots discernible to people with colorblindness or reduced color perception. We opted for magenta/green combinations in microscopy images and the viridis and tol packages in R.

## BACK MATTER

## Acknowledgements

We are grateful to Mark Zweifel for advice on protein purification and quantitation of RNA-protein interactions. Katherine Walstrom provided advice and guidance on molecular modeling. We thank Tejiri Agbamu for technical assistance with genome editing. This work utilized resources from the University of Colorado Boulder Research Computing Group, which is supported by the National Science Foundation (awards ACI-1532235 and ACI-1532236), the University of Colorado Boulder and Colorado State University. This work was completed in part using resources provided by the University of Minnesota Genomics Center (RRID: SCR_012413) and the University of Minnesota Supercomputing Institute (MSI). This work utilized microscopy resources from NIH grant 1S10 OD025127 and support from the CSU Microscope Imaging Network. Some strains were provided by the Caenorhabditis Genetics Center, which is funded by National Institutes of Health Office of Research Infrastructure Programs (P40 OD010440). Some figure elements were created in BioRender.

## Contribution

Conceptualization: C.A.S., D.M.P., T.,T., N.T., E.C.T., D.G., E.O.N.; Methodology, formal analysis, and investigation: C.A.S, D.M.P., T.T., N.T., E.C.T., K.C., M.D.G., D.G., E.O.N.; Writing, review, and editing: C.A.S, D.M.P., T.T., N.T., E.C.T., K.C., M.D.G., D.G., E.O.N.; Supervision, project administration and funding acquisition: D.G., E.O.N.

## Funding

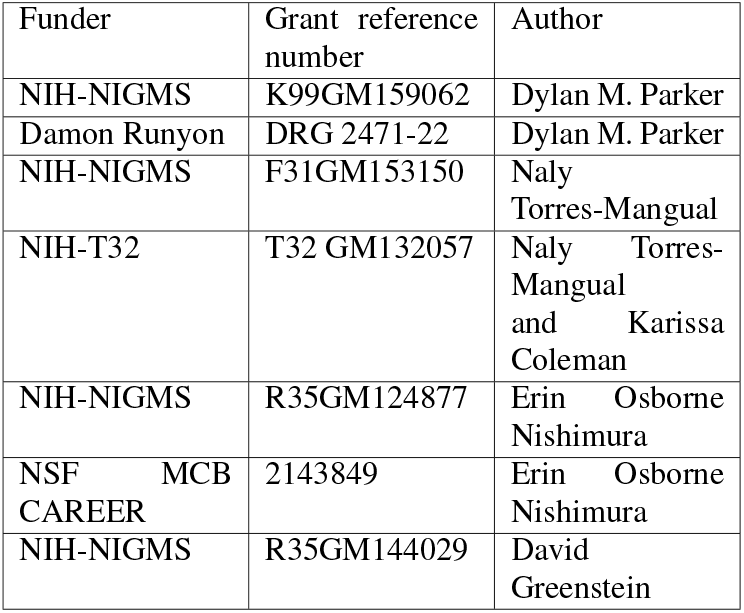

| Funder | Grant reference number | Author |
| --- | --- | --- |
| NIH-NIGMS | K99GM159062 | Dylan M. Parker |
| Damon Runyon | DRG 2471-22 | Dylan M. Parker |
| NIH-NIGMS | F31GM153150 | Naly Torres-Mangual |
| NIH-T32 | T32 GM132057 | Naly Torres-Mangual and Karissa Coleman |
| NIH-NIGMS | R35GM124877 | Erin Osborne Nishimura |
| NSF MCB CAREER | 2143849 | Erin Osborne Nishimura |
| NIH-NIGMS | R35GM144029 | David Greenstein |

## Data availability

Strains and plasmids and detailed protocols are available upon request. All scripts, code, and analytical methods are available on this associated GitHub repository: https://github.com/erinosb/SPN4_maternal_mRNA. Sequencing data is available through NCBI Gene Expression Omnibus (GEO) database under accession number GSE307638 and accessible at https://www.ncbi.nlm.nih.gov/geo/query/acc.cgi?acc=GSE307638. Raw sequencing files are available at SRA (Short Read Archive) under Project PRJNA1322436 at https://www.ncbi.nlm.nih.gov/Traces/study/?acc=PRJNA1322436&o=acc_s%3Aa. Raw images will be made available at Dryad.

## Supplementary

Supplementary information follows or will be available upon publication.

