## SupplementalInfo for "The SPN-4 Rbfox RNA-binding protein selects maternal mRNAs for CCR4-NOT-dependent clearance in early *Caenorhabditis elegans* embryos"

<u>Supplement:</u>

10 Supplemental Tables

14 Supplemental Figures

### Supplemental Tables

**Table S1. LIN-41-, OMA-1 and SPN-4-associated transcripts identified by immunopurification of RNA-binding proteins and RNA sequencing. (Related to Fig. 1).**

**Table S2 *C. elegans* strains used for this study**

| Strain | Genotype |
|---|---|
| N2 | Wild type, Bristol isolate |
| EV960 | *ccr-4(tm1312)* IV/*nT1[qIs51]* IV;V |
| FX30234 | *tmC9[F36H1.2(tmIs1221)]* IV |
| JK5996 | *puf-11(q971)* IV |
| JK6321 | *puf-11(q971) puf-3(q966)* IV*/nT1[qIs51]* IV; V |
| DG2566 | *fog-1(q253*ts*)* I; *oma-1(zu405te33)* IV; *tnIs17[pCS410 oma-1p::oma-1::s-tag::tev::gfp, unc-119(+)]* |
| DG2581 | *spe-9(hc88*ts*)* I; *oma-1(zu405te33)* IV; *tnIs17[pCS410 oma-1p::oma-1::s-tag::tev::gfp, unc-119(+)]* |
| DG3913 | *lin-41(tn1541[gfp::stag::lin-41])* I |
| DG3923 | *fog-1(q253*ts*) lin-41(tn1541[gfp::tev::s-tag::lin-41])* I |
| DG4158 | *spn-4(tn1699[spn-4::gfp::3xflag])* V |
| DG4398 | *fog-1(q253*ts*)* I; *spn-4(tn1699[spn-4::gfp::tev::3xflag])* V |
| DG4400 | *spe-9(hc88*ts*)* I; *spn-4(tn1699[spn-4::gfp::tev::3xflag])* V |
| DG4485 | *spe-9(hc88*ts*) lin-41(tn1541[gfp::tev::s-tag::lin-41])* I |
| DG5256 | *spn-4(tm291)/ tmC3[egl-9(tmIs1230) spn-4(tn2056[spn-4::gfp::3xflag])]* V |
| DG5324 | *puf-11(q971) tmC9[F36H1.2(tmIs1221)]* IV |
| DG5354 | *lin-41(tn1541[gfp::stag::lin-41] tn2074[lgPolyCΔ in 3'UTR]* I |
| DG5364 | *puf-11(q971) puf-3(q966)/puf-11(q971) tmC9[F36H1.2(tmIs1221)]* IV; *spn-4(tn1699[spn-4::gfp::3xflag])* V |
| DG5365 | *puf-11(q971) puf-3(q966)/puf-11(q971) tmC9[F36H1.2(tmIs1221)]* IV |
| DG5398 | *lin-41(tn1541[gfp::stag::lin-41] tn2078[medFoxΔ in 3'UTR]* I |
| DG5399 | *lin-41(tn1541[gfp::stag::lin-41] tn2079[lgFoxΔ in 3'UTR]* I |
| DG5410 | *lin-41(tn1541[gfp::stag::lin-41] tn2084[smPolyCΔ in 3'UTR]* I |
| DG5440 | *puf-11(q971) puf-3(q966)/puf-11(q971) tmC9[F36H1.2(tmIs1221)]* IV; *spn-4(tm291)/tmC3[egl-9(tmIs1230)]* V |
| DG5444 | *let-711(tn2102)/+* III; *puf-11(q971) puf-3(q966)* IV [a] |
| DG5454 | *puf-11(q971) puf-3(q966)/puf-11(q971) tmC9[F36H1.2(tmIs1221)]* IV; *+/tmC3[egl-9(tmIs1230)]* V |
| DG5458 | *let-711(tn2109)/+* III; *puf-11(q971) puf-3(q966)* IV [a] |
| DG5459 | *ccf-1(tn2110)/+* III; *puf-11(q971) puf-3(q966)* IV [a] |
| DG5464 | *puf-11(q971) puf-3(q966)/puf-11(q971) tmC9[F36H1.2(tmIs1221)]* IV; *spn-4(or191*ts*)/tmC3[egl-9(tmIs1230)]* V |
| DG5487 | *let-711(tn2087)/+* III; *puf-11(q971) puf-3(q966)* IV [b] |
| DG5491 | *let-711(tn2105)/+* III; *puf-11(q971) puf-3(q966)* IV [c] |
| DG5492 | *let-711(tn2106)/+* III; *puf-11(q971) puf-3(q966)* IV [c] |
| DG5493 | *let-711(tn2108)/+* III; *puf-11(q971) puf-3(q966)* IV [c] |
| DG5517 | *spn-4(tn2091)/+* V; *puf-11(q971) puf-3(q966)* IV [c] |
| DG5530 | *let-711(tn2088)* III; *puf-11(q971) puf-3(q966)* IV [b] |
| DG5533 | *let-711(tn2082)/+* III; *puf-11(q971) puf-3(q966)* IV [d] |
| DG5538 | *puf-11(q971) puf-3(q966)/puf-11(q971) tmC9[F36H1.2(tmIs1221)]* IV; *spn-4(tn2091)/tmC3[egl-9(tmIs1230)]* V |
| DG5632 | *let-711(tn2130[let-711::gfp::aid::3xflag])* III |
| DG5634 | *ccf-1(tn2132[ccf-1::gfp::aid::3xflag])* III |
| DG5671 | *+/hT2[umnIs60(myo-2p::mKate2)]* I; *+/hT2[bli-4(e937)]* III*; puf-11(q971) puf-3(q966)/puf-11(q971) tmC9[tmIs1221]* IV |

| | |
|---|---|
| DG5672 | *+/hT2[umnIs60(myo-2p::mKate2)]* I; *let-711(tn2135)/hT2[bli-4(e937)]* III; *puf-11(q971) puf-3(q966)/puf-11(q971) tmC9[F36H1.2(tmIs1221)]* IV |
| DG5673 | *+/hT2[umnIs60(myo-2p::mKate2)]* I; *let-711(tn2136)/hT2[bli-4(e937)]* III; *puf-11(q971) puf-3(q966)/puf-11(q971) tmC9[F36H1.2(tmIs1221)]* IV |
| DG5674 | *+/hT2[umnIs60(myo-2p::mKate2)]* I; *let-711(tn2137)/hT2[bli-4(e937)]* III; *puf-11(q971) puf-3(q966)/puf-11(q971) tmC9[F36H1.2(tmIs1221)]* IV |
| DG5675 | *+/hT2[umnIs60(myo-2p::mKate2)]* I; *let-711(tn2138)/hT2[bli-4(e937)]* III; *puf-11(q971) puf-3(q966)/puf-11(q971) tmC9[F36H1.2(tmIs1221)]* IV |
| DG5680 | *wrdSi18[mex-5p::TIR1::F2A::mTagBFP2::tbb-2 3'UTR]* I*; ccf-1(tn2132[ccf-1::gfp::aid::3xflag])* III |
| DG5682 | *wrdSi18[mex-5p::TIR1::F2A::mTagBFP2::tbb-2 3'UTR]* I; *let-711(tn2130[let-711::gfp::aid::3xflag])* III |
| DG5701 | *+/hT2[umnIs60(myo-2p::mKate2)]* I; *let-711(tn2135)/hT2[bli-4(e937)]* III |
| DG5702 | *+/hT2[umnIs60(myo-2p::mKate2)]* I; *let-711(tn2136)/hT2[bli-4(e937)]* III |
| DG5703 | *+/hT2[umnIs60(myo-2p::mKate2)]* I; *let-711(tn2137)/hT2[bli-4(e937)]* III |
| DG5704 | *+/hT2[umnIs60(myo-2p::mKate2)]* I; *let-711(tn2138)/hT2[bli-4(e937)]* III |
| DG5713 | *ccf-1(gk40)/tmC29[unc-49(tmIs1259)]* III |
| DG5728 | *ccf-1(tn2166)/tmC29[unc-49(tmIs1259)]* III |
| DG5730 | *ccf-1(tn2167)/tmC29[unc-49(tmIs1259)]* III |
| DG5732 | *ccf-1(tn2168)* III |
| DG5751 | *+/hT2[umnIs60(myo-2p::mKate2)]* I; *ccf-1(tn2166)/hT2[bli-4(e937)]* III; *puf-11(q971) puf-3(q966)/puf-11(q971) tmC9[F36H1.2(tmIs1221)]* IV |
| DG5752 | *+/hT2[umnIs60(myo-2p::mKate2)]* I; *ccf-1(tn2167)/hT2[bli-4(e937)]* III*; puf-11(q971) puf-3(q966)/puf-11(q971) tmC9[F36H1.2(tmIs1221)]* IV |
| DG5754 | *ccf-1(tn2168)* III; *puf-11(q971) puf-3(q966)/puf-11(q971) tmC9[F36H1.2(tmIs1221)]* IV |
| DG5755 | *+/hT2[umnIs60(myo-2p::mKate2)]* I; *ccf-1(gk40)/hT2[bli-4(e937)]* III; *puf-11(q971) puf-3(q966)/puf-11(q971) tmC9[F36H1.2(tmIs1221)]* IV |
| DG5779 | *lin-41(tn1541[gfp::stag::lin-41] tn2200 [smFox∆ in 3'UTR*] I |
| DG5919 | *lin-41(tn2238[smFox∆ in 3'UTR])* I |
| DG5965 | *lin-41(tn2238[smFox∆ in 3'UTR])* I; *spn-4(tm291)/tmC3[egl-9(tmIs1230) spn-4(tn2056[spn-4::gfp::3xflag])]* V |
| DG5972 | *chs-1(tn2255[Fox∆ in 3'UTR])* I |
| DG5974 | *ccr-4(tm1312)/tmC9[F36H1.2(tmIs1221)]* IV |
| DG6008 | *ccf-1(tn2168)* III*; ccr-4(tm1312)/tmC9[F36H1.2(tmIs1221)]* IV |

[a] EMS suppressor strain used for WGS; 1x backcross to DG5365.
[b] EMS suppressor strain used for WGS; 3x backcross to DG5365.
[c] EMS suppressor strain used for WGS; 2x backcross to DG5365.
[d] EMS suppressor strain used for WGS; 5x backcross to DG5365.

**Table S3. Sequence of oligonucleotides used in this study for genome editing, PCR, sequencing and plasmid constructions.**

**Table S4. Probesets used for smFISH and smiFISH.**

**Table S5. Transcript counts for comparing the wild-type and *spn-4(tm291)* mutants—raw values and statistics. (Related to Fig. 3 and S5).**

**Table S6. Transcript counts for analyzing *lin-41* 3'UTR deletions—raw values and statistics. (Related to Fig. 4 and S6 and).**

**Table S7. Transcript counts for analyzing the genetic interaction between a *lin-41* Rbfox-motif deletion and *spn-4* mutants —raw values and statistics. (Related to Fig. 5 and S7).**

**Table S8. Transcript counts for analyzing the Rbfox-motif deletion in the *chs-1* 3'UTR deletion—raw values and statistics. (Related to Fig. 6).**

**Table S9. Transcript counts for analyzing the requirement of LET-711 and CCF-1 for SPN-4-dependent maternal mRNA clearance—raw values and statistics. (Related to Fig. 10, S4 and S13).**

**Table S10. GO Ontology output. (Related to Fig. S2).**

#### Supplemental Figures

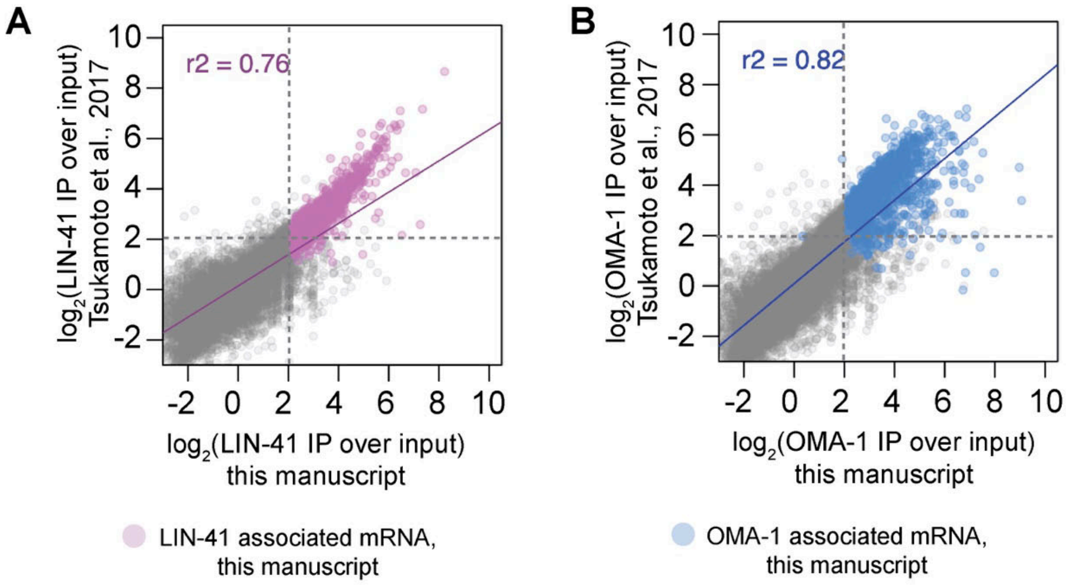


**Fig. S1. A comparison of LIN-41- and OMA-1-associated transcripts identified in this study to those identified by Tsukamoto et al. (2017).** (Related to Fig. 1 and Table S1). In this study, LIN-41- and OMA-1-associated mRNAs were sequenced using a low-input RNA-seq library whereas Tsukamoto et al. (2017) prepared sequencing libraries directly from the recovered mRNA (see Fig. 1 in the main text) (Tsukamoto et al., 2017) (A, B) Scatter plots comparing the $\log_2$ values for immunopurified RNA over input for Tsukamoto et al. (2017) on the y-axis to results from this study (x-axis) for LIN-41 (A) and OMA-1 (B). Pearson correlation was calculated to include all points ("everything") using the cor function in R stats.

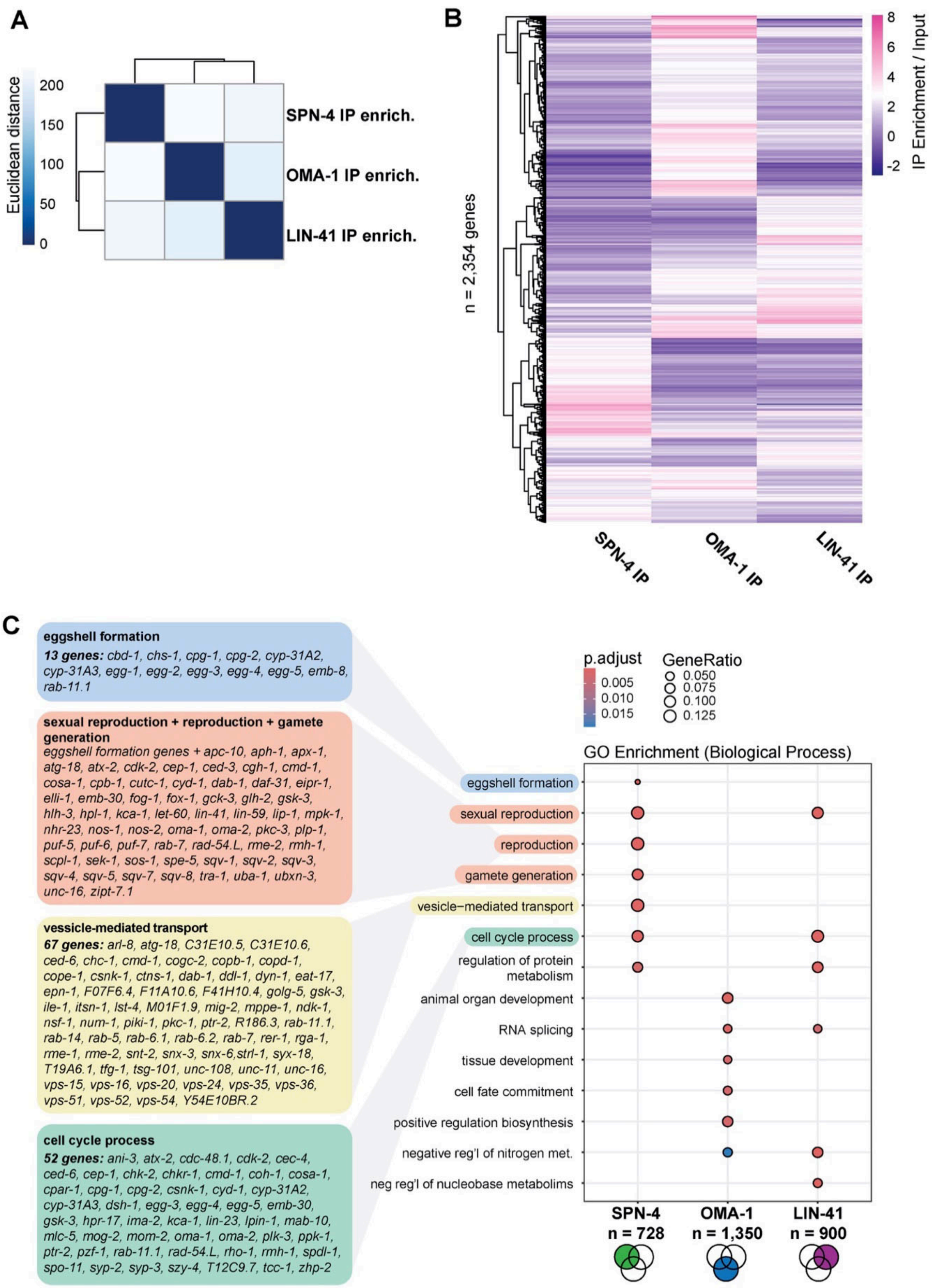


**Fig. S2. Characteristics of SPN-4-, OMA-1- and LIN-41-associated transcripts.** (Related to Fig. 1 and Table S1). (A) This plot illustrates the relationship between mRNAs captured by the three different immunopurification assays. Transcripts were filtered for a minimum expression level (mean $\log_2$(-2.5)) and culled of cTel and 21U gene entries to yield 16,917 genes. Euclidean distance was calculated, clustered and plotted. Darker shading represents a more similar relationship. (B)

Transcripts associated with at least one of the three RNA-binding proteins were plotted as a heatmap. Distance metrics were calculated using the Euclidean method and were clustered using the complete method, without applying z-scoring. (C) Gene Ontology (GO) terms enriched in the sets of SPN-4-, LIN-41- or OMA-1-associated transcripts are tabulated as dotplots in which terms shared across gene sets are comparable across a given row. The exact genes driving each GO term are listed and color-matched to their respective terms.

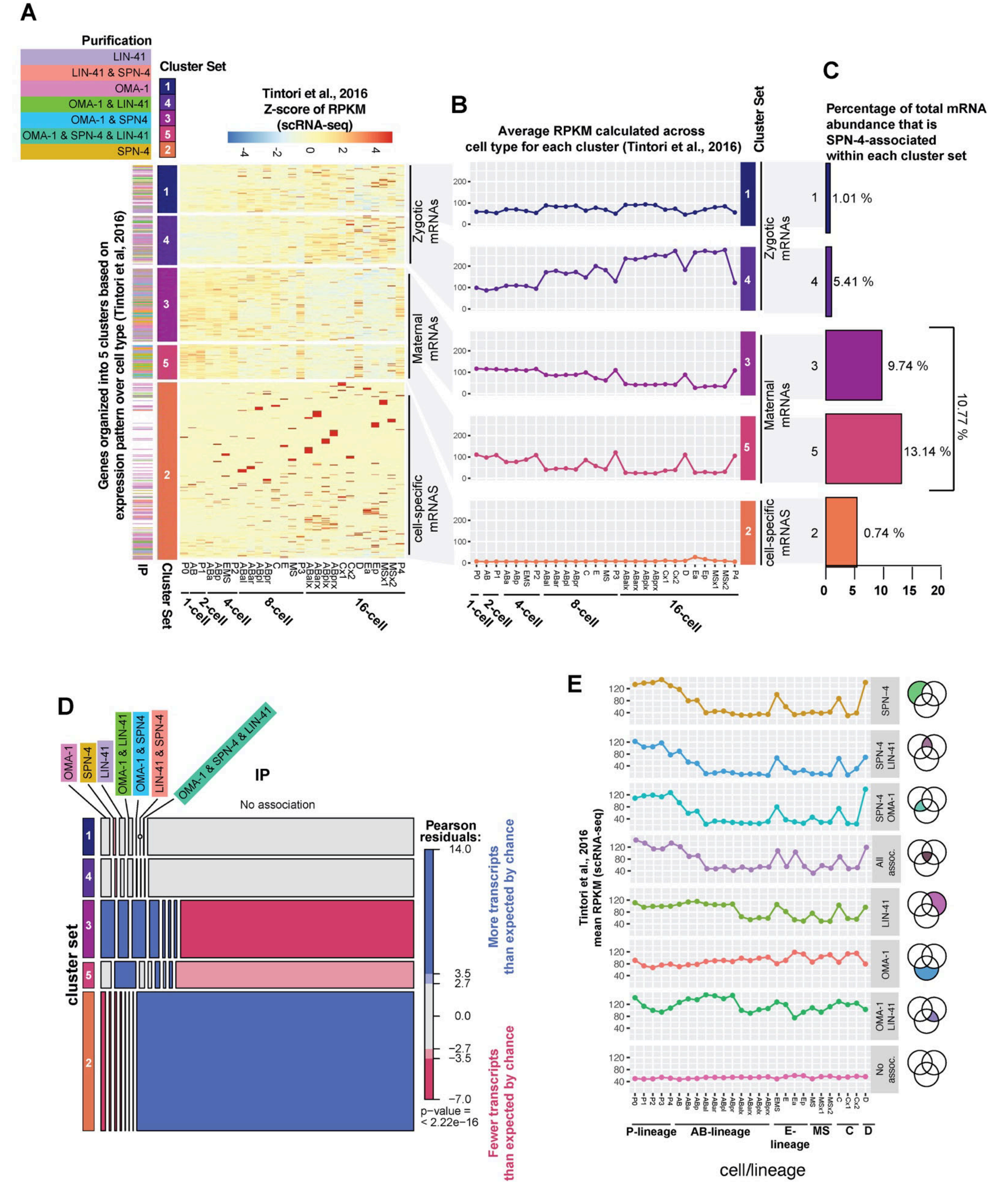


**Fig. S3. Single-cell RNA-seq transcriptome profiles from the first 31 blastomeres.** (Related to Fig. 2). (A) scRNA-seq transcriptome profiles from the first 31 embryonic blastomeres were reported in Tintori et al., (2016) (Tintori et al., 2016). That data is depicted here in heatmap form. Genes were filtered for total RPKM > 5 and variance > 10 to yield 14,776 genes. Heatmaps were generated with Canberra distance, complete clustering, row scaling, and cutree = 5. SPN-4-, OMA-1- or LIN-41-associated mRNAs were annotated in the specified colors. Two clusters represented newly transcribed zygotic mRNAs (Cluster Sets 1 and 4), two represented maternal mRNAs that undergo decay (Cluster Sets 3 and 5), and one was comprised of zygotic mRNAs transcribed with cell-type specificity (Cluster Set 2). (B) Genes were clustered based on categories generated in hierarchical clustering in (A). Mean expression for genes across each cluster was calculated for each cell type and plotted as line plots showing mean(RPKM) over cell type. Cluster sets differentiate maternal mRNA from zygotic mRNA and from cell-specific zygotic expression patterns. (C) The percentage of genes within each cluster set [generated in (A)] that have transcripts associated with SPN-4. (D) Contingency tables were generated from frequencies of SPN-4, OMA-1, and LIN-41 association and cluster identity. Contingency tables are shown plotted as Mosaic plots in which the area is proportional to frequencies. Mosaic plots are representations of contingency tables, which relate RNA-binding protein association to the membership in the various scRNA-seq clusters. Deviations between observed and expected numbers were quantified as Pearson residuals are shown, in which high residuals represent a larger observed membership than expected by chance and low residuals represent a smaller one. This illustrated that all RNA-binding protein-associated cohorts were over-represented in Cluster Set 3 and SPN-4-associated cohorts were over-represented in Cluster Set 5. Shading illustrates deviation from expected frequencies by chance according to log-linear modeling, with darker shading illustrating greater deviation from chance. (E) Plots of the mean transcript abundance over each developmental stage (Tintori et al., 2016) grouped by association with SPN-4, LIN-41 and/or OMA-1 (as in Fig. 2). Unlike Fig. 2, the data here are arranged spatially, then temporally to separate the P-lineage from somatic blastomeres.

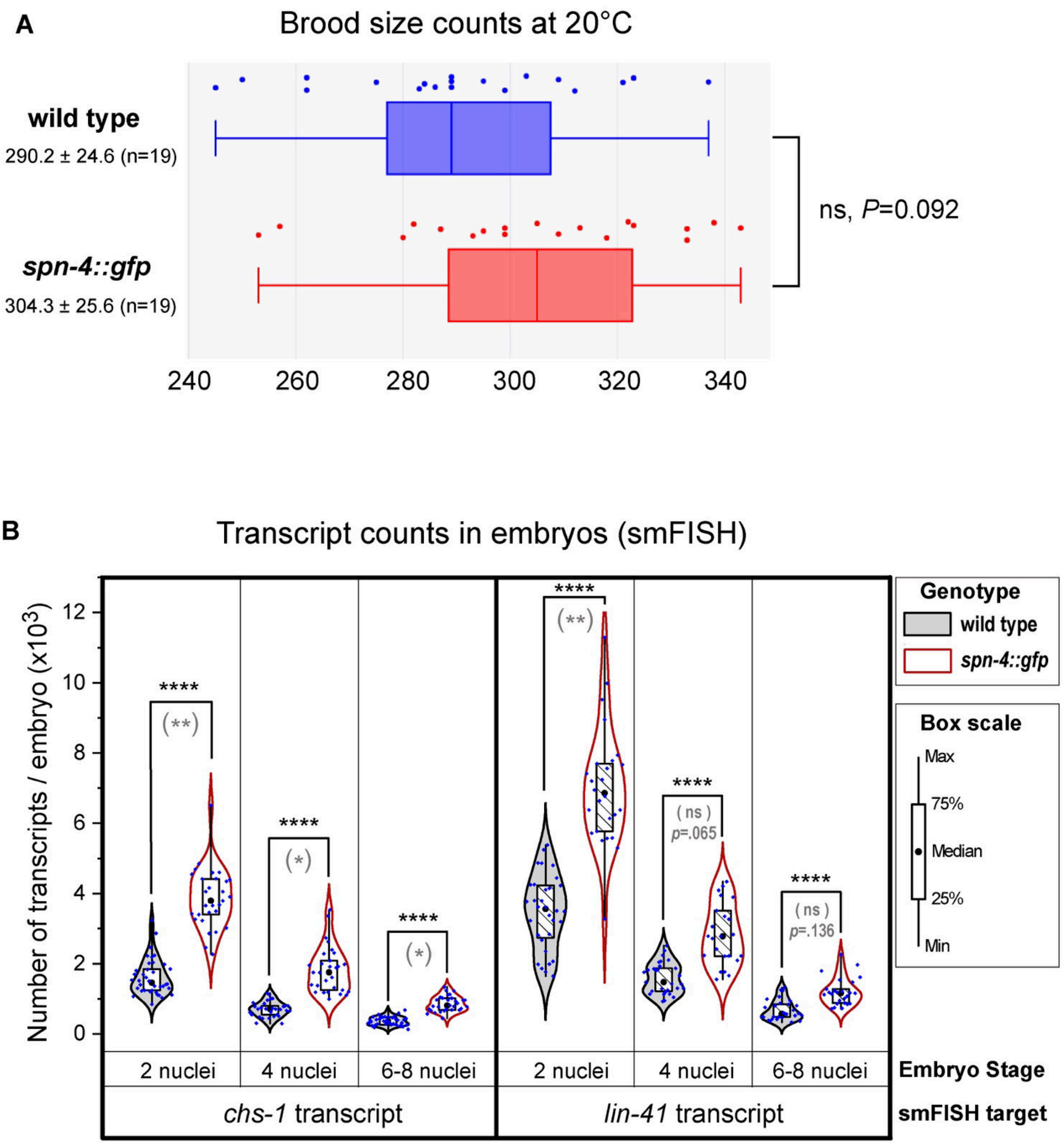


**Figure S4. SPN-4-associated transcripts are more abundant in *spn-4(tn1699[spn-4::gfp])* embryos, consistent with the idea that it is a weak hypomorph.** (Related to Fig. 3). (A) Wild-type and DG4158 *spn-4(tn1699[spn-4::gfp])* animals have brood sizes that are not significantly different (unpaired Student's *t*-test). (B) *lin-41* and *chs-1* transcript numbers are increased in early DG4158 *spn-4(tn1699[spn-4::gfp])* embryos relative to the wild type. The graph shows violin and box plots with individual data points depicting the number of *chs-1* (left side) or *lin-41* (right side) transcripts in individual embryos at early developmental stages (2, 4 or 6-8 nuclei). Wild-type transcript distributions are the same as those shown in Fig. 10 and S13; these data were collectively analyzed for significance at each developmental stage. Black asterisks above a group-connecting bracket indicate the distribution is significantly different from the one observed for stage-matched wild-type embryos using Welch's one-way ANOVA, followed by a Games-Howell post-hoc test to account for unequal variances and multiple comparisons. Because some distributions failed the Shapiro-Wilk test for normality, we also report significance values derived from the non-parametric Kruskal-Wallis test, followed by Dunn's post-hoc test with Benjamini-Hochberg correction for multiple comparisons. Non-parametric significance results are gray and in parentheses. Two or three replicate experiments for each genotype and probe; total n-value >22 at each developmental stage. Significance values: **** $P$<0.0001, *** $P$< 0.001, ** $P$<0.01, * $P$<0.05, ns=not significant. Exact $P$- and n-values are reported in Table S9.

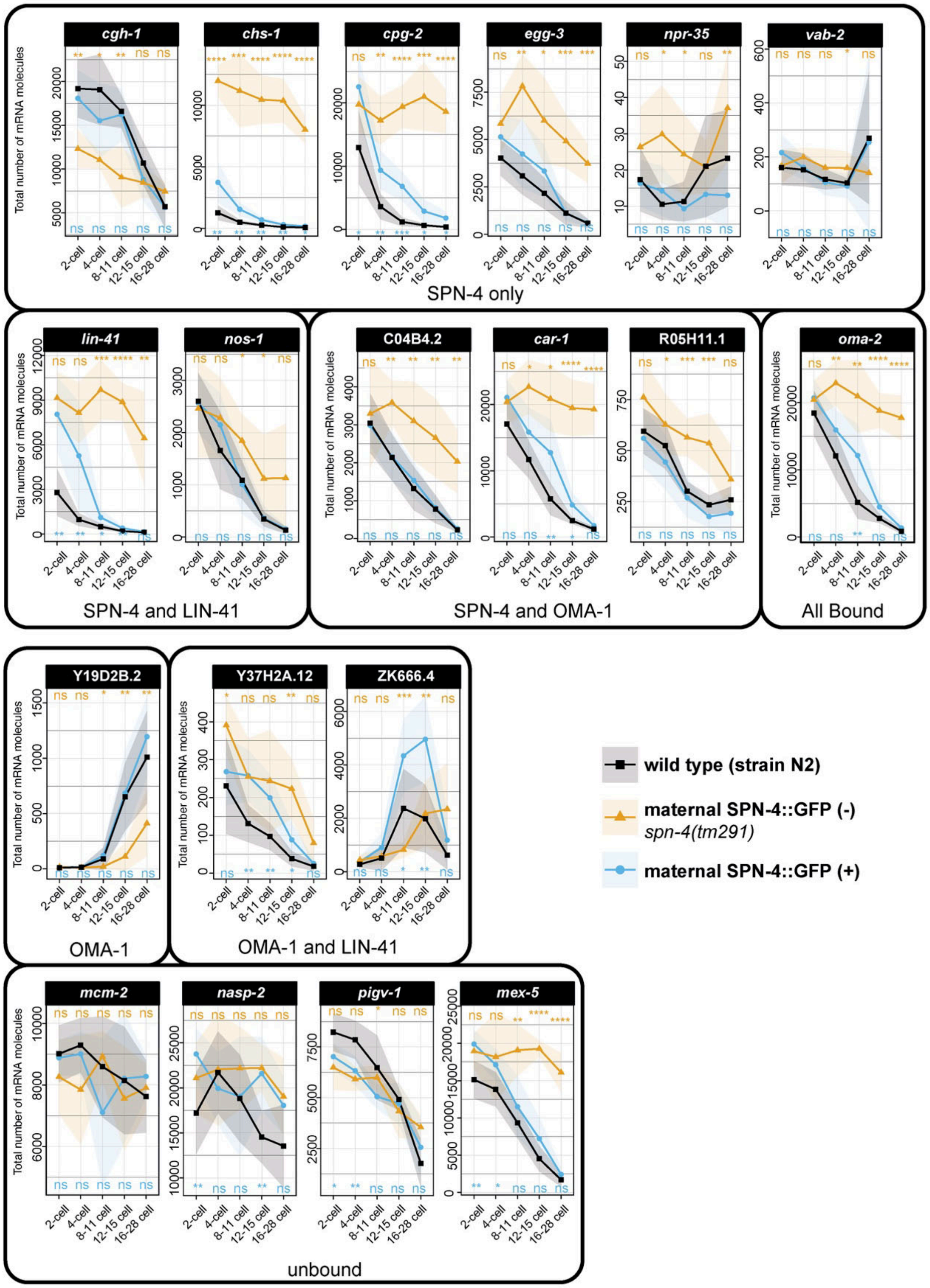


**Fig S5. smFISH plots for select transcripts.** (Related to Fig. 3). The abundance of key transcripts in the wild type (strain N2) or without maternal *spn-4* activity as imaged by smFISH or smiFISH and quantified over five developmental stages. Transcripts are grouped according to their association with SPN-4, LIN-41, and/or OMA-1. Typically, 7 embryos (with a range of 4-12) per transcript, genotype and stage combination were collected across two to three replicates. Ribbons, standard deviation. Statistics = Welch's *t*-tests adjusted by Benjamini-Hochberg multiple test correction. ****: *P* <= 0.0001, ***: *P* <= 0.001, **: *P* <= 0.01, *: *P* <= 0.05, and ns: *P* > 0.05 (not significant).

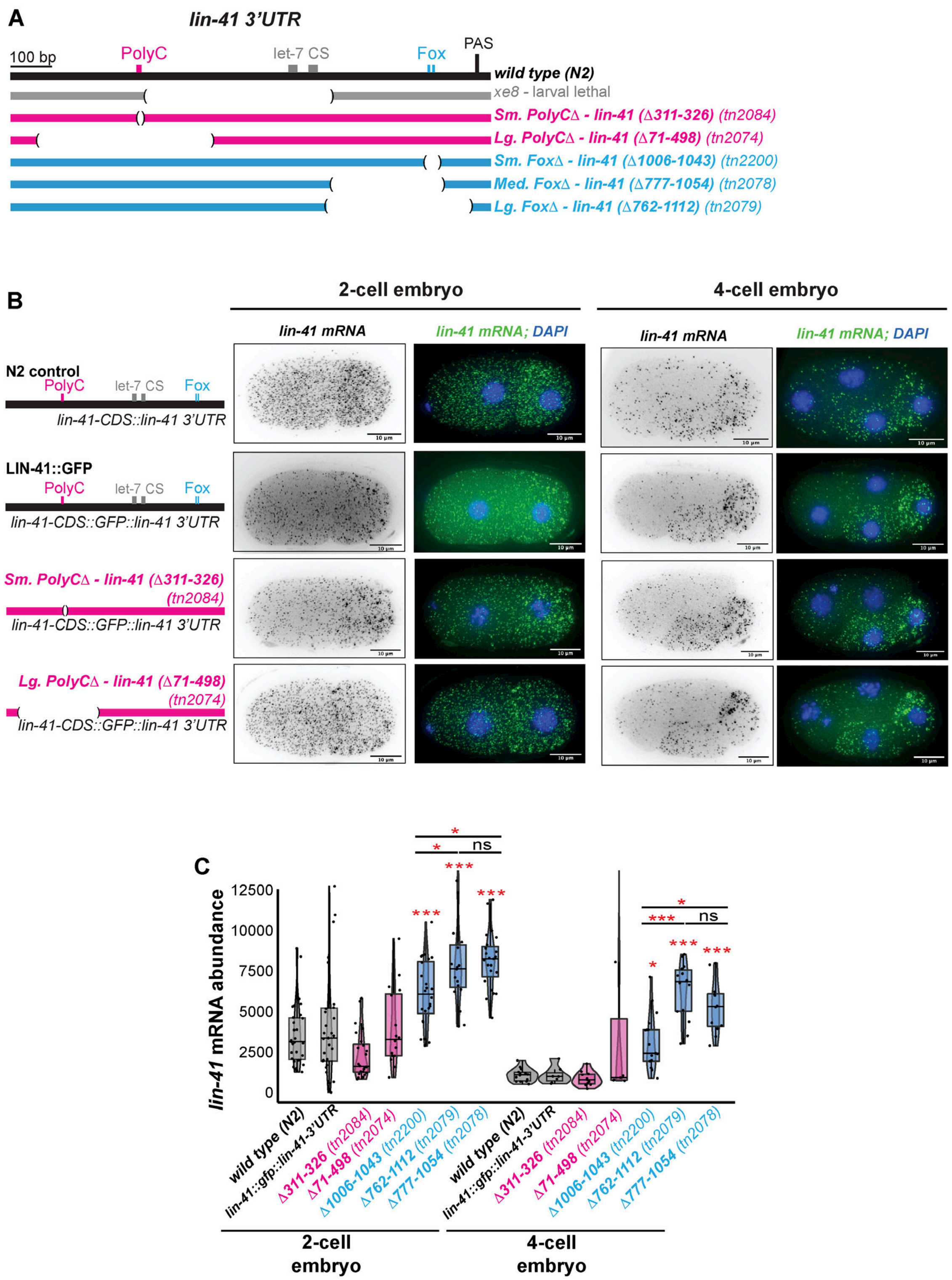


**Fig. S6. The polyC RNA sequence motif in the *lin-41* 3'UTR is not required for maternal mRNA clearance**. (Related to Fig. 4). (A) The PolyC sequence within the *lin-41* 3'UTR was disrupted as either a small (Δ311-326) or large (Δ71-498) deletion in the *lin-41(tn1541[gfp::lin-41])* genetic background. (B) The impact of the polyC-motif deletions on *lin-41* mRNA abundance, compared to the *lin-41(tn1541[gfp::lin-41])* or wild type (N2) control, was assessed by smFISH. Images are representative of 9-31 embryos of each condition combination assayed over 2-4 replicates. Exact *P*- and n-values are reported in Table S6. Bars,10 µm. (C) Quantification, same as in Fig. 4.

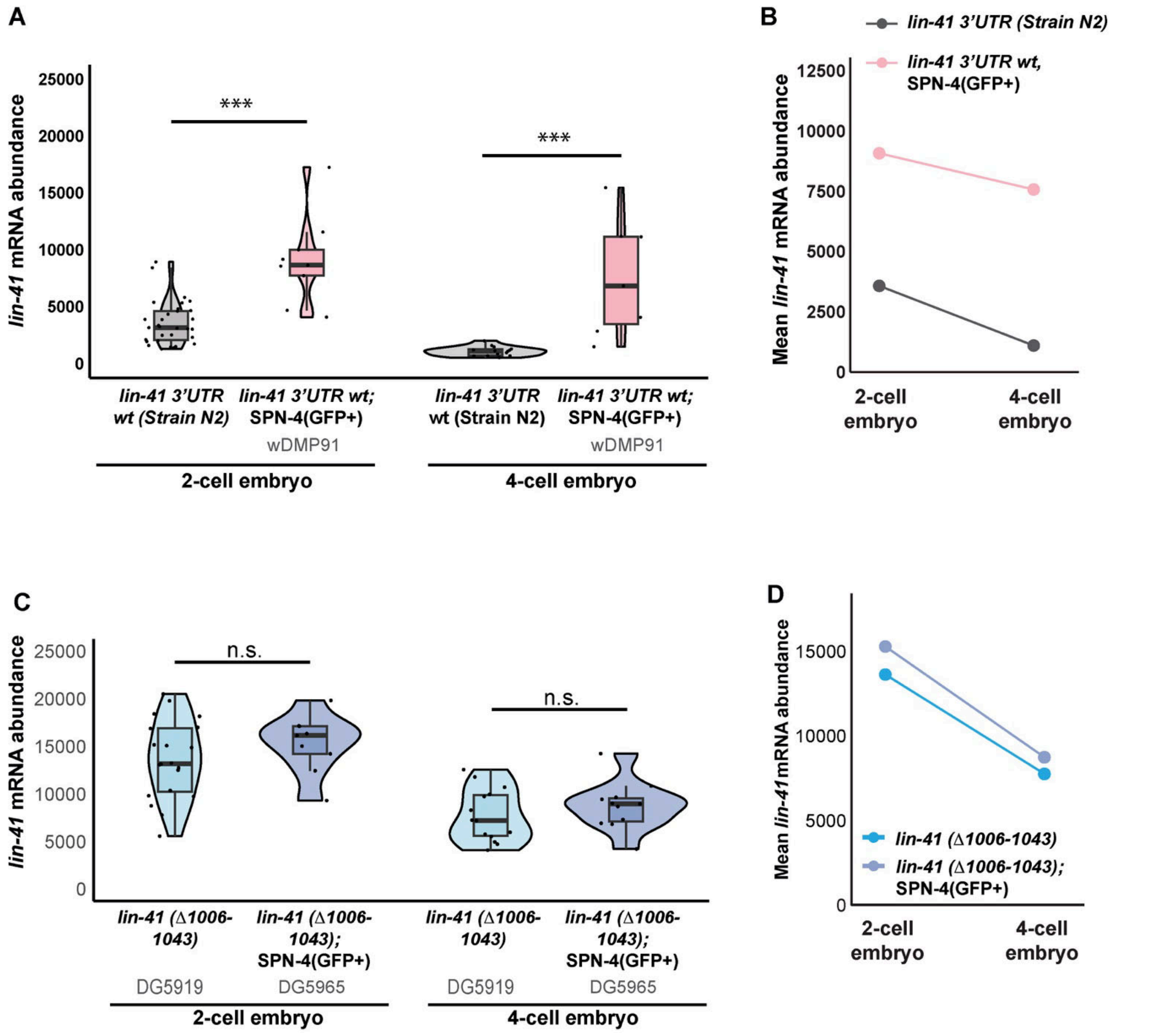


**Fig. S7. SPN-4::GFP, despite being a hypomorph, does not synergize with the *lin-41* Rbfox-motif deletion.** (Related to Fig. 4). (A) Comparison of *lin-41* mRNA levels in embryos produced by the wild type (strain N2) and embryos from strain DG5256 *spn-4(tm291)/tmC3[egl-9(tmIs1230) spn-4(tn2056[spn-4::gfp::3xflag])]* that have a maternal *spn-4::gfp(+)* genotype. Embryos that express SPN-4::GFP exhibit elevated levels of *lin-41* transcripts compared to the wild type. (B) Same dataset as in (A) but with mean *lin-41* mRNA abundance calculated and plotted across the two cell stages. (C) A comparison of *lin-41* mRNA levels from embryos produced by strain DG5919 *lin-41(tn2238*Δ1006-1043*)*, which lacks the Rbfox-motif sequence in its 3'UTR, and embryos produced by strain DG5965 *lin-41(tn2238*Δ1006-1043*); spn-4(tm291)/tmC3[egl-9(tmIs1230) spn-4(tn2056[spn-4::gfp::3xflag])]* that have a maternal *spn-4::gfp(+)* genotype. (D) Same data as (C) but calculating mean *lin-41* mRNA abundance and comparing across cell stages. This result supports the conclusion that SPN-4 functions mainly through the Rbfox motif to regulate *lin-41* maternal mRNA clearance. For all assays: n > 8 for each condition. Statistical tests were performed for each cell stage using anova analysis, and pairwise differences were assessed using the post-hoc Tukey's Honest Significant Differences analysis to account for multiple testing. *** *P*-adj < 0.001; ** *P*-adj < 0.01 ; * *P*-adj < 0.05; ns = not significant.

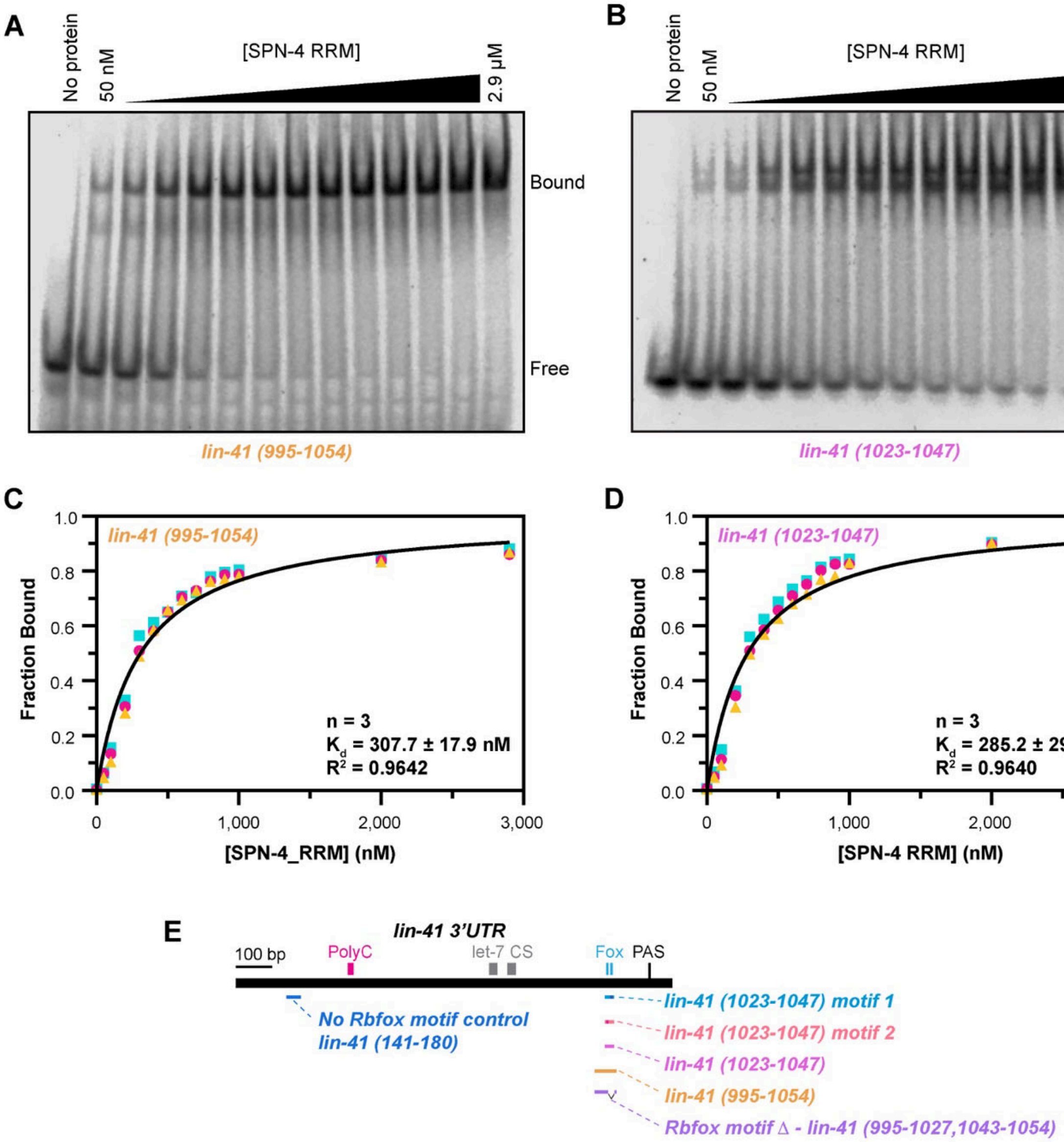


**Fig. S8. *In vitro* binding assays for the 60-nt and 25-nt RNA probes that contain the *lin-41* Rbfox motif.** (Related to Fig. 7). (A,B) Representative *in vitro* binding EMSA assays for the interaction of the SPN-4 RRM with the *lin-41(995-1054)* RNA probe (A) or the *lin-41(1023-1047)* RNA probe (B) run on native 8% polyacrylamide gels. (C,D) Lineplot curves plotting the mean fraction bound over SPN-4 RRM concentration. Calculated $K_d$ values are shown. n=3 replicates. (E) A map of the *lin-41* 3'UTR showing the position of the probes. The sequences of the probes are shown in Fig. 7D. To assess whether the SPN-4 RRM might bind cooperatively, we calculated the Hill coefficient (h) using the equation log(Y/1-Y)=hlog[C]-log$K_d$; where Y is the fraction of bound RNA and [C] is the concentration of the SPN-4 RRM. For binding to the 60-nt *lin-41(995-1054)* RNA probe, h=1.03, indicating that the binding is not cooperative.

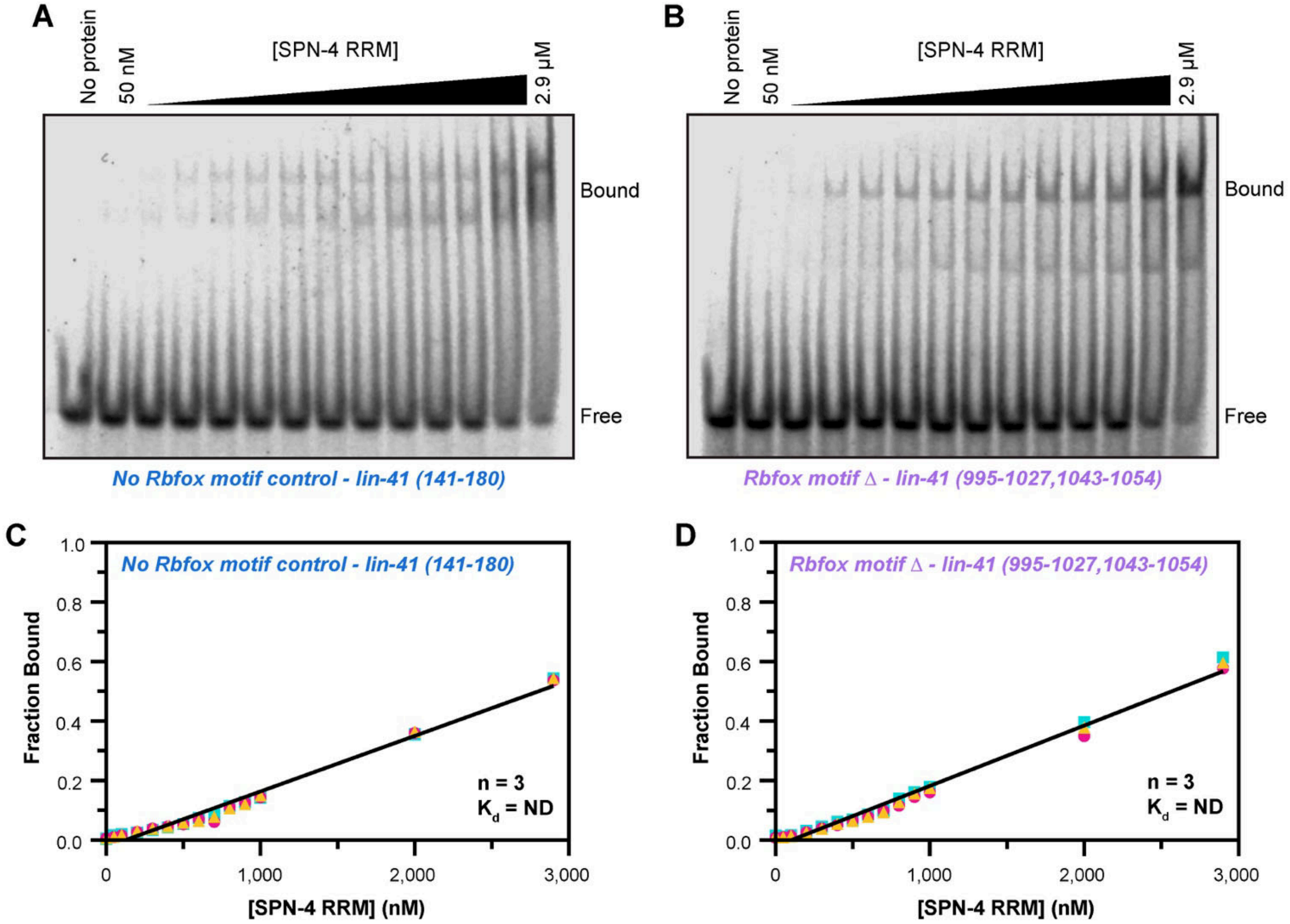


**Fig. S9.** ***In vitro*** **binding assays for the 40-nt and 43-nt RNA probes that do not contain the** ***lin-41*** **Rbfox motif.** (Related to Fig. 7). (A,B) Representative *in vitro* binding EMSA assays for an interaction between the SPN-4 RRM and the *lin-41(141-180)* control RNA probe (A) or the *lin-41(995-1027,1043-1054)* control RNA probe (B) that lacks the Rbfox motif sequence. Assays were run on native 8% polyacrylamide gels. (C,D) Lineplot curves plotting the mean fraction bound over SPN-4 RRM concentration. The SPN-4 RRM exhibits non-specific and non-saturable binding to these control probes. n=3 replicates.

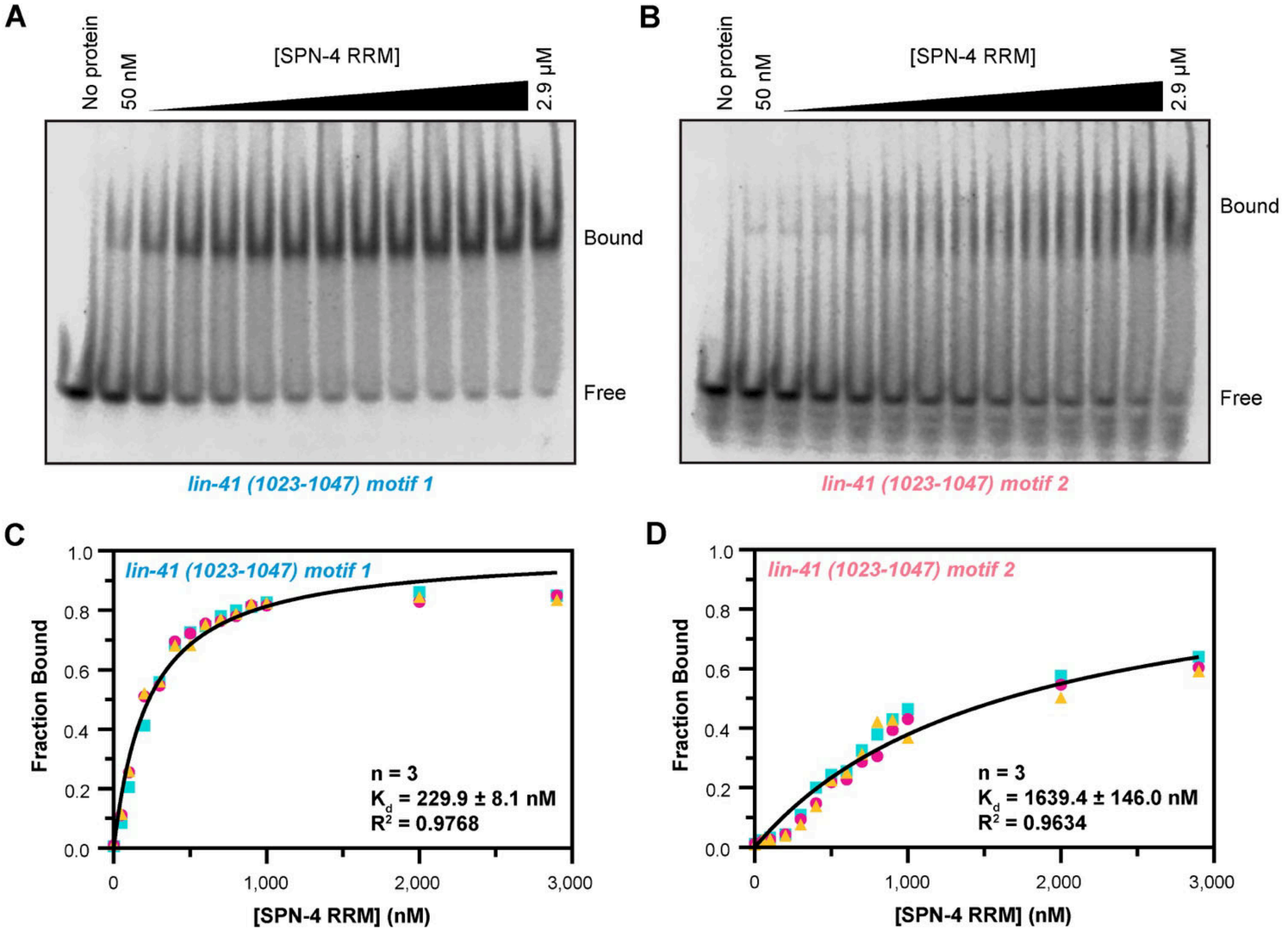


**Fig. S10. SPN-4 preferentially binds RNA containing the Rbfox consensus sequence in motif 1.** (Related to Fig. 7). (A, B) Representative in vitro binding EMSA assays for an interaction between the SPN-4 RRM and the *lin-41(1023-1047)* motif 1-containing RNA probe (A) or the *lin-41(1023-1047)* motif 2-containing RNA probe. Assays were run on native 8% polyacrylamide gels. (C,D) Lineplot curves plotting the mean fraction bound over SPN-4 RRM concentration. Calculated $K_d$ values are shown. n=3 replicates.

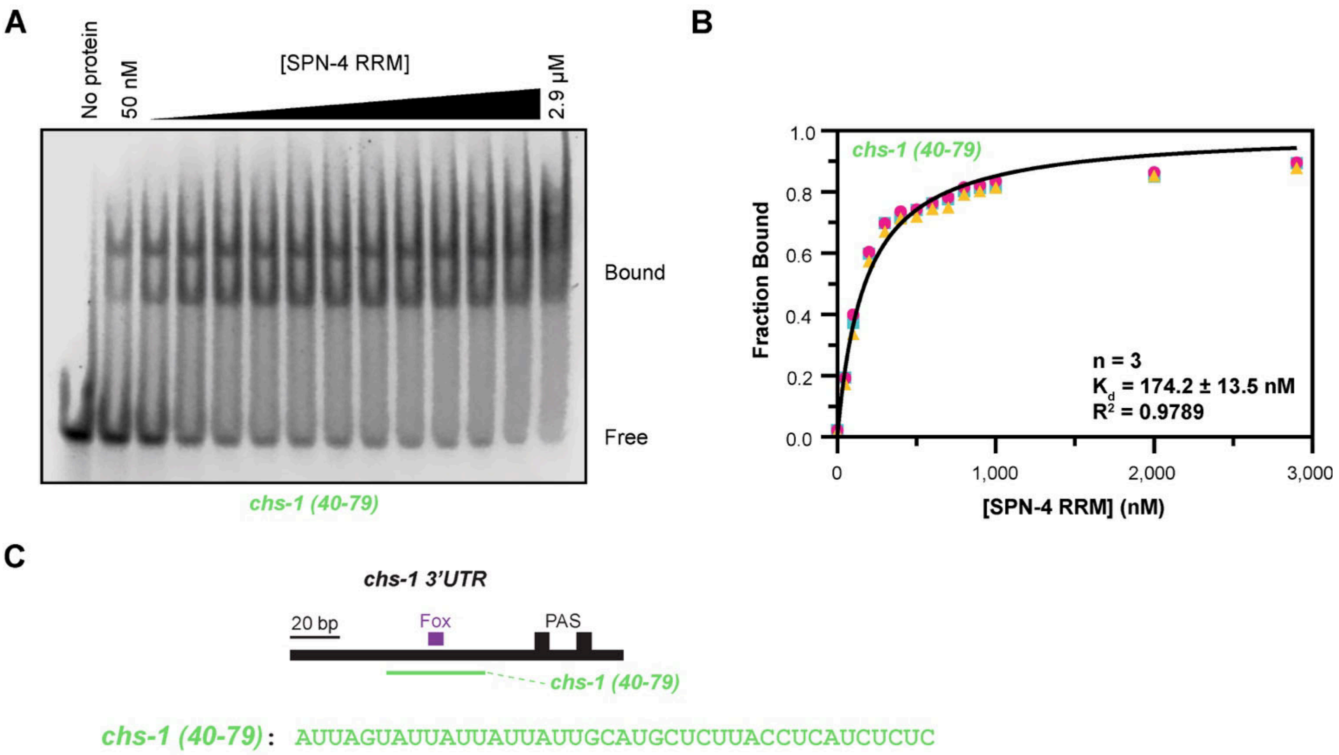


**Fig. S11. The SPN-4 RRM binds the *chs-1* 3'UTR.** (Related to Fig. 7). (A) Representative *in vitro* binding EMSA assay for the interaction of the SPN-4 RRM with the 40-nt *chs-1(40-79)* RNA probe that contains the Rbfox motif. Assays were run on native 8% polyacrylamide gels. (C,D) Lineplot curves plotting the mean fraction bound over SPN-4 RRM concentration. Calculated $K_d$ values are shown. n=3 replicates. (C) Map of the *chs-1* 3'UTR and the sequence of the *chs-1* probe. For binding of the SPN-4 RRM to the *chs-1* RNA probe, we observed a doublet of two bound species. To ascertain whether the SPN-4 RRM might bind cooperatively to the *chs-1* probe, we calculated the Hill coefficient (h) using the equation $\log(Y/1\text{-}Y)=h\log[C]-\log K_d$; where Y is the fraction of bound RNA and [C] is the concentration of the SPN-4 RRM. For binding to the 40-nt *chs-1(40-79)* RNA probe, h=0.79, indicating that the binding is not cooperative. The basis for the formation of a doublet is not known.

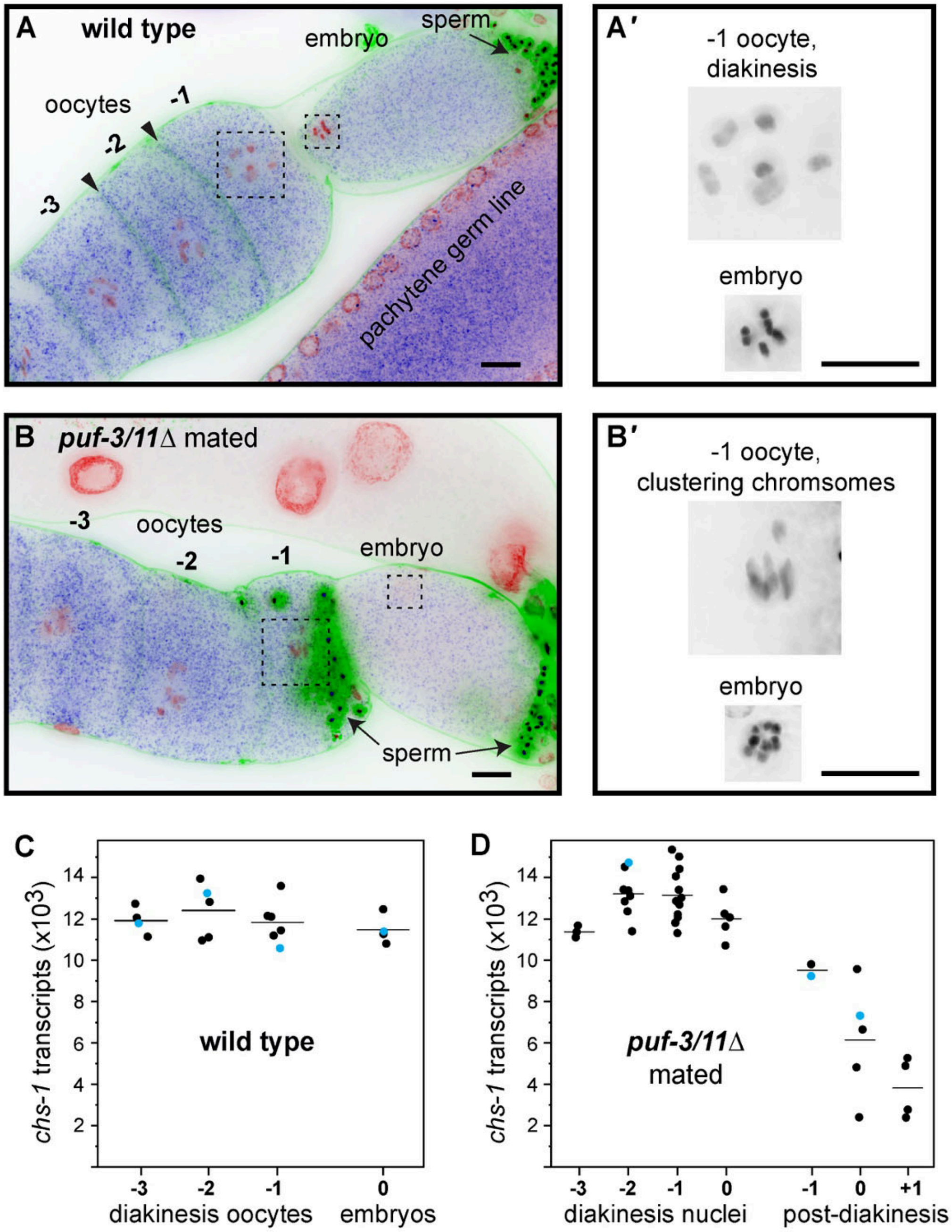


**Fig. S12. *chs-1* transcript counts in wild-type and *puf-3/11*Δ oocytes and newly fertilized embryos.** (Related to Fig. 8). (A) *chs-1* transcripts detected by smFISH (blue) in a wild-type dissected gonad. Oocyte cell boundaries (arrowheads) are identified using Alexa-488-labeled wheat-germ agglutinin (WGA, green), which also robustly stains sperm (arrow). Red shows DNA. The oocytes and the newly fertilized embryo in the spermatheca contain similar levels of *chs-1* transcripts. DNA in boxed areas is magnified 2.5-fold in (A') to illustrate the distinct patterns of chromosome organization in diakinesis oocytes relative to a newly fertilized embryo with clustered chromosomes preparing to undergo the first meiotic division. (B) *chs-1* transcripts detected by smFISH (blue) in the gonad of a *puf-3/11*Δ hermaphrodite permitted to mate with same-genotype males. *puf-3/11*Δ hermaphrodites exhibit low meiotic maturation rates; mating was conducted to match the meiotic maturation rates of wild-type hermaphrodites and observe newly fertilized embryos in the uterus in sufficient quantities. The newly fertilized *puf-*

*3/11*Δ embryo contains fewer *chs-1* transcripts than the oocytes. Image details are the same as for panel (A). Maximum intensity projections of 2.5-micron Z-stacks, rather than the entire Z-stack of images, are shown for better visualization of individual transcripts in these deconvolved images. Note that the chromosomes of the newly fertilized embryo (shown in B') are from a different focal plane. (C,D) Quantification of *chs-1* transcript levels in oocytes and newly fertilized embryos. In the wild type (C) *chs-1* transcript levels remain relatively stable in oocytes and newly fertilized embryos. In contrast, *chs-1* transcript levels decline rapidly after diakinesis in *puf-3/11*Δ embryos (D). Horizontal lines indicate the mean transcript count for each group. Blue dots indicate the number of *chs-1* transcripts in oocytes and newly fertilized embryos in the images shown in panels (A,B). Black dots indicate measurements from other dissected gonads. These data were obtained from 8 wild-type and 17 *puf-3/11*Δ gonad images suitable for counting at least one embryo or oocyte (total images: n=15 wild type, n=25 *puf-3/11*Δ). Few post-diakinesis oocytes were observed in these mated *puf-3/11*Δ hermaphrodites relative to unmated day-1 adult *puf-3/11*Δ hermaphrodites (Fig. 8G,H), likely due to the altered physiological state caused by the presence of additional sperm provided through mating. Nevertheless, these more quantitative data are consistent with the idea that *chs-1* transcript degradation occurs prematurely in *puf-3/11*Δ animals during the oocyte-to-embryo transition. Proximal oocytes (–1 to –3) are indicated; 0 indicates embryos or oocytes in the spermatheca, with +1, indicating the first embryo in the uterus. Bars, 10 μm.

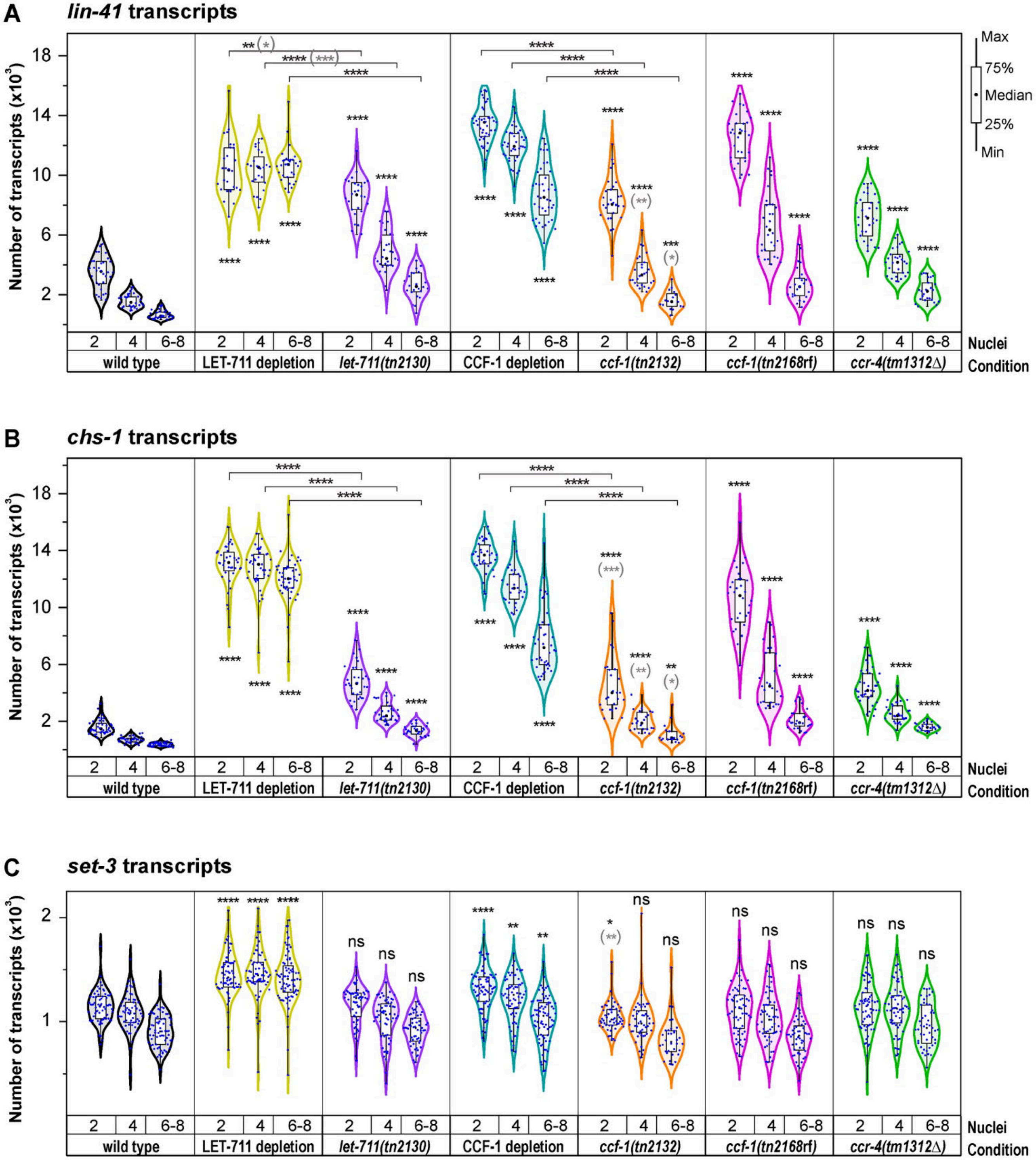


**Fig. S13. *lin-41*, *chs-1* and *set-3* smFISH data related to the CCR-4-NOT complex.** (Related to Fig. 10). (A,B) *lin-41* and *chs-1* transcript numbers are increased in early embryos when the CCR-4-NOT complex subunits LET-711, CCF-1 and CCR-4 are not wild type, with large and more durable effects (up to 32-fold) occurring under the LET-711AID and CCF-1AID protein depletion conditions that cause embryonic lethality. (C) *set-3* transcript numbers are slightly increased by LET-711AID and CCF-1AID protein depletion (up to 1.6-fold). All graphs show violin and box plots with individual data points depicting the number of *lin-*

*41* (A), *chs-1* (B) and *set-3* (C) transcripts in individual embryos at each developmental stage (2, 4 or 6-8 nuclei) and condition (genetic background or auxin-dependent protein depletion). Small black asterisks (not above brackets but located below or above a transcript count distribution) indicate the distribution is significantly different from the one observed for stage-matched wild-type embryos (far left panels in A-C). We also compared the LET-711 AID and CCF-1AID depletions to embryos expressing the same AID-tagged fusion protein under non-depleting conditions (no TIR1 driver or auxin analog). These control embryos had the following genotypes: *let-711(tn2130[let-711::gfp::aid*::3xflag])* or *ccf-1(tn2132[ccf-1::gfp::aid*::3xflag]).* Large black asterisks above a connecting bracket indicate that the transcript counts observed after AID protein depletion are significantly different from those controls lacking the TIR1 driver and auxin analog. Significance values in black were obtained using Welch's one-way ANOVA, followed by a Games-Howell post-hoc test to account for unequal variances and multiple comparisons. Because some distributions failed the Shapiro-Wilk test for normality, we report significance values derived from the non-parametric Kruskal-Wallis test, followed by Dunn's post-hoc test with Benjamini-Hochberg correction for multiple comparisons. Gray asterisks representing these non-parametric significance results are in parentheses; they are only shown when the significance value differs from the Welch's one-way ANOVA result. Significance values: *** *P*<.0001, *** *P*< .001, ** *P*<.01, * *P*<.05, ns=not significant. Two or three replicate experiments for each genotype and two-color probe combination (e.g.: *set-3* + *lin-41* transcript detection). Total *n*-value greater than or equal to 16 at each developmental stage in the different *lin-41* and *chs-1* experiments. Total *n*-value >33 at each developmental stage for *set-3*. Exact *P*- and *n*-values are reported in Table S9.

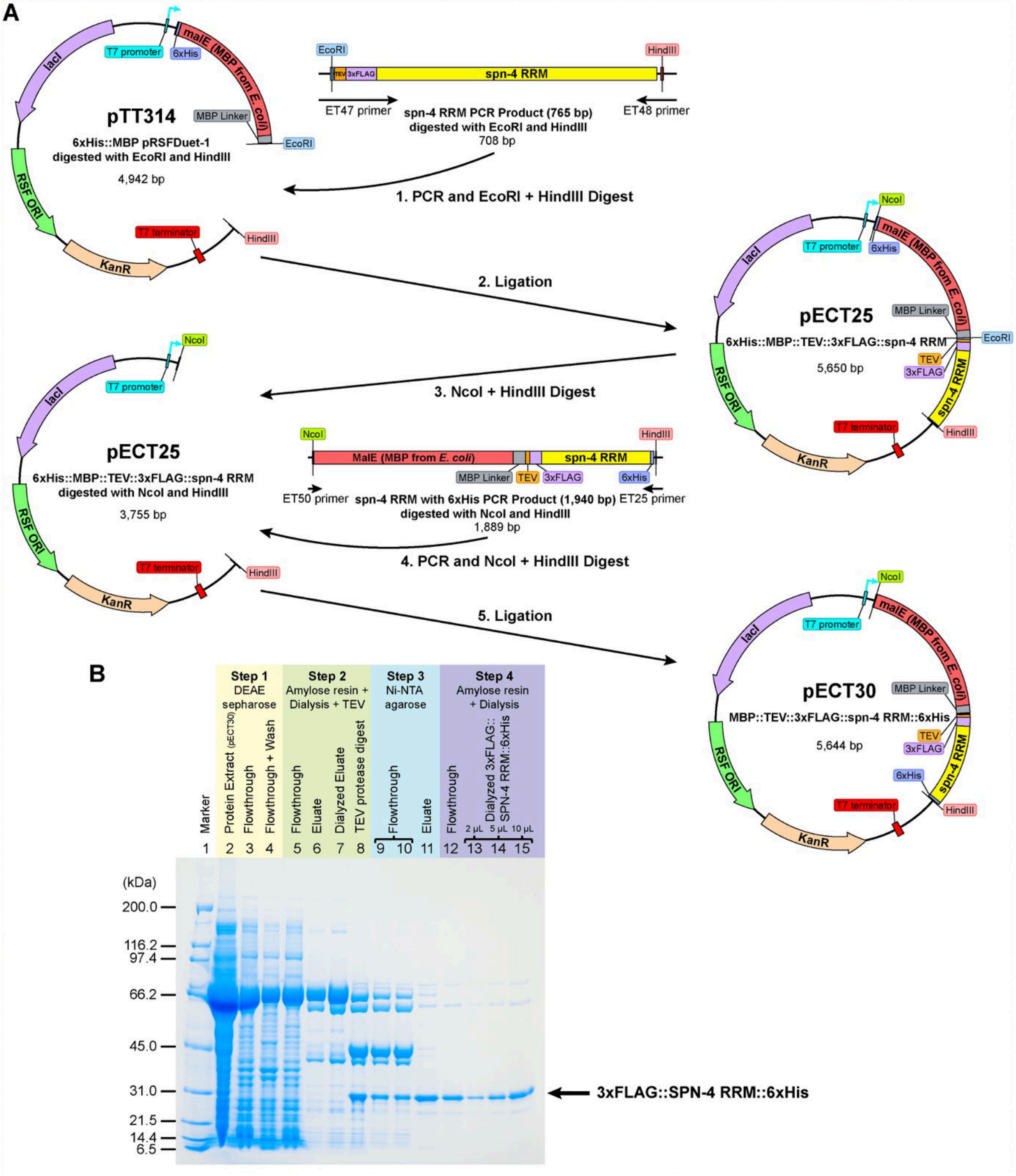


**Fig. S14. Overview of SPN-4 RRM expression plasmid construction and purification.** (Related to Materials and Methods and Fig. 7 and S8-11) (A) Schematic of cloning methods used to generate the SPN-4 RRM expression plasmid, pECT30. (B) A colloidal Coomassie-stained protein 4-12% Bis-Tris NuPAGE gel showing steps used to purify the SPN-4 RRM. All the RNA-binding experiments (Fig. 7 and S8-11) used the SPN-4 RRM shown in lanes 13-15. The gel is representative of two biological replicates.
